# The Non-Phototrophic Hypocotyl3 (NPH3)-domain protein NRL5 is a trafficking-associated GTPase essential for drought resistance

**DOI:** 10.1101/2023.05.10.540297

**Authors:** Neha Upadhyay-Tiwari, Xin-Jie Huang, Yi-Chen Lee, Shashi Kant Singh, Chuan-Chi Hsu, Shih-Shan Huang, Paul E. Verslues

## Abstract

The mechanisms of plant resistance to low water potential (ψ_w_) during drought are unclear but may involve signaling and trafficking at the plasma membrane as well as metabolic reprogramming, including proline accumulation. Forward genetic screening using a *Proline Dehydrogenase 1* (*ProDH1) promoter:reporter* line identified a mutant with extreme low ψ_w_ hypersensitivity due to a single amino acid substitution (P335L) in the Non-Phototrophic Hypocotyl3 (NPH3) domain of NPH3/RPT2-Like5 (NRL5)/Naked Pins in Yucca8 (NPY8). Further experiments found that NRL5, and other NPH3-domain proteins, are GTPases. NRL5 interacted with RAB small GTPases and the SNARE proteins VAMP721/722 and had polar localization. NRL5^P335L^ had greatly reduced GTPase activity, impaired RAB and VAMP721/722 interaction and disrupted polar localization. These data demonstrate that NRL5-mediated restraint of proline catabolism is required for drought resistance and also more broadly define unexpected functions of the NPH3 domain such that the role of NPH3-domain proteins in signaling, trafficking, and cellular polarity can be critically re-evaluated.

**One-Sentence Summary:** A protein containing the plant-specific NPH3-domain has GTPase activity, trafficking interaction and drought resistance function.

## Main Text

Even moderate severity drought stress (moderate reduction of water potential, ψ_w_) leads to reprogramming of cellular processes to retain water while also preparing for more severe water limitation. Thus, plants can detect and initiate signaling in response to small shifts in water status. How this occurs is not known but may include sensing and signaling events at the plasma membrane-cell wall interface (*1-3*). Intracellular trafficking that maintains appropriate composition of the plasma membrane and cell wall, or other cellular compartments involved in stress signaling, may thus also be hypothesized to influence drought sensitivity. However, trafficking regulation in plants, and its relationship to drought response, is unclear and likely to involve both proteins that are functionally conserved compared to their metazoan counterparts as well as plant-specific proteins yet to be identified.

Many plants accumulate free proline during drought. Proline serves as a protective solute to increase cellular osmolarity and precise regulation of the proline cycle of synthesis and catabolism balances redox status (*4, 5*). Stress-induced proline accumulation depends upon increased gene expression and protein level of Δ^1^-Pyrroline-5-Carboxylate Synthetase1 (P5CS1), which catalyzes the first and rate limiting step of proline synthesis, along with concomitant down regulation of the gene encoding Proline Dehydrogenase 1 (*ProDH1*). Proline Dehydrogenase catalyzes the first step of mitochondrial proline catabolism where proline is converted to Δ^1^-pyrroline-5-carboxylate (P5C), the common intermediate of both proline synthesis and catabolism (*4, 6*). ProDH can donate electrons to Coenzyme Q1 (CoQ1) and thus can feed reductant directly into mitochondrial electron transport (*7*). This allows proline to serve as an alternative respiratory substrate (*6, 8*). ProDH can also be a substantial source of ROS production, either by supplying excess reductant to the mitochondrial electron transport chain or by the direct reduction of molecular oxygen to generate reactive oxygen (*8-10*).

The regulation of *ProDH1* expression is complex as it is induced by exogenous proline and induced during drought recovery (re-watering) when high levels of proline need to be catabolized (*4, 6, 8*). However, during drought stress *ProDH1* is down regulated despite high levels of proline accumulation. In this case, drought-related signaling overrides the metabolic signal of high proline (*11, 12*). In contrast, during the hypersensitive response to pathogen infection, *ProDH* up-regulation, along with increased proline synthesis, leads to high levels of mitochondrial ROS and P5C, which promote cell death (*13, 14*). These seemingly opposing purposes of proline metabolism, to sustain cellular activity on the one hand but induce cell death on the other, have been observed in both plants and in metazoans, where proline production and ProDH activity can be a determinant of cancer progression (*3, 15-17*). There is some evidence that plasma membrane-derived signals are required for stress-induced proline accumulation (*18, 19*); however, the mechanisms that regulate and balance the seemingly opposing roles of proline metabolism remain unknown in any organism.

Proline metabolism represents a poorly understood intersection of metabolic and stress signaling mechanisms. To begin to understand the enigmatic mechanisms regulating proline metabolism, we conducted a forward genetic screen using a *ProDH1 promoter:luciferase* (*ProDH1_pro_:LUC*) reporter and identified mutants in which *ProDH1_pro_:LUC* was no longer fully down-regulated during low ψ_w_ stress (*20*). One mutant so identified had a single amino acid change (P335L) in the Non-Phototrophic Hypocotyl (NPH3) domain of NPH3/RPT2-Like5 (NRL5)/Naked Pins in Yucca8 (NPY8). NPH3-domain proteins are a large protein family with more than 30 members in Arabidopsis which typically contain the plant-specific NPH3 domain fused to an N-terminal BTB domain and an unstructured C-terminal extension (*21-23*). The best studied NPH3-domain proteins are NPH3 itself, which interacts with Phot photoreceptors to control their protein stability (*24, 25*), and NPY1-5 (also known as MAB/MEL proteins), which influence auxin signaling by controlling polarity of the PIN auxin transporters (*26-28*). Despite these important signaling roles, the cellular function of the NPH3 domain itself is unknown and most NPH3 domain proteins, including NRL5, remain uncharacterized. Our further study of NRL5 led to the surprising discovery that the NPH3-domain has GTPase activity and interacts with trafficking-related proteins. The NRL5^P335L^ mutation disrupts GTPase activity and trafficking interaction and also alters NRL5 polar localization in leaf and hypocotyl cells. Our results broadly define a new and unexpected function of the NPH3 domain such that the role of NPH3-domain proteins in diverse cellular processes such as auxin signaling, phototropism, and cellular polarity can be critically re-evaluated. These findings will also allow the function of the many other uncharacterized NPH3-domain proteins to more readily be determined. Our data also provide a dramatic demonstration that drought-related signaling, which we now know requires NRL5 to function properly, is needed to restrain proline catabolism and thus allow proline to safely accumulate for drought resistance without inducing damaging ROS and P5C production.

## Results

### An NPH3-domain mutation (NRL5^P335L^) causes drought hypersensitivity because of mis-regulated proline metabolism

Screening of an EMS mutagenized population identified a number of mutants that failed to normally down-regulate *ProDH1_pro_:LUC* after transfer of seedlings to low ψ_w_ stress (*20*). Of these, mutant number 4090 had high expression of *ProDH1_pro_:LUC* in both control and low ψ_w_ stress treatments (Fig. S1A and B) and also hyperaccumulated proline (Fig. S1C). It was also dramatically hypersensitive to moderate severity low ψ_w_ (-0.7 MPa) which slowed, but did not stop, growth of wild type (Fig. 1A and B). We identified a single amino acid substitution (P335L) in NRL5 as the putative causal mutation in 4090 (hereafter referred to as *nrl5-1*, Fig. S1D-F). This was confirmed by the ability of a genomic fragment containing the *NRL5* promoter and coding region (*NRL5_pro_:NRL5-YFP*) to complement both *nrl5-1* (Fig. S1A and B) and a T-DNA knockdown allele (*nrl5-2;* Fig. 1A to C, Fig. S1G). In contrast, *NRL5_pro_:NRL5^P335L^-YFP* failed to complement *nrl5-2* (Fig. 1A to C) despite the NRL5 and NRL5^P335L^ proteins accumulating to similar level (Fig. S1H). These results were consistent with the similar phenotypes of *nrl5-1* and *nrl5-2* (Fig. S1I). *nrl5-2* also had increased DNA damage at low ψ_w_ (Fig. S2) and increased expression of drought-regulated genes (Fig. S3A), indicating that *nrl5* mutants were severely damaged by moderate severity low ψ_w_ which caused minimal damage to wild type.

**Fig. 1:**
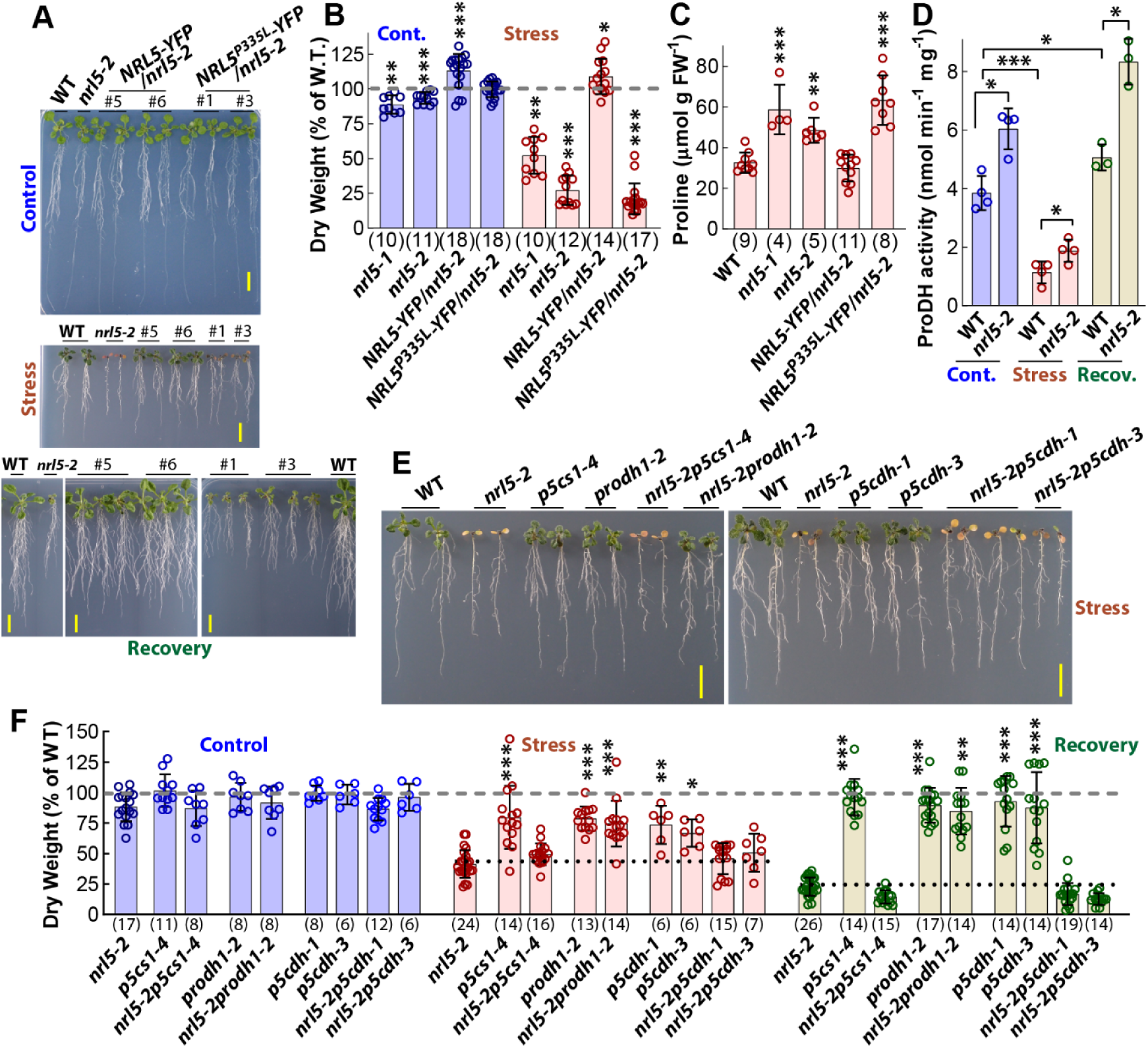
*NRL5* mutants are hypersensitive to low ψ_w_ because of mis-regulated proline metabolism. (**A**) *NRL5* mutants grow normally in unstressed conditions but are hypersensitive to moderate severity low ψ_w_ and have impaired recovery after stress release. Five-day-old seedlings were transferred to fresh control plates and imaged 5 days after transfer or transferred to low ψ_w_ stress (-0.7 MPa) and imaged 10 days after transfer. Stress treated seedlings were transferred back to control plates for four days for recovery. Scale bars indicate 1 cm. Images are representative from more than five independent experiments. Representative images of *nrl5-1* in the same treatments are shown in Fig. S1I. (**B**) NRL5, but not NRL5^P335L^, complemented *nrl5-2* low ψ_w_ hypersensitivity. The transgenic complementation data are combined from 3 independent transgenic lines for each construct. Data were analyzed by one sample Wilcoxon Signed Rank Test for comparison to wild type (WT, 100%), n values are shown in parentheses. Representative images of the complementation lines can be seen in panel A. (**C**) NRL5, but not NRL5^P335L^, complemented the proline hyperaccumulation of *nrl5-2*. Data are from the same transgenic lines assayed in A and B. Data were analyzed by ANOVA with comparison to WT (with P value correction), n values shown in parentheses. (**D**) *nrl5-2* had elevated ProDH enzymatic activity. For stress and recovery, seedlings were transferred to -0.7 MPa for 4 days and then back to control (-0.25 MPa) for one day. Data were analyzed by ANOVA, n = 4 for control and stress treatments, n = 3 for recovery. (**E**) Representative images of seedlings exposed to low ψ_w_ (-0.7 MPa, 10 days) showed that *nrl5-2* low ψ_w_ sensitivity was alleviated in *nrl5-2prodh1-2* but not in double mutants with other enzymes of proline metabolism. Scale bars indicate 1 cm. Seedling images of these genotypes from the control and stress recovery treatments are shown in Fig. S5A. (**F**) Quantitation of seedling dry weight demonstrated that *nrl5* low ψ_w_ sensitivity and inability to recover growth after return to unstressed conditions was alleviated by inactivation of *ProDH1* but not by mutation of *P5CS1* or *P5CDH*. Dry weight was measured five days after transfer of 5-day-old seedlings to unstressed control (-0.25 MPa) or 10 days after transfer to low ψ_w_ stress (-0.7 MPa). Recovery was measured 4 days after transfer of seedlings from the stress treatment back to the high ψ_w_ control. Data were analyzed by Kruskal-Wallis test with comparison to *nrl5-2* in the same treatment, n values are shown in parentheses. For B, C, D, and F, data are from 3 or 4 independent experiments with 2 or 3 biological replicates per experiment (except D which had one replicate per experiment). Error bars show S.D. *, **, and *** indicate P ≤ 0.05, P ≤ 0.01, P ≤ 0.001, respectively.

During low ψ_w_ stress, *nrl5* mutants had increased expression of both *P5CS1*, which encodes the enzyme responsible for the majority of stress-induced proline synthesis, and *P5CS2* (Fig. S3B). Also, both proteomic analysis of *nrl5-2* and immunoblotting found that *nrl5* mutants have increased P5CS1 protein levels during low ψ_w_ stress (Fig. S4A and B; the proteomic results are described more fully below). This was consistent with the higher level of proline accumulation in *nrl5* mutants compared to wild type. Interestingly, *nrl5-2* also had increased ProDH enzymatic activity (Fig. 1D), even though ProDH1 protein levels were not changed (Fig. S4A and C). This pattern indicated that proline synthesis and catabolism were elevated in *nrl5* mutants, allowing proline to accumulate to high levels even though it was still being actively catabolized.

Generation of double mutants between *nrl5-2* and the proline metabolism mutants *p5cs1-4*, *prodh1-2* and *p5cdh* (the latter of which lacks P5C Dehydrogenase, the enzyme which converts P5C to glutamate in mitochondria) showed that blocking proline catabolism upstream of P5C production (*nrl5-2prodh1-2*) alleviated the low ψ_w_ sensitivity and slow recovery of *nrl5-2* (Fig. 1E and F, Fig. S5A). In contrast, blocking proline catabolism downstream of P5C production (*nrl5-2p5cdh-1, nrl5-2p5cdh-3*) did not alleviate the *nrl5-2* phenotype. Partially blocking proline synthesis using *p5cs1-4* was also not sufficient to alleviate the *nrl5-2* phenotype (Fig. 1E and F, Fig. S5A).

In this set of mutants, low ψ_w_ hypersensitivity did not correlate with the level of low ψ_w_-induced proline accumulation (Fig. S5B). Instead, hypersensitivity to low ψ_w_ was corelated with elevated levels of compounds that react with o-aminobenzaldehyde (o-AB), which includes P5C (Fig. S5C), and also corelated with elevated ROS (Fig. S6). The levels of o-AB-reactive compounds and ROS were elevated in *nrl5-2*, *nrl5-2p5cs1-4* and *nrl5-2p5cdh* mutants but reduced to wild type level in *nrl5-2prodh1-2.* This was consistent with *nrl5-2prodh1-2* having nearly wild type level of growth in the low ψ_w_ and recovery treatments (Fig. 1E and F, Fig. S5A). Note that the background levels of o-AB reactive compounds in wild type and mutants may include metabolites other than P5C; however, the increased level seen in *nrl5, nrl5-2p5cs1-4* and *nrl5-2p5cdh* mutants which was eliminated in *nrl5-2prodh1-2,* can be more confidently attributed to P5C (Fig. S5C). Thus, the combined genetic and physiological data indicate that *nrl5* mutants were hypersensitive to low ψ_w_ because of an inability to restrain proline catabolism and thereby limit P5C and ROS production. This occurred despite *nrl5* mutants accumulating high levels of proline which would often be associated with increased drought resistance (*3*).

### NRL5, but not NRL5^P335L^, interacts with RABE1c, RABH1b and VAMP721/722

The NRL5^P335L^ substitution is in its NPH3 domain, a plant specific protein domain of unknown function (*22*). To investigate why NRL5^P335L^ was non-functional and unable to complement *nrl5-2*, we immunoprecipitated NRL5-YFP and identified RAB small GTPases and VAMP721 as putative NRL5 interactors (Data S1). RAB small GTPases are highly conserved components of intracellular trafficking. When RABs are in their active, GTP-bound, state they facilitate vesicle trafficking, including targeting of vesicles to specific membranes (*29*). After GTP hydrolysis, inactive GDP-bound RABs are typically recycled to a source membrane where they can be reactivated by GTP binding. GST-pull down assays demonstrated that NRL5 interacted with RABE1c and RABH1b in the presence of GTPγS, a nonhydrolyzable GTP analog which keeps RABs in their active state, but had reduced interaction in the presence of GDP (Fig. 2A; input shown in Fig. S7A). In contrast, NRL5^P335L^ did not interact with RABE1c or RABH1b even in the presence of GTPγS (Fig. 2A). We did not detect NRL5 interaction with RABA4a (Fig. S7B), indicating specificity of NRL5 interaction with certain RAB subgroups.

**Fig. 2:**
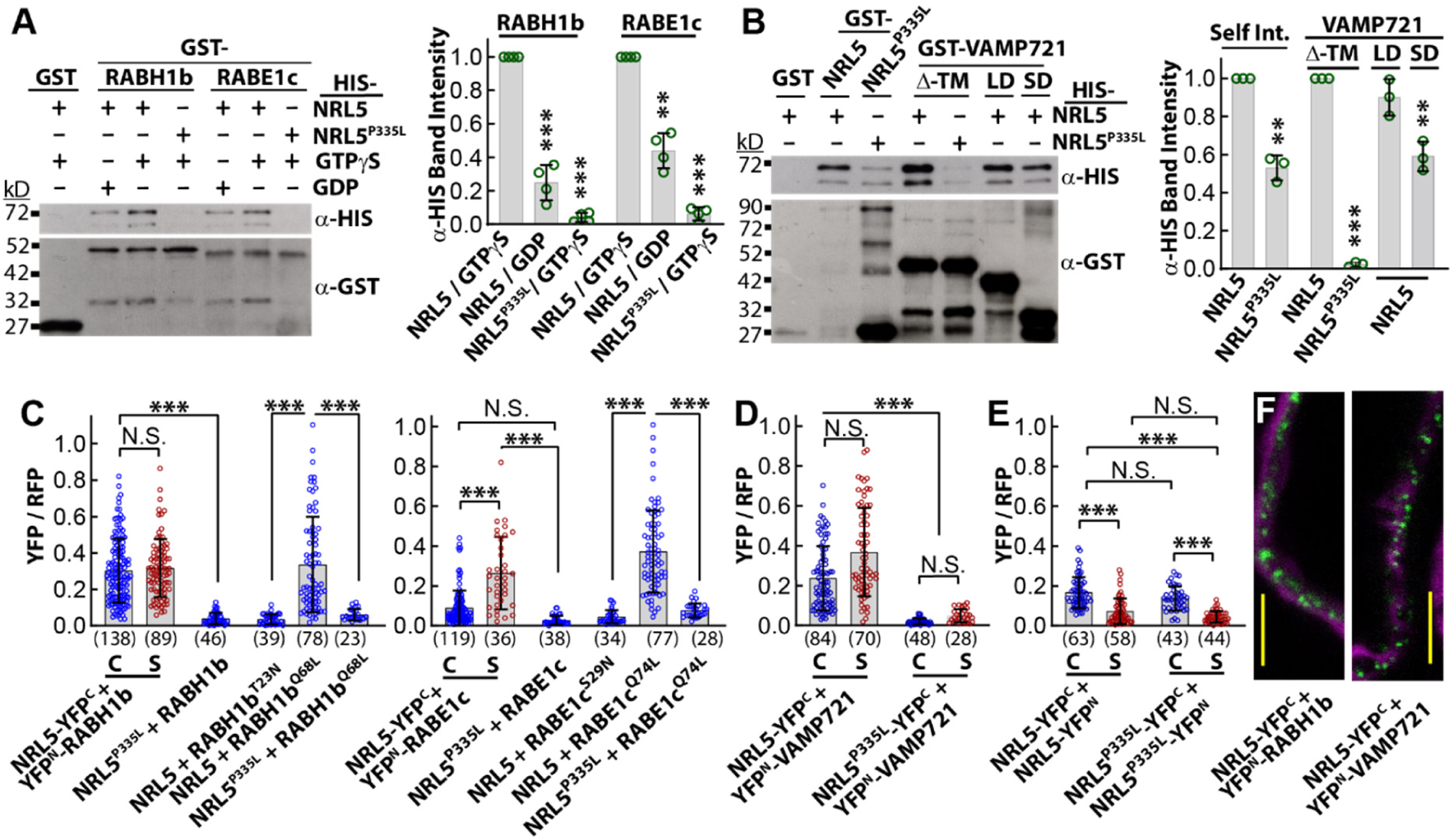
NRL5 interacts with RABH1b, RABE1c and VAMP721/722. (**A**) In vitro pull-down assay demonstrated that NRL5, but not NRL5^P335L^ (both with 6x-HIS tag), interacted with RABH1b and RABE1c in the presence of GTPγS, but had reduced interaction in the presence of GDP. Blots showing equal input of each protein are shown in Fig. S7A. Band intensity quantitative data was analyzed by one sample T-test, n = 4, error bars show S.D. Molecular weight of respective proteins: GST-RABH1b and GST-RABE1c = 50 kD, HIS-NRL5/ NRL5^P335^ = 71 kD, GST= 27 kD. (**B**) NRL5, but not NRL5^P335L^, interacted with VAMP721. But, NRL5^P335L^ could still interact with itself. Data analysis was the same as described for A but with n = 3. Input blots and quantitation are shown in Fig. S7C. A similar pattern of interaction was observed for VAMP722 (Fig. S7D). Additional assays demonstrated that NRL5 self-interaction required the BTB-domain but was not substantially affected by the P335L mutation (Fig. S7E). Molecular weight of respective proteins: HIS-NRL5/NRL5^P335L^ = 71 kD, GST-NRL5 = 98 kD, GST-VAMP721^Δ-TM^ = 47 kD, GST-VAMP721-Longin Domain (LD) = 40 kD, GST-VAMP721-Snare Domain (SD) = 33 kD, GST= 27 kD. (**C**) Quantitation of rBiFC fluorescence intensity ratio showed that NRL5 interacted with wild type or constitutively active RAB (RABH1b^Q68L^, RABE1c^Q74L^) but had greatly reduced interaction with inactive RAB (RABH1b^T23N^, RABE1c^S29N^). NRL5^P335L^ had minimal or no interaction with either wild type or constitutive active RABH1b or RABE1c. Data points are YFP/RFP ratios of individual cells and are combined from three independent experiments. Error bars indicate S.D., n values are shown in parentheses. Data were analyzed by Kruskal-Wallis test. C = Control, S = Stress (-0.7 MPa). (**D**) rBiFC shows interaction of NRL5 and VAMP721. The rBiFC signal was increased by low ψ_w_ for NRL5 but reduced to background level for NRL5^P335L^. Data analysis and presentation are as described for C. (**E**) A similar level of rBiFC signal was detected for self-interaction of NRL5 or NRL5^P335L^ indicating that the P335L mutation does not interfere with NRL5 self-interaction. The rBiFC signal was reduced by low ψ_w_ treatment in both cases. (**F**) Representative rBiFC images for NRL5 interaction with RABH1b or VAMP721 in plants exposed to low ψ_w_ stress (Green = rBiFC signal; magenta = propidium iodide staining of cell wall). Scale bars indicate 10 μm. Additional rBiFC images can be seen in Fig. S8. **, *** and N.S. indicate P ≤ 0.005, P ≤ 0.001 and non-significant differences, respectively.

VAMP721, and its close homolog VAMP722, are R-SNAREs involved in vesicle targeting and docking at the plasma membrane (*30, 31*). NRL5, but not NRL5^P335L^, interacted with VAMP721 (Fig. 2B; input Fig. S7C) and VAMP722 (Fig. S7D). NRL5 interacted with both the Longin and SNARE domains of VAMP721 (Fig. 2B). In contrast to the RAB and VAMP interactions, NRL5 self-interaction (likely dimerization), which occurs via the BTB domain, was not blocked by the P335L mutation (Fig. 2B and Fig S7E).

Interestingly, further pull-down assays found that RABE1c could also interact with VAMP721 (Fig. S7F). In contrast to the NRL5-RABE1c interaction, and many other RAB interactions, the RABE1c-VAMP721 interaction was detected regardless of whether RABE1c was in its GTPγS-bound form or inactive GDP-bound form (Fig. S7F). We also confirmed that the NRL5-VAMP721 interaction was not affected by the presence of GTPγS (Fig. S7G). These results raise the possibility that NRL5, VAMP721/722 and certain RABs may compete for interaction with each other, with the results depending, in part, on the RAB activation state.

Ratiometric Bi-molecular fluorescence complementation (rBiFC) assays detected NRL5 interaction with both wild type and constitutively active versions of RABE1c and RABH1b (RABE1c^Q74L^ and RABH1b^Q68L^); but, not with constitutively inactive RABE1c^S29N^ or RABH1b^T23N^ (Fig. 2C, representative images shown in Fig. S8). In contrast, no interaction was detected for NRL5^P335L^ with either wild type or the constitutive active version of either RAB. Likewise, VAMP721 interacted with NRL5, but not NRL5^P335L^ (Fig. 2D and Fig. S8). The rBiFC assays also confirmed that self-interaction of NRL5 was not affected by the P335L substitution (Fig. 2E and Fig. S8A and E). For all detectable interactions, the rBiFC signal was observed as foci close to the periphery of the cell (Fig. 2F and Fig. S8B). Given the well-known functions of RABs and VAMP721/722 in vesicle trafficking, this pattern suggested that the NRL5 is also associated with vesicle trafficking. We also noted that, NRL5 self-interaction was significantly decreased by low ψ_w_ while NRL5 interaction with RABs and VAMP721 was increased or unchanged (Fig. 2C-E).

As a further control for the specificity of rBiFC assays, we observed that RPT2, which is a member of a different branch of the NPH3-domain family (*22*), had minimal or no detectable interaction with RABE1c, RABH1b or VAMP721 (Fig. S9). However, RPT2 did self-interact with a relative fluorescence intensity similar to that of NRL5 (compare Fig. S9A to Fig. 2E and Fig. S8A).

### NRL5, and other NPH3-domain proteins, have GTPase activity

The switching of RABs between their inactive versus active states is regulated by effector proteins that control the rate of GTP hydrolysis (*29*). In the course of testing whether NRL5 might be such an effector protein that influences RAB GTPase activity, we made the surprising discovery that NRL5 is itself a GTPase. Highly purified NRL5 generated by *in vitro* translation catalyzed P_i_ release with a similar rate and half-maximal GTP concentration as the GTPase activity of RABE1c, RABH1b, or RABA4a (Fig. 3A to C, Fig. S10A and B). In contrast, NRL5^P335L^ had greatly reduced, but not completely abolished, activity (Fig. 3A and B, Fig. S10A). No activity was observed when ATP or GTPγS were used as substrates (Fig. 3A). Similar results were observed for NRL5 purified from *E. coli* (Fig. S10A and B).

**Fig. 3.**
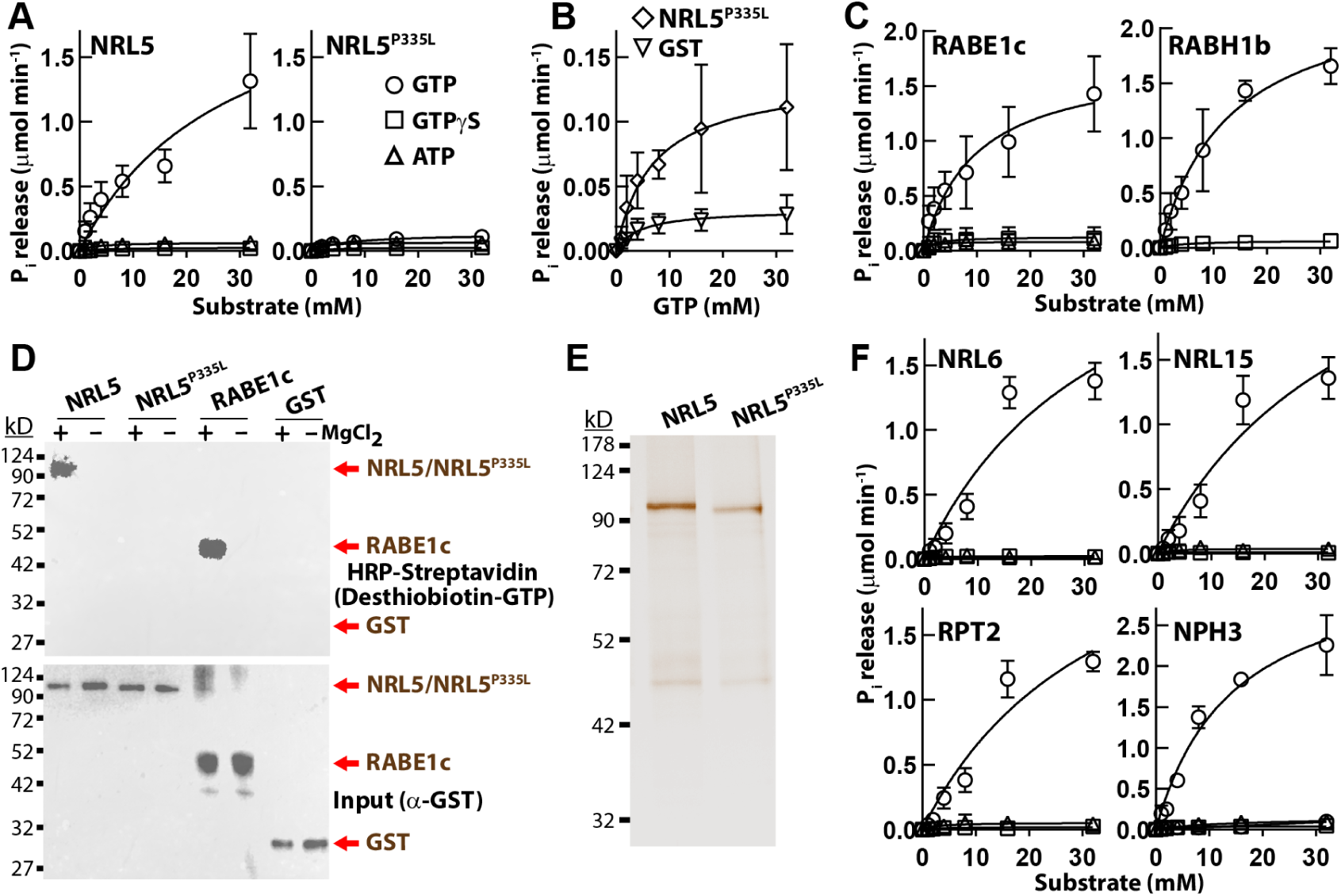
NRL5 associates with GTP and has GTPase activity. (**A**) NRL5 had GTPase activity while activity of NRL5^P335L^ was greatly reduced. Both NRL5 and NRL5^P335L^ were produced by *in vitro* translation. One μM protein was used in each assay with the indicated substrate concentrations and P_i_ release quantified after 30 min (reaction volume was 20 μl). Data were analyzed by non-linear regression (K_cat_, K_m_, and V_max_ values are shown in Fig. S10A). Data are mean ± S.D. (n = 4 for GTP, n = 3 for GTPγS and ATP). Additional kinetic parameters are given in Fig. S10A and GTPase activity for NRL5 and NRL5^P335L^ purified from *E. coli* is shown in Fig. S10B. (**B**) NRL5^P335L^ has a low residual level of GTP hydrolysis activity. The GTP hydrolysis data of NRL5^P335L^ was replotted along with data for GST control protein (produced by in vitro translation and purified in the same manner as NRL5 and NRL5^P335L^) Data are means ± S.D. (n = 4). Additional data of GST control protein purified from *E. coli* is given in Fig. S10. (**C**) RABH1b and RABE1c purified from *E. coli* have similar GTP hydrolysis kinetics as NRL5. Data are mean ± S.D. (n = 6 for RABE1c GTP hydrolysis, n = 3 for all other data). Additional kinetic parameters and data for RABA4a can be found in Fig. S10. (**D**) Desthiobiotin-GTP labeling confirms that NRL5 associates with GTP while NRL5^P335L^ has greatly reduced GTP association (NRL5/NRL5^P335L^pT7CFE1-NHIS-GST-CHA = 105 kD). Proteins (0.5 μM) were incubated with 20 μM Desthiobiotin-GTP in a 25 μl reaction volume for 10 minutes with or without MgCl_2_. RABE1c was used as a positive control (GST-RABE1c = 50 kD) and GST as a negative control (pT7CFE1-NHIS-GST-CHA = 30 kD). The anti-GST blot (5 % of input) shows equal input amount for each Desthiobiotin-GTP labeling reaction. The experiment was repeated with consistent results. A representative example of several similar experiments conducted with NRL5 and NRL5^P335L^ purified from *E. coli* can be seen in Fig. S10D. (**E**) Silver stained gel demonstrates the purity of NRL5 and NRL5^P335L^ generated by in vitro translation. The expected molecular weight of the NRL5/NRL5^P335L^pT7CFE1-NHIS-GST-CHA produced is 105 kD. (**F**) GTPase activity of additional NPH3 domain proteins produced in *E. coli*. Kinetic data are shown in Fig. S10A and gels showing purity of proteins purified from *E. coli* are shown in Fig. S10E. Data are means ± S.D. (n = 6).

Use of a desthiobiotin-GTP probe, which biotinylates GTP-binding proteins (*32*), confirmed that NRL5 associated with GTP and this association required Mg^2+^ as a cofactor (Fig. 3D). This was similar to other GTP-binding proteins, including RABE1c which was used as a positive control. Note that the protein band labeled by desthiobiotin-GTP matched the molecular weight of purified NRL5 analyzed by silver-stained SDS-PAGE gel (Fig. 3E). In both cases, these bands matched the expected molecular weight of the GST-NRL5 fusion produced by in-vitro translation (Fig. 3E). The GTP-association was greatly reduced for NRL5^P335L^ (Fig. 3D). Similar results were obtained for NRL5 and NRL5^P335L^ produced in *E. coli* (Fig. S10D).

Combining desthiobiotin-GTP labeling with LC-MS analysis found biotinylation of lysine residues flanking the P335 site critical for GTPase activity (Fig. S11A, Data S2). Albeit, that there was also labeling of the unstructured C-terminus as well as the N-terminal portion of the BTB domain, which is not required for GTPase activity (see below). Sequence alignment and structural prediction showed that P335 is not conserved among NPH3-domain proteins and is thus unlikely to participate directly in GTP binding or hydrolysis. Rather it is in a linker region between conserved helices (Fig. S11B to D). Thus, the P335L mutation could affect the spatial arrangement of these conserved helices. This could explain how the P335L mutation greatly reduced, but did not totally eliminate, NRL5 GTPase activity (Fig. 3B). It can also explain how the P335L mutation impaired not only GTPase activity and desthiobiotin-GTP labeling (Fig. 3D), but also VAMP721/722 interaction which does not require GTP (Fig. 2B).

We also observed GTPase activity for three additional NPH3-domain proteins, NRL6, NRL15/NPY7 and Root Phototropism2 (RPT2), selected from several branches of the NPH3 protein family (Fig. 3F and Fig. S10A; see Christie et al. (*22*) and Pedmale et al. (*23*) for phylogenetic trees of Arabidopsis NPH3-domain proteins). In addition, NPH3 itself, the best characterized NPH3-domain protein, had similar GTPase activity (Fig. 3F). These results indicated that GTPase activity is a common feature of NPH3-domain proteins.

### Cross-interactions of NRL5, RABE1c, RABH1b and VAMP721/722 affect GTPase activity

The observation that NRL5 is itself a GTPase did not eliminate the possibility that it could also act as an effector that controls RAB GTPase activity (or, vice versa RAB interaction may affect NRL5 activity). Interestingly, combining NRL5 with RABE1c or RABH1b resulted in nearly complete elimination of GTPase activity, indicating that the proteins inhibit each other (Fig. 4A). In contrast, combining NRL5 with RABA4a did not lead to the same inhibition of GTPase activity (Fig. 4A), consistent with the lack of detectable interaction in GST-pulldown assays (Fig. S7B). This indicated that inhibition of GTPase activity required a specific NRL5-RAB interaction. Truncated NRL5 lacking the BTB domain still had GTPase activity and had similar ability to inhibit P_i_ release when combined with RABH1b or RABE1c (Fig. 4B). In contrast, NRL5^P335L^ failed to inhibit RABH1b or RABE1c GTPase activity (Fig. 4B), consistent with its greatly reduced RAB interaction (Fig. 2A).

**Fig. 4.**
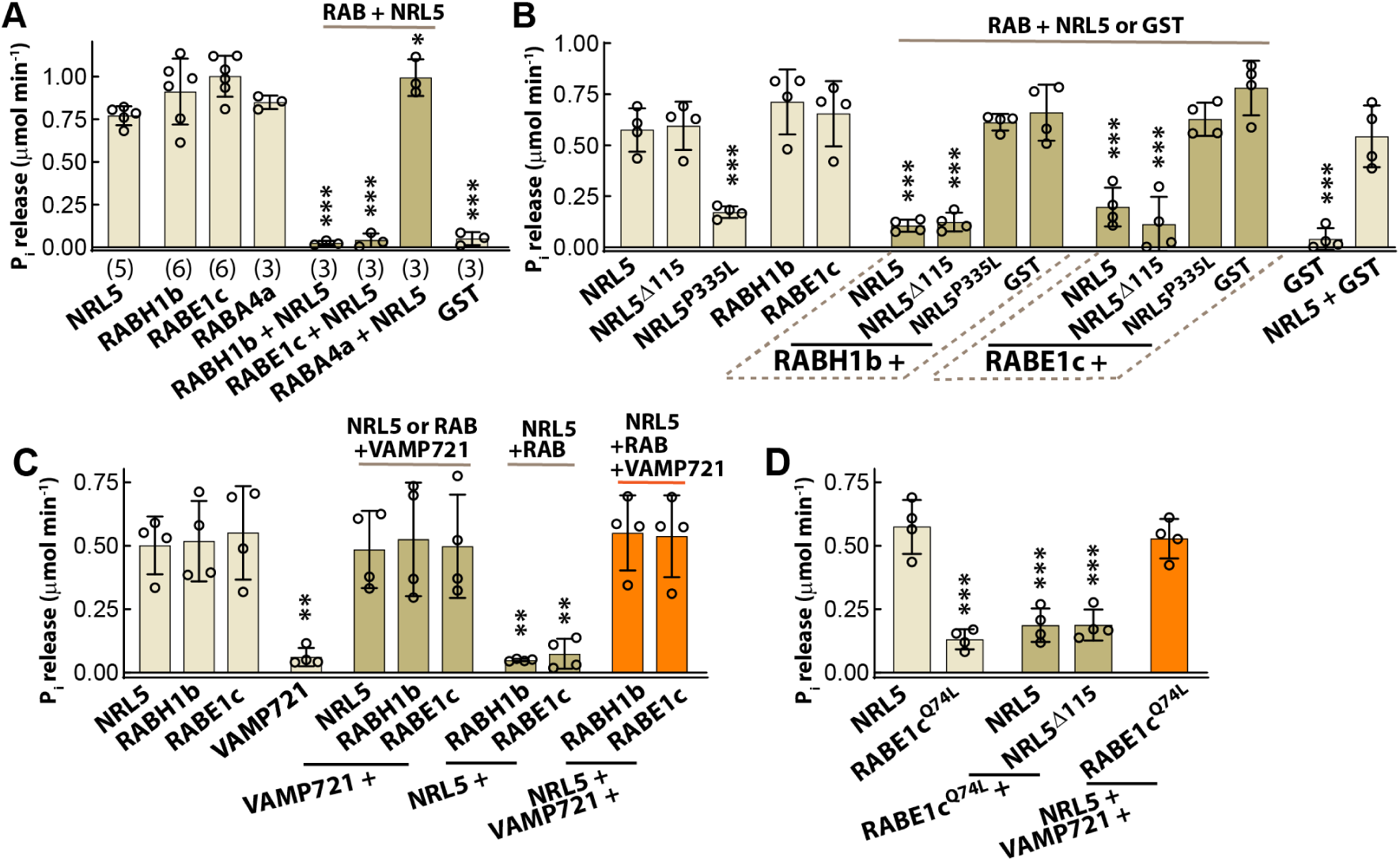
NRL5 and RABH1b/RABE1c inhibit each other’s GTPase activity but this inhibition can be blocked by VAMP721. (**A**) NRL5 and RABH1b/RABE1c inhibited each other’s GTPase activity while the combination of NRL5 and RABA4a did not lead to the same inhibition of GTPase activity. This was consistent with interaction of NRL5 with RABE1c and RABH1b (Fig. 2A) but the lack of detectable interaction between NRL5 and RABA4a (Fig. S7B). Reactions were conducted with 1 μM of each protein and 5 mM GTP (20 μl reaction volume) with P_i_ release quantified after 30 min. Data were analyzed by ANOVA comparison to NRL5 with n values indicated in parentheses. Error bars show S.D. (**B**) Inhibition of RABE1c/RABH1b GTPase activity did not require the BTB domain of NRL5 (NRL5^Δ115^ had the first 115 amino acids of the BTB domain removed) but did require a functional NPH3 domain as NRL5^P335L^ was unable to inhibit RAB GTPase activity. Assays contained 0.25 μM of each protein and 15 mM GTP with P_i_ release quantified after 30 min. Data were analyzed by ANOVA comparison to NRL5 (n =4), error bars indicate S.D. Data are from one experiment out of three independent experiments conducted with consistent results. (**C**) Addition of VAMP721 alleviated the inhibition of GTPase activity caused by NRL5-RABE1c/RABH1b interaction. The three proteins were present at equi-molar ratio (0.25 μM of each protein). Other experimental conditions and data analyses were the same as for B. (**D**) Constitutive active RABE1c (RABE1c^Q74L^) could no longer hydrolyze GTP but did still inhibit NRL5 GTPase activity in a manner that was alleviated by the presence of VAMP721. All proteins were present in equi-molar ratio (0.25 μM of each protein) with other conditions and analysis the same as in B and C. Note that the data of NRL5 assayed alone shown in this panel is the same as in panel B as the assays were conducted in same set of experiments. The NRL5 alone data are shown again here for clarity of presentation. *, **,*** and indicate P ≤ 0.05, P ≤ 0.005 and P ≤ 0.001, respectively.

Further GTPase assays found that VAMP721 did not influence the GTPase activity of either NRL5 or RABE1c/RABH1b (Fig. 4C). However, addition of VAMP721 to NRL5-RABE1c or NRL5-RABH1b mixtures (with each of the three proteins present in an equal molar amount) led to a partial restoration of GTPase activity (Fig. 4C). We also found that constitutive active RABE1c (RABE1c^Q74L^), which is locked in the active state but has no GTP hydrolysis activity, inhibited NRL5 activity (Fig. 4D). Adding VAMP721 to NRL5-RABE1C^Q74L^ mixtures led to restoration of GTPase activity (Fig. 4D), confirming that at least a portion of the GTPase activity in NRL5-RABE1c-VAMP721 mixtures is from NRL5.

Together with the binding assays results (Fig. 2 and Fig. S7), these data (Fig. 4) indicate that VAMP721 (and likely VAMP722) can compete for interaction with NRL5 or RABE1c/RABH1b while not inhibiting the GTPase activity of either one. Thus, when all three proteins were mixed for GTPase assay, interaction of some NRL5 and RAB with VAMP721/722 allowed for recovery of GTPase activity with part of this activity coming from NRL5 and part from the RAB GTPase. These data raise the possibility that the dynamics of NRL5 and RABE1c/RABH1b interaction with each other versus interaction with VAMP721/722 could influence GTPase activity and RAB activation state.

### NRL5 co-localization with RABE1c and VAMP721, as well as accumulation in BFA bodies, further suggests a trafficking related function

By a combination of transformation and crossing, we generated plants expressing three constructs, *NRL5_pro_:NRL5-CFP*, *35S:YFP-RABE1c* and *35S:mCherry-VAMP721,* and analyzed the co-localization patterns of these three proteins. In hypocotyl cells of unstressed plants, NRL5 co-localized with RABE1c along the outside of the cell, close to the plasma membrane (Fig. 5A, Control). NRL5 partially co-localized with VAMP721 on foci in the cell interior (indicated by white arrow in control images, Fig. 5A). Although both NRL5 and VAMP721 could be detected on these foci, the VAMP721 signal was consistently higher than the NRL5 signal. Time lapse analysis showed that the internal foci of VAMP721 and NRL5 were mobile (Fig. S12; Supplemental Movie S1-S8). The smaller foci of NRL5 along the plasma membrane, where colocalization with RABE1c mainly occurred, were not permanent, but also did not exhibit substantial lateral movement along the plasma membrane. Note that the localization of RABE1c primarily along the plasma membrane was consistent with previous characterization of RABE1d (*33*). The VAMP721 localization we observed in the unstressed control was also consistent with previous reports (*34-36*).

**Fig. 5:**
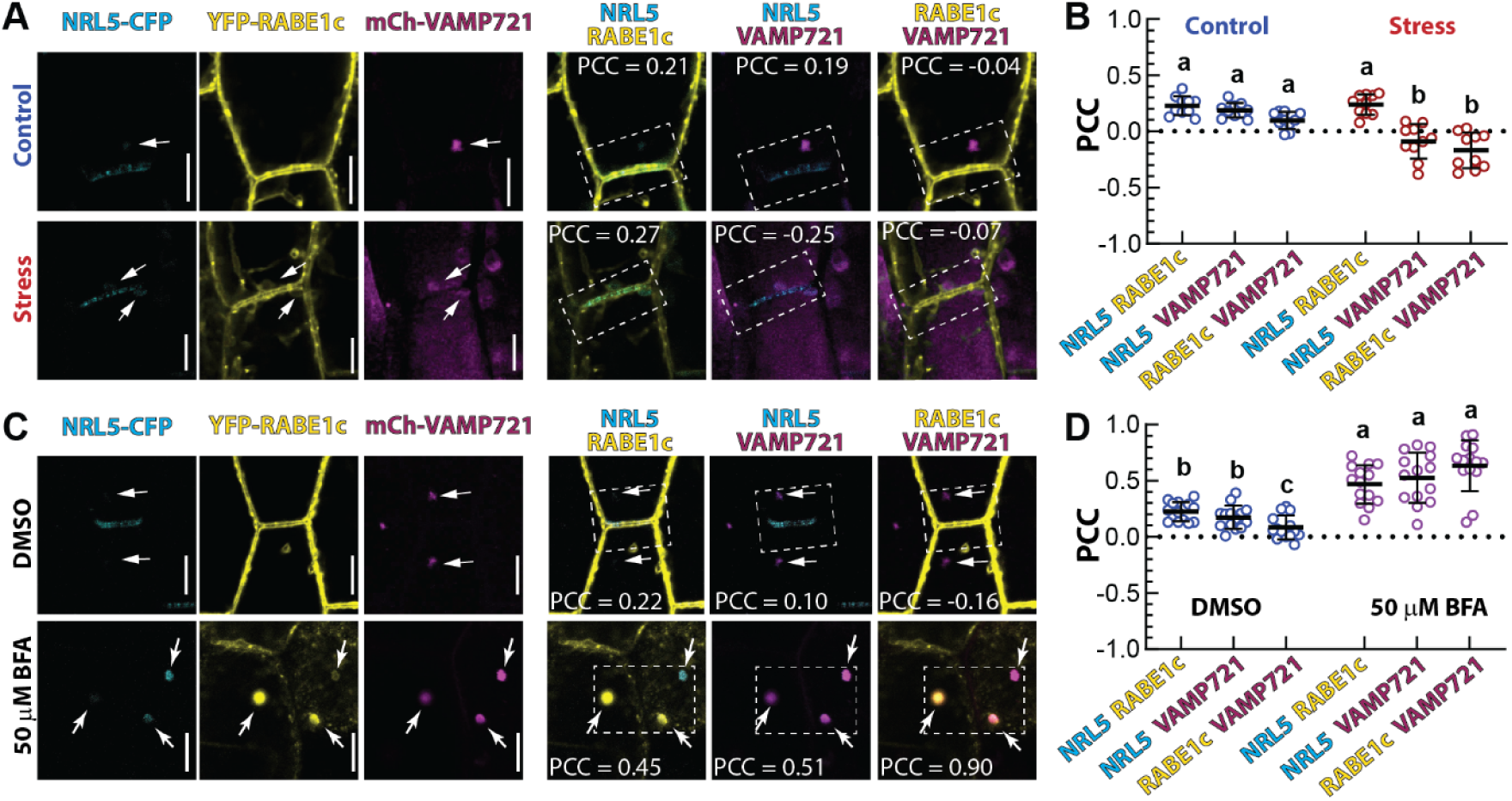
NRL5, RABE1c and VAMP721 colocalization and accumulation in BFA bodies. (**A**) Representative images of hypocotyl cells from a transgenic line expressing *NRL5_pro_:NRL5-CFP*, *35S:YFP-RABE1c* and *35S:mCherry-VAMP721*. Single channel images as well as merged images of two channels are shown. Five-day-old seedlings were transferred to fresh control agar plates (Control) or to -0.7 MPa PEG-infused agar plates (Stress) for four days before imaging. Dashed line boxes indicate the area selected for Pearson Correlation Coefficient (PCC) analysis to quantify co-localization between the indicated two proteins. Arrows indicate an example of foci in the cell interior where VAMP721 was observed to co-localize with a smaller amount of NRL5, but not RABE1c (see further analysis in Fig. S13 and Supplemental Movies S1 to S8). PCC values for each of the merged images are shown. Scale bars indicate 10 μm. (**B**) PCC analysis of the indicated protein pairs in control and stress treatment. Ten cells were analyzed from multiple hypocotyls for each genotype and treatment (n =10). Black bars indicated the mean and error bars show S.D. Data not sharing the same letter are significantly different from one another (ANOVA, P ≤ 0.05). (**C**) Representative images of plants from the unstressed control treated with 50 μM Brefeldin A (BFA) or DMSO (solvent control) for 2 h. In the DMSO treatment, arrows indicate VAMP721 foci in the cell interior. These foci would also transiently contain lower levels of NRL5. In the BFA treatment, arrows indicate BFA bodies. Dashed boxes indicate the area used for PCC analysis. Scale bars indicate 10 μm. (**D**) PCC analysis of the DMSO and BFA treatments. Data format and analysis are as described for B (n = 14).

During low ψ_w_ stress, NRL5 and RABE1c had similar colocalization pattern close to the plasma membrane and we could also detect structures containing NRL5, RABE1c and VAMP721 close to the plasma membrane (white arrows in stress images, Fig. 5A). These membrane-adjacent sites of co-localization were less motile than the internal foci of VAMP721 and NRL5 (Fig. S12, Supplemental Movie S1-S8). In general, low ψ_w_ caused VAMP721 to have a more diffuse cytoplasmic localization with fewer of the foci seen in unstressed cells. Quantitative analysis found that low ψ_w_ had little effect on the extent of NRL5-RABE1c colocalization but significantly decreased NRL5-VAMP721 and RABE1c-VAMP721 colocalization (Fig. 5B).

A similar pattern was observed in leaf pavement cells where we detected NRL5-RABE1c colocalization along patches of the plasma membrane in both control and stress conditions (Fig. S13). Leaf pavement cells also had plasma membrane-adjacent structures where both NRL5 and VAMP721 could be found, even though overall co-localization of these two proteins, as indicated by the PCC analysis, was low (Fig. S13). Low ψ_w_ also disrupted VAMP721 localization in leaf cells similar to the effect seen in hypocotyl cells.

The interaction and co-localization of NRL5 with RABs and VAMP721/722, all of which have established function in intracellular trafficking (*34-38*), along with NRL5 GTPase activity, suggested that NRL5 is also associated with intracellular trafficking. In this case, we would expect NRL5 to accumulate in Brefeldin A (BFA) bodies since BFA inhibits post-Golgi secretion and endosomal recycling of plasma membrane proteins (*39*). Consistent with this hypothesis, BFA treatment disrupted the localization of NRL5 along the plasma membrane and caused it to accumulate in BFA bodies where it extensively colocalized with RABE1c and VAMP721 (Fig. 5C, D). The accumulation of NRL5 in BFA bodies further indicates that NRL5 participates in intracellular trafficking and show that its localization along the edge of the cell, and its polarity (see below), depend on trafficking.

### A functional NPH3 domain and RAB interaction are required for NRL5 polarity

In transgenic plants where the *nrl5-2* mutation was complemented with *NRL5_pro_:NRL5-YFP*, the NRL5-YFP protein accumulated mainly in epidermal cells of leaf and hypocotyl as well in root stele tissue (Fig. S14). In these cells, NRL5 had a strikingly polar localization. In leaf pavement cells, NRL5 was typically found along one edge of a cell and did not localize next to stomata or other small, non-lobed cells. In many cases, NRL5 was localized on opposite ends of adjacent cells and seemed to trace a path through the leaf epidermis (Fig. S14B). In hypocotyl epidermis, NRL5 concentrated at the rootward (bottom) side of cells (Fig. S14). A similar pattern was observed in root stele cells.

Both low ψ_w_ stress and the P335L mutation diminished NRL5 polarity. In leaf pavement cells, this was seen as the formation of multiple patches of NRL5 or NRL5^P335L^ along the periphery of the cell rather than the single patch of NRL5 observed in unstressed plants (Fig. 6A, Fig. S15). Consistent with this, low ψ_w_ and the P335L mutation significantly decreased the skewness of the YFP signal (Fig. 6B). It was particularly striking that NRL5^P335L^ would localize alongside guard cells, and other small non-lobed cells, in both control and low ψ_w_ stress conditions (Fig. 6A). Such localization next to stomates or small cells was not observed for wild type NRL5 in unstressed plants and only occasionally observed in plants at low ψ_w_ (Fig. 6A and Fig. S15). In hypocotyl cells, NRL5^P335L^, and NRL5 at low ψ_w_, seemed to start “climbing” the edges of the cells, as evidenced by a decreased ratio of base-versus-edge fluorescence intensity (Fig. 6C and D). In the root tip, loss of polarity was not as apparent and the main effect of low ψ_w_ and the P335L mutation was to decrease the level of NRL5 protein and confine it to fewer cells close to the quiescent center (Fig. S16).

**Fig 6:**
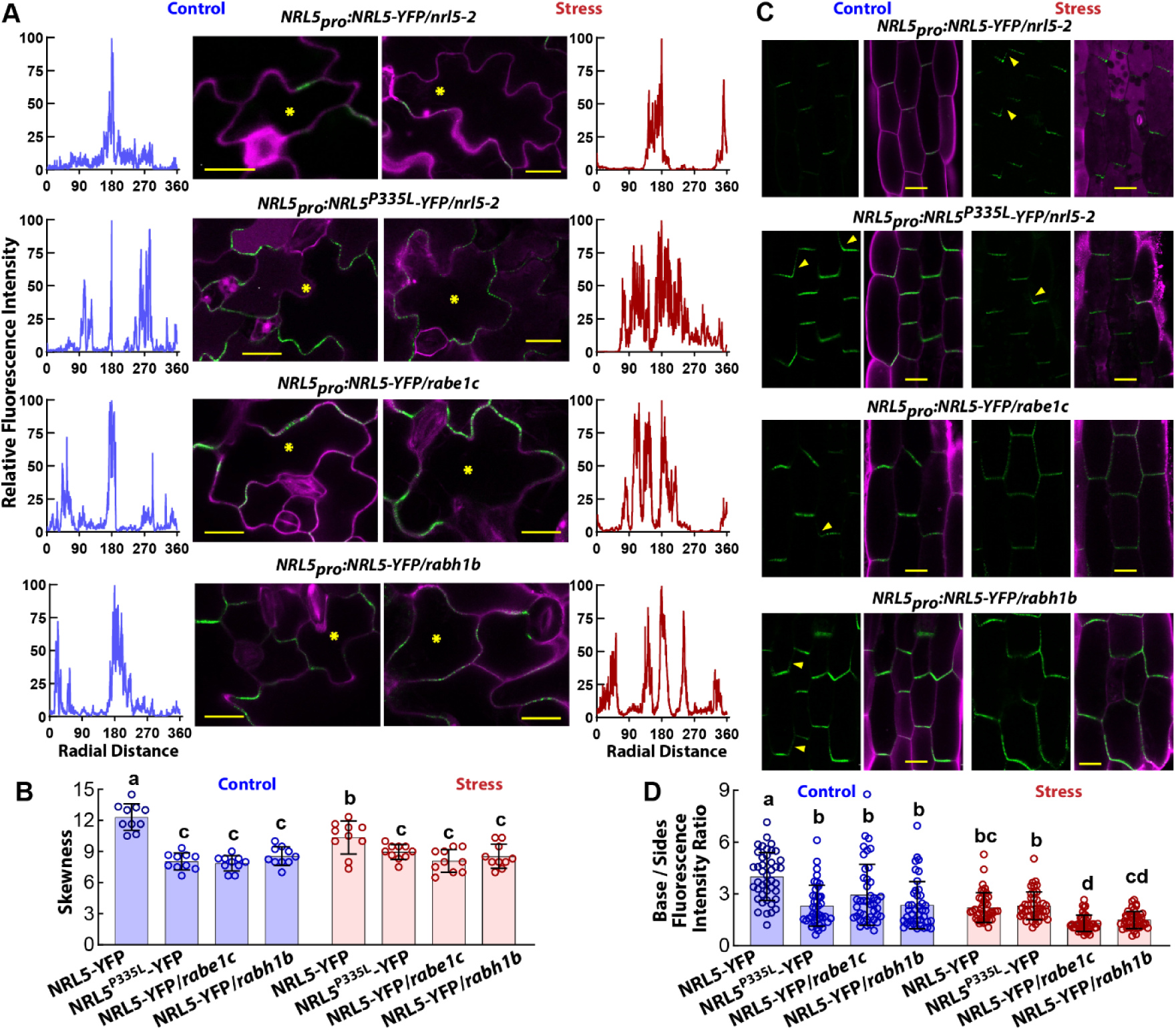
NRL5 polarity is disrupted by the P335L mutation or by lack of RABE1c/RABH1b. (**A**) Representative images showing NRL5-YFP and NRL5P335L-YFP localization in leaf pavement cells. The *NRL5_pro_:NRL5-YFP* construct was also introduced into the *rabe1c* and *rabh1b* genetic backgrounds by crossing. Graphs on the left and right side of the images show relative fluorescence intensity along a line scan tracing of the cell periphery. For each cell analyzed, the point of highest fluorescence intensity was set to 100 and other points normalized to that value. The normalized data were plotted on the basis of radial distances (360°) around the cell with the point of highest fluorescence set to 180°. Note that the correspondence of radial distance to physical distance varies for cells of different sizes. Cells were imaged four days after transfer of five-day-old seedlings to either fresh control media or low ψ_w_ (-0.7 MPa). Scale bars indicate 20 μm. Additional images of NRL5 localization in pavement cells can be found in Fig. S15. (**B**) Skewness analysis of leaf pavement cells shows that NRL5^P335L^ or introduction of the *rabe1c* or *rabh1b* mutations leads the NRL5-YFP signal to be more widely distributed and less concentrated than for NRL5 in the wild type background. This was consistent with the reduced polarity seen in A and Fig. S14. Low ψ_w_ also caused a reduced skewness. Each data point is the average to three cells/fields from the same leaf for each of ten leaves, from multiple seedlings, analyzed (n = 10). Data were analyzed by ANOVA with groups not sharing the same letter being significantly different (corrected P ≤ 0.05). Data are shown as means ± S.D. (**C**) Representative images of NRL5-YFP localization in hypocotyl cells of the same genotypes and treatments described in A. Yellow arrows indicate examples of NRL5 or NRL5^P335L^ localizing on the lateral sides of cells rather than only on the bottom (rootward) side. More substantial loss of polarity is seen for NRL5 in the *rabe1c* or *rabh1b* backgrounds after exposure to low ψ_w_ (-0.7 MPa, 4 days). Scale bars indicate 20 μm. (**D**) Ratio of fluorescence intensity along the base (rootward side) of a hypocotyl cell compared to the summed fluorescence intensity on both lateral sides of the cell. Each data point is for an individual cell with six cells analyzed from each of seven hypocotyls for each genotype and treatment (n = 42). Data were analyzed by Kruskal-Wallis test with groups not sharing the same letter being significantly different (corrected P ≤ 0.05). Data are shown as means ± S.D.

We also constructed plants where *NRL5_pro_:NRL5-YFP* was expressed in either the *rabe1c* or *rabh1b* mutant backgrounds. In leaf pavement cells, absence of either RAB resulted in partial loss of NRL5 polarity in both control and low ψ_w_ conditions (Fig. 6A and B, Fig. S15). Typically, several patches of NRL5 along the plasma membrane were observed in the *rabe1c* or *rabh1b* mutant background, rather than the single patch of NRL5 found in wild type cells. In hypocotyl cells of *rabe1c* or *rabh1b*, NRL5 “climbed” the sides of the cells to even greater extent than NRL5^P335L^ and this was further exacerbated by low ψ_w_ (Fig. 6C and D). In root cells, *rabe1c* had a reduced level of NRL5 protein (Fig. S16). Together, these data indicated that NRL5 interaction with RABs contributed to maintenance of its polarity. This occurred despite the fact that RABE1c itself was not polarly localized (Fig. 5).

The loss of polarity for NRL5^P335L^ was even more apparent in rBiFC assays of NRL5 versus NRL5^P335L^ self-interaction. NRL5 homocomplexes detected by rBiFC were localized on one edge of leaf pavement cells (Fig. S8E). In contrast, the rBiFC data showed that NRL5^P335L^ homocomplexes exhibited a complete loss of polarity. The NRL5^P335L^ loss of polarity in these assays was more extensive than in stable transgenic plants, likely because the rBiFC visualized only NRL5^P335L^ homocomplexes whereas in stable transgenic lines NRL5^P335L^-YFP could form heterocomplexes with other NRLs and this allowed some degree of polarity to be maintained. It should also be noted that RPT2 self-interaction was not polar (Fig. S9B). This suggested that polar localization may be a specific feature of a subset of NPH3-domain proteins including NRL5.

Both the localization of NRL5 along the edge of the cell and its inclusion in BFA bodies suggested that it may be associated with the plasma membrane. Consistent with this hypothesis, NRL5 partially co-localized with a plasma membrane marker (Fig. S17A and B). The P335L mutation and low ψ_w_ diminished NRL5 plasma membrane co-localization. In all our microscopy observations, it was also apparent that NRL5 was present as puncta along the plasma membrane, or in close vicinity to it. However, NRL5 did not co-localize with a plasmodesmata marker (Fig. S17C and D), indicating that NRL5 puncta are of other origin, likely related to vesicle trafficking. Together these results demonstrated that that a functional NPH3 domain is required for NRL5 polar localization and co-localization with plasma membrane and that RAB interactions mediated by the NPH3 domain are important for NRL5 polarity.

### *nrl5-2* low ψ_w_ sensitivity requires VAMP721/722 and RABE1c, but NRL5 does not affect their protein abundance

Given these functional connections among NRL5 and its interaction partners, we also tested their genetic interaction in low ψ_w_ resistance. Most strikingly, *nrl5-2vamp721* and *nrl5-2vamp722* alleviated the low ψ_w_ sensitivity phenotype of *nrl5-2* (Fig. 7A-C). Likewise, *nrl5-2rabe1c* was phenotypically similar to the *rabe1c* single mutant, albeit that the *rabe1c* mutant was itself somewhat more sensitive to low ψ_w_-induced growth inhibition than wild type (Fig. 7A-C). These results indicated that *nrl5-2* low ψ_w_ hypersensitivity was caused by improper function of VAMP721/722, and RABE1c during low ψ_w_ stress. The *nrl5-2rabh1b* results were less clear-cut as *rabh1b* had constitutively reduced growth (Fig. 7), consistent with report that RABH1b is involved in trafficking of cellulose synthase to the plasma membrane and that *rabh1b* has cell wall defects which limit its growth (*38*). However, in this case also the *nrl5-2rabh1b* double mutant more closely resembled *rabh1b* than *nrl5-2* (Fig. 7)

**Fig. 7:**
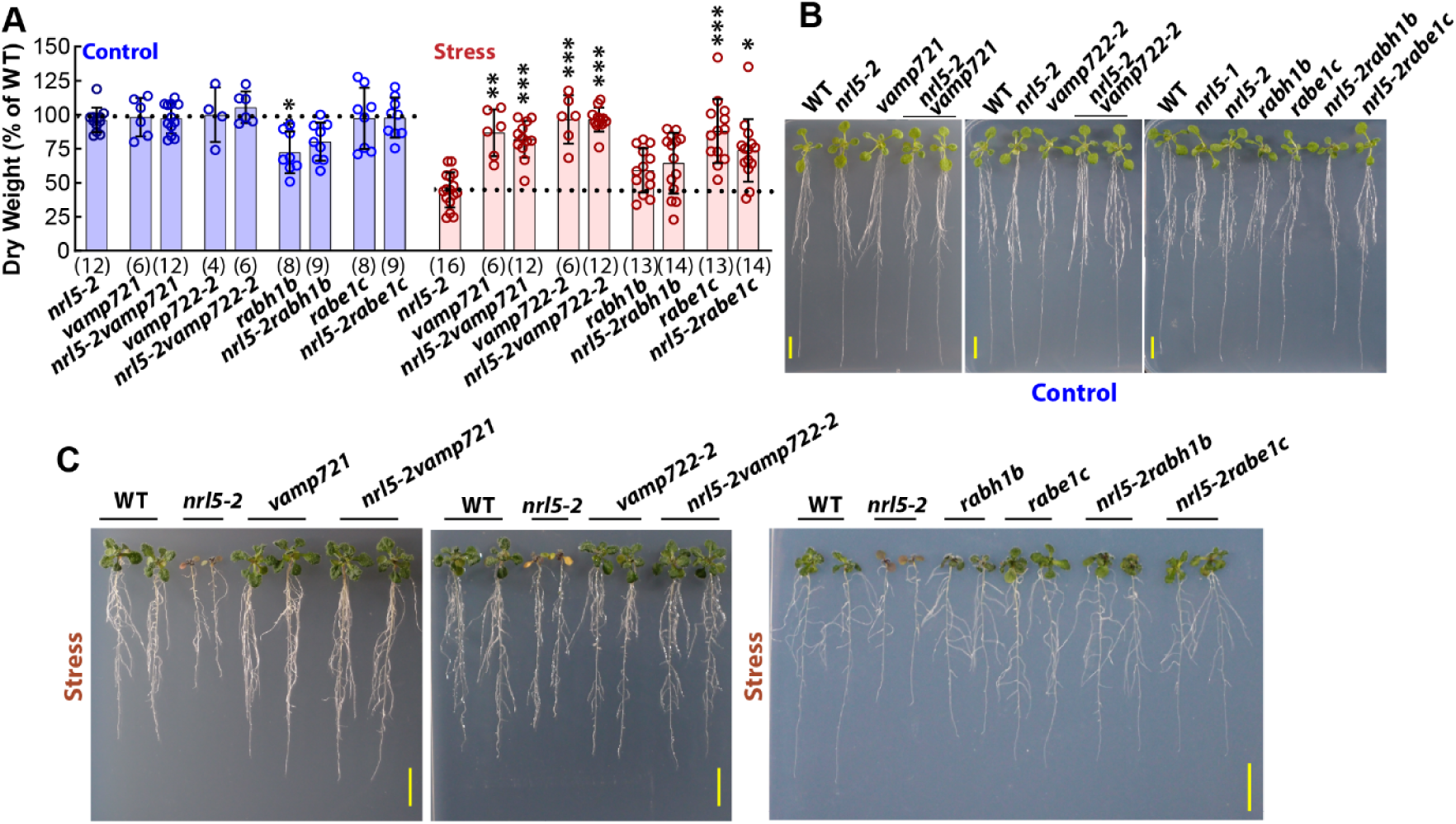
The low ψ_w_-hypersensitive phenotype of *nrl5-2* depends upon the presence of VAMP721/722 or RABE1c/RABH1b. (**A**) Seedling dry weight analysis found that double mutants of *nrl5-2* with *rabe1c/rabh1b* or *vamp721/722* are phenotypically similar to the *rab* or *vamp* single mutants. Data are mean ± S.D. (n values indicated in parentheses, each data point is from one assay plate where the test genotype was grown alongside the wild type). Data are combined from two or three independent experiments. Dry weights were measured 5 days after transfer of 5-day-old seedlings to fresh control plates or 10 days after transfer to low ψw plates (-0.7 MPa). Data were analyzed by Kruskal-Wallis test with comparison to *nrl5-2* in the same treatment (n values shown in parentheses). ** and *** indicate P ≤ 0.01 or P ≤ 0.001, respectively. Dashed lines indicate the wild type value for each treatment. (**B**) Representative images of wild type and mutant seedlings 5 days after transfer to control plates. Scale bars indicate 1 cm. (**C**) Representative images of seedlings 10 days after transfer to -0.7 MPa low ψw stress. Scale bars indicate 1 cm.

Quantitative proteome profiling of wild type and *nrl5-2* in the unstressed control or after four days exposure to -0.7 MPa low ψ_w_ stress identified numerous low ψ_w_ -induced changes in protein abundance in wild type and *nrl5-2* (Data S3-S5). However, none of the RABE1 or RABH1 proteins, or VAMP721/722, were among the differentially abundant proteins in *nrl5-2* (Fig. S18A). Thus, our data gave no indication that NRL5 targeted RABE1c, RABH1b or VAMP721/722 for degradation. NPH3 acts as an E3 ligase adaptor protein to target PHOT photoreceptors for degradation; however, NPH3 interacts with PHOTs via its C-terminal coiled-coil domain while NRL5 interacts with RABE1c, RABH1b and VAMP721/722 via its NPH3-domain. Together these data indicated that NRL5 interaction with RAB and VAMP proteins has a different purpose other than controlling protein stability.

### Patterns of altered protein abundance in *nrl5-2* during low ψ_w_ stress

To further analyze the *nrl5-2* proteomics data, hierarchical clustering was performed for proteins with substantially altered abundance (log2 fold change ≥ 0.5 and P ≤ 0.05) in wild type stress versus control or in *nrl5-2* compared to wild type in the stress or control treatments. Overall, *nrl5-2* had many more proteins of increased abundance than of decreased abundance. The clustering analysis made it clear that many of the proteins increased in *nrl5-2* during low ψ_w_ were down-regulated or unchanged by low ψ_w_ in wild type (Fig. S18B). We hypothesize that most of these increases in protein abundance were indirect effects of greater stress-induced damage in *nrl5-2* compared to wild type. However, we do not rule out the possibility that NRL5 can also act as an E3 ligase adaptor to directly control the abundance of some of these proteins. There was also a smaller group of proteins, including P5CS1, that were up-regulated by low ψ_w_ in wild type and further increased in *nrl5-2* (Fig. S18; detailed list and heat map shown in Fig. S19). This group included a number of dehydrin and Late Embryogenesis Abundant (LEA) proteins that are typically involved in protecting cellular structure during severe dehydration associated with abiotic stress or seed development.

Interestingly, there was also a group of proteins that were of increased abundance in wild type after low ψ_w_ treatment but failed to increase in *nrl5-2* (“Pattern I” in Fig. S18B and listed individually in Fig. S20A). A high portion of these (14 out of 30) had predicted localization in the plasma membrane or extracellular space. This is likely an underestimate of the portion of plasma membrane or extracellular proteins in this cluster since proteins with predicted localization in plastid (NRL11), E.R. (AT1G43910) or vacuole (TOUCH 3/CALMODULIN-LIKE 12) may in fact be plasma membrane or extracellular proteins as well. This suggested the hypothesis that the decreased abundance of these proteins in *nrl5-2* was caused by disrupted trafficking that prevented them from reaching their proper subcellular localization where they are most stable; or, in the case of plasma membrane proteins, causes them to be more rapidly removed for degradation. While such an effect would be consistent with our overall findings, we emphasize that this is a hypothesis which awaits further investigation. It was also interesting to note that this set of proteins included two glutamine synthetases (GLN1;1 and GLN1;2) that may contribute to the proline accumulation phenotype of *nrl5* mutants by controlling the amount of glutamate available as a substrate for proline synthesis.

There was also a group of proteins that were down regulated or unchanged by low ψ_w_ in wild type but strongly increased in *nrl5-2* (Pattern II, Fig. S18B, Fig. S20B). It was particularly striking that this group included several Cruciferins that were among the most strongly increased proteins in *nrl5-2*. Cruciferins are seed storage proteins that are normally localized to protein storage vacuoles but may also be secreted if not trafficked properly (*40*). There is also some evidence that they can have lipase activity after proteolytic processing (*41*). It will be of interest to determine if this striking induction of cruciferins and other seed-related proteins is an indirect effect of stress-induced damage in *nrl5-2* or is part of a developmental and signaling pattern that has gone awry and is contributing to the low ψ_w_ hypersensitivity of *nrl5* mutants.

## Discussion

NPH3-family proteins have been implicated in key cellular processes, most notably phototropism and auxin signaling (*22, 27, 28*). However, molecular function of the NPH3 domain, the defining feature of this protein family, has remained unknown. Our results show that the NPH3-domain is a plant-specific type of GTPase. Most GTPases are involved in intracellular trafficking (*29, 42*). This, alongside NRL5 interaction with RABH1b/RABE1c and VAMP721/722 as well as their cross-regulation of each other’s activities, indicates that NRL5 has previously unrecognized functions in intracellular trafficking (see model and description in Fig. S21). The high degree of NPH3-domain conservation across more than thirty NPH3 family members in Arabidopsis, and our finding that multiple NPH3-domain proteins have GTPase activity, indicates that GTPase activity, and possibly trafficking function as well, is a core feature of this protein family. NRL5 is part of the NPY subgroup of NPH3-domain proteins, of which NPY1 to 5 are polarly localized and involved in maintaining polarity of PIN auxin transporters (*27, 28, 43*). For NPH3 itself, it has been noted that its localization at the plasma membrane versus undefined bodies close to plasma membrane, determines its activity in phototropism (*44*). Discovery of NPH3-domain GTPase activity and interaction with trafficking proteins will allow such observations to be critically re-evaluated and put into a functional context. This new knowledge of NPH3 domain molecular function will also facilitate analysis of the majority of NPH3-domain proteins whose cellular and physiological functions are not yet known. More broadly, the finding that plants have maintained a type of GTPase not found in metazoans illustrates how plants have adapted the core conserved trafficking machinery to their unique needs.

While the GTPase activity and trafficking association provide a broad principle about NPH3-domain function, the dramatic low ψ_w_ sensitivity phenotype resulting from mutation of a single NPH3-domain protein speaks to specificity among these proteins. The interaction of NRL5 with RABE1c and RABH1b, as well as proteome analysis of *nrl5-2*, support a hypothesis whereby NRL5 affects trafficking and abundance of proteins involved in stress signaling at the plasma membrane-cell wall interface (Fig. S21). We also made the surprising observation that *nrl5-2rabe1c,* as well as *nrl5-2vamp721* and *nrl5-2vamp722,* had a recovery of low ψ_w_ resistance compared to *nrl5-2*.

### Implications of NRL5 genetic and physical interactions with RABs and VAMP721/722

To interpret the recovery of low ψ_w_ resistance seen in *nrl5-2rabe1c* result, it is helpful to consider previous report that ectopic expression of constitutive active RABE1d (RABE1d^Q74L^) increased secretion of Pathogenesis-Related 1 (PR1) and a number of other unidentified proteins. The authors concluded that increased level of active RABE1d activated multiple protein secretion pathways (*33*). RABE1d is nearly identical to, and at least partially redundant with, RABE1c (*29, 33, 45*). Speth et al. *(33)* also found that transgenic lines where expression of multiple *RABE1* genes was reduced by co-suppression had reduced growth and cell division. They proposed that this was due to reduced trafficking of proteins needed for cell plate and cell wall formation. Other studies have also implicated both RABE1c and RABH1b in trafficking and secretion needed to build or maintain the cell wall (*37, 38*). Interestingly, Speth et al. (*33*) also observed that while RABE1d was normally found all along the plasma membrane, bacterial inoculation caused it to adopt a polar localization along one side of leaf mesophyll cells. Putting these results together with our genetic and biochemical data, we hypothesize that the *nrl5-2rabe1c* double mutant had a recovery of low ψ_w_ resistance because it no longer had an excess of inactive RABE1c at certain key stages or locations of vesicle trafficking where NRL5 normally holds RABE1c in its active state by blocking its GTPase activity.

In the case of VAMP721/722, it is very interesting that NRL5 interacted with both the SNARE domain and Longin domain. The Longin domain of VAMP721 can fold back over the SNARE domain to block it from interacting with the SNARE domains of cognate Q-SNAREs during the first steps of vesicle fusion with a target membrane. For metazoan VAMP7, the open versus closed configuration is controlled by interaction with regulatory proteins as well as phosphorylation of the Longin domain (*46, 47*). Recently it has been shown that in plants, transient phosphorylation of the VAMP721 Longin domain is important to promote the open configuration needed for Q-SNARE interaction (*30, 48*). Surprisingly, expression of phosphomimic VAMP721, where the open configuration is predominant, inhibited secretion to the cell wall and reduced growth (*30*). This seemed to be caused by constitively open VAMP721 interacting with Q-SNAREs at inappropriate places or times such that membrane fusion could not be completed, thus blocking secretory trafficking (*30*). This model can explain the surprising reversal of *nrl5-2* low ψ_w_ hypersensitivity in *nrl5-2vamp721* and *nrl5-2vamp722*. If NRL5 is involved in keeping VAMP721/722 in the closed configuration, then the absence of NRL5 may lead to VAMP721/722 adopting the open configuration inappropriately and thus inhibiting trafficking, similar to the effect seen for phosphomimic VAMP721 (*30*). Combining the *nrl5-2* mutation with either *vamp721* or *vamp722* reduced the dosage of open configuration VAMP721/722 thus allowing trafficking to procede and the *nrl5* low ψ_w_ hypersenstive phenotype to be alleviated. Here again, we readily acknowledge that this is a speculative model that requires testing. We hope that our results will guide and stimulate such experiments. It should also be noted that there is much literature about VAMP721/722 involvement in vesicle targeting and protein secretion to the plasma membrane and cell wall (*34-36, 49*). In particular, VAMP721 can coordinate vesicle fusion with opening of potassium channels to maintain osmotic balance across the plasma membrane (*48, 50*). Thus, it is perhaps not so surprising that VAMP721/722 could influence the response to low ψ_w_ where the osmolarity of cells must substantially increase to allow continued water uptake while still maintaining an appropriate coordination between cell volume and solute deposition. Overall, our data showing new connections of NRL5 to trafficking are consistent with at least one other study that found an NPH3-domain protein localized to vesicles (*51*).

Our data also show that NRL5 interaction with RABE1c and RABH1b is crucial to maintain its polarity and it will be of interest to see if this is related to observation that RABE1d becomes polarized under certain conditions (*33*). It will be of interest to determine how NRL5-RAB interaction fits into broader hypothesis of cell polarity maintenance (*43*) and how these interactions affect trafficking and localization of proteins that may have more direct involvement in stress sensing and signaling. The P335L mutation disrupts both NRL5 GTPase activity and its polarity; thus, it is not yet clear whether GTPase activity is required for NRL5 polar localization. The finding that RAB interaction is needed for NRL5 polarity is of particular interest in terms of how it may be relevant to NPYs and their function in maintaining polarity of PIN auxin transporters (*27, 28*). Note that even though NRL5 is closely related to NPYs, hence the NPY8 alternative name for NRL5 (*43*), there has not to our knowledge been any study of whether NRL5 has similar protein interactions as the better studied NPY1 to NPY5. It is also possible that NRL/NPYs form heterodimers via their BTB domains; although, here again there has been little previous discussion of such heterodimer formation and how it may impact NRL/NPY function.

### Aberrant low ψ_w_ response and proline metabolism in *nrl5* mutants

Another major finding of our experiments is that NRL5 mutants are dramatically hypersensitive to low ψ_w_. It is important to note that this low ψ_w_ hypersensitivity is not because *nrl5* mutants fail to respond to low ψ_w_. Rather it is because they have an aberrant response to low ψ_w_ that is much more damaging to the plant than the low ψ_w_ stress itself. A key part of this aberrant response is mis-regulation of proline metabolism. The proline metabolism profile of *nrl5* mutants (elevated P5CS along with high ProDH activity but no increase of P5CDH) resembles the proline metabolism profile required for cell death during the hypersensitive response to pathogen infection (*13, 14, 52, 53*). Pathogen perception and hypersensitive response require sensing and trafficking at the cell wall-plasma membrane interface, such as the secretion of PR1 mentioned above (*33*). For *nrl5* mutants, one possible reason for their aberrant low ψ_w_ response is that the cell wall-plasma membrane interface is more susceptible to damage, or has changed in such a way that the plant misinterprets the low ψ_w_ stimulus and mounts a hypersensitive-like response. The cell wall and plasma membrane proteins of altered abundance in *nrl5-2* provide a good starting point to look for proteins that have previously unrecognized roles in low ψ_w_ signaling and whose trafficking and localization may be particularly dependent upon NRL5. However, it must be noted that because the P335L mutation affects NRL5 GTPase activity and NRL5 polar localization, as well as RAB and VAMP721/722 interaction, it is not yet clear which of these aspects of NRL5 function are most important for its effect on drought response. Further structure-function studies of NRL5 will be needed to unravel these different aspects of NRL5.

One outcome of the aberrant low ψ_w_ response of *nrl5* mutants is the over production of ROS and accumulation of P5C (as measured by o-AB reaction). Indeed, the dramatic low ψ_w_ hypersensitivity of *nrl5* mutants provided a direct demonstration that there is potential for damaging ROS and P5C production whenever proline accumulates. Drought-related signaling pathways, which we now know require NRL5 to function properly, suppresses the damaging aspects of proline catabolism and allows proline to safely accumulate (*3*). Blocking ROS and P5C production in the *nrl5-2prodh1-1* double mutant alleviated the *nrl5-2* low ψ_w_ hypersensitive phenotype. Interestingly though, *nrl5-2p5cdh* had similar, or even greater, low ψ_w_ hypersensitivity as *nrl5-2* but did not accumulate higher levels of P5C than *nrl5* or *p5cdh*1 single mutants. Thus, it will also be of interest to determine whether a proline-P5C cycle, which serves to amplify proline-dependent ROS production in mammalian cells (*15, 16*), also exists in plants (*53, 54*) and whether such a cycle contributes to the low ψ_w_ hypersensitivity of *nrl5*. It will also, of course, be of interest to identify the specific signaling mechanisms that connect NRL5 GTPase activity and trafficking-related function in the vicinity of the plasma membrane to increased proline catabolism in the mitochondria. Since ProDH activity was increased in NRL5 without a detectable increased in ProDH protein level, this may include post-translational regulation of ProDH, which is so far little understood.

While the above hypotheses are promising pathways for future study, they focus on a proposed role of NRL5 in trafficking of proteins to the plasma membrane and cell wall. We also emphasize that we cannot rule out other trafficking-related functions of NRL5. We also do not rule out the possibility that NRL5 uses its BTB domain to act as a ubiquitin E3 ligase adaptor to target proteins for degradation in a manner similar to NPH3 control of Phototrophin degradation (*25*). NPH3 interacts with Phototropin via its C-terminal coiled-coil domain and interacts with Cullin 3 via its BTB domain (*25, 55*). The unstructured C-terminal domain is not conserved among NPH3 domain proteins and, in the trafficking context, may be the basis for NRL5 specificity in affecting trafficking of stress-related proteins. In the protein degradation context, divergence of the C-terminal unstructured domain may be a basis for NRLs to target specific proteins for degradation. This key difference among NRLs, along with the role of heterodimerization between different NPH3-domain proteins, is of interest for further investigation to understand why plants need to have so many NPH3-domain proteins. Nearly all NPH3-domain proteins have an N-terminal BTB domain and it is yet unclear whether its primary role is to mediate dimerization of NPH3 domain proteins or mediate interaction with Cullin 3 E3 ligase complexes. These two functions are not mutually exclusive and NRL5 effect on drought resistance could involve both trafficking and protein degradation aspects.

The identification of NRL5 as both a critical protein for drought resistance and a trafficking-associated GTPase broadens the role of NPH3-domain proteins and allows for a critical reanalysis of their function in well-established signaling pathways. Our results also provide a dramatic demonstration that proline metabolism is indeed a double-edged sword of plant biology with the potential to protect or disrupt plant function during stress depending upon its regulation (*3*). Further study of NRL5 at both the biochemical and genetic levels will open new approaches to identify drought sensing and signaling mechanisms controlling proline metabolism while also better understanding how plant cells coordinate intracellular trafficking

## Materials and methods

### Plant materials and generation of mutant and transgenic lines

The *nrl5-1* mutant (also referred to as 4090 or 4090-2 in Fig. S1) was obtained by forward genetic screen using an ethyl methanesulphonate (EMS)-mutagenized *ProDH1_pro_:LUC2* population as previously described (*20*). The 4090-2 mutant (in Col-0 background) was crossed to Landsberg *erecta* and homozygous mutants selected from the segregating F_2_ population based on the high *ProDH1_pro_:LUC2* phenotype. Approximately 100 bulked segregant seedlings were used for whole genome sequencing followed by SNP analysis and mapping conducted using the Next Generation Mapping utility (*20, 56*). Candidate genes were initially tested by complementation crosses to the corresponding T-DNA mutant (obtained from the Arabidopsis Biological Resource Center) and conducting luciferase imaging of F_1_ seedlings. A list of T-DNA lines used in this study is given in Data S6. T-DNA mutants were genotyped and homozygosity confirmed using primers designed using the SIGnAL web resource; http://signal.salk.edu/. To construct double mutants, *nrl5-2* was crossed with T-DNA mutants of *p5cs1-4*, *pdh1-2*, *p5cdh*, *rabh1b*, *rabe1c, vamp721 and vamp722-1,* all of which have been characterized in previous publications. The F_2_ generation was screened for homozygous double mutants using primers specific for *nrl5-2* and respective mutants, as listed in Data S6.

To express *NRL5* under the control of its native promoter, a genomic fragment containing the *NRL5* gene body up to the stop codon and 2 kb of upstream of promoter sequence was amplified from Col-0 and cloned into pENTR/D-TOPO (Invitrogen). Site-directed mutagenesis of this clone was conducted to generate a genomic clone of NRL5^P335L^ using the QuikChange II site-directed mutagenesis kit (Agilent) following the manufacturer’s instructions. Both of these clones were moved to binary vector pGWB540 (C-terminal fusion to YFP) and NRL5 clone was also moved to pGWB543 (C-terminal fusion to eCFP) by Gateway LR reaction and then transformed into *Agrobacterium tumefaciens* strain GV3101 for floral dip transformation of the *nrl5-2* mutant or wild type Col-0. For the generation of *35s:YFP-RABE1c* transgenics, RABE1c cDNA was amplified from wild type Col-0 using a two-step nested PCR to add Gateway adaptor sequences follow by recombination into the pEarleyGATE104 vector (35S promoter; n-terminal YFP). For the generation of *35s:mCherry-VAMP721* transgenics, 35S promoter sequence was amplified from the pEarleyGATE104 vector and cloned in pDONR221P4-P1r entry vector, mCherry was amplified from the pCMB-pDESr vector and cloned into pDONR221P1-P2 vector and VAMP721 coding sequence was amplified from Col-0 wild type cDNA and cloned in the pDONR221P2r-P3 vector. The three entry clones were moved into binary vector pB7m34GW by LR reaction and transformed into *Agrobacterium tumefaciens* strain GV3101. All constructs were verified by sequencing.

### Plant growth conditions and stress treatments

Seedling growth and low water potential (ψ_w_) treatment on PEG-infused agar plates was performed as described previously (*57, 58*). Seeds were plated on half-strength Murashige-Skoog nutrient media with the addition of 2 mM MES buffer (pH 5.7) and 1.5% agar, but without addition of sugar. Seeds were stratified for three days and then plates were positioned vertically in a growth chamber at 22°C and continuous light of 90-120 μmol m^-2^ sec^-1^. For growth assay, five-day-old seedlings were transferred to fresh half-strength MS plates (Control, -0.25 MPa) or PEG-infused agar plates of moderate severity low ψ_w_ (-0.7 MPa); note that plates were always freshly prepared and infused with PEG-8000 for 15 h before use to avoid drying of the media and to make plates of consistent ψ_w_. To quantify seedling weights, 4-6 seedlings were weighed together, dried at 65°C and reweighed to obtain the dry weight. The per seedling weight of the test genotype was normalized versus the per-seedling weight of wild type grown on the same agar plate. Seedling weights were measured 5 days after transfer for the unstressed control and ten days after transfer for the low ψ_w_ treatment to allow the wild type to reach approximately the same weight in each treatment. For growth assays involving transgenic lines, three independent transgenic lines were assayed for each construct and each line assayed in two or three independent biological experiments. A similar level of replication was performed for mutant lines. For other assays (proline, P5C, TUNEL, ROS, gene expression, enzyme activity), six-day-old seedlings were transferred to fresh control plates or -0.7 MPa low ψ_w_ treatment and both stress and control samples collected four days after transfer with exception of TUNEL assay where seedlings were collected nine days after transfer to the stress treatment.

### Proline, P5C, ROS and TUNEL assays

Proline was assayed following the ninhydrin method modified for use in 96 well plates (*59*). P5C was assayed using the O-amino-benzaldehyde (o-AB) assay (*54, 60*) with minor modifications. Samples (50 mg) were ground in liquid nitrogen and 600 µl 3% sulfosalicylic acid added followed by centrifugation at 15000xg for 10 min. Then, 300 µl supernatant was combined with 300 µl 10% trichloroacetic acid (TCA) and 100 µl 0.05 M o-AB added (o-AB was dissolved in 40% ethanol). Samples were incubated at room temperature for 25 min, centrifuged at 15000xg for 10 min and the 440 nm absorbance of the supernatant read using a spectrophotometer. P5C concentration was calculated using its molar absorption coefficient (2580 cm^-1^ M^-1^).

Levels of reactive oxygen species (ROS) were assayed using the fluorescent stain 2’, 7’-Dichlorodihydrofluorescein diacetate (DCFH-DA). For unstressed control seedlings, 10 μM DCFH-DA working solution was prepared from 10 mM DCFH-DA in DMSO stock solution. For stress treated seedlings, the 10 μM DCFH-DA working solution was prepared in in 285 mM mannitol, which is iso-osmotic with the -0.7 MPa PEG-agar plates and was used to prevent rehydration and swelling of the plant tissue during the assay. Seedlings were incubated in 10 μM DCFH-DA solution for 30 min followed by washing with water (for control seedlings) or 285 mM mannitol (for stress seedlings) for 1 min. Seedlings were mounted on the slide with water or 285 mM mannitol for control or stress-treated seedlings, respectively. The fluorescence signal from DCFH-DA was detected using a Leica Stellaris 8 microscope at 485 nm excitation and 535 nm emission wavelengths. Fluorescence intensity was quantified from at least 4-5 seedlings (control and stress treated) from each experiment. For quantification of ROS levels, a whole cell was selected as a region of interest (ROI) in Image J (using 16-bit images), and the mean gray value over the ROI determined using the analyze/measure function. Two or three independent experiments were conducted with consistent results in each experiment.

Terminal deoxynucleotidyl transferase dUTP nick end labeling (TUNEL) assay was performed following the protocol of Fendrych et al. (*61*) with minor modifications. Whole seedlings or excised roots were fixed by vacuum infiltration with 4% paraformaldehyde in PBS for 1 h at room temperature. The fixed tissue was washed five times with PBS, permeabilized by two-minute incubation with 0.1% Triton X-100 in 0.1% sodium citrate on ice and then again washed five times with PBS. Subsequent labelling steps followed the instructions of the One-step TUNEL in situ Apoptosis Kit-FITC (Elabscience, catalog number E-CK-A320). In brief, 100 µl of TdT Equilibration buffer was added to the sample and incubated at 37 °C for 30 min. After removal of TdT equilibration buffer, root tips were incubated with 50 µl of labelling working solution (35 µl TdT Equilibration buffer + 10 µl labelling solution + 5 µl TdT enzyme) at 37 °C for 1 h in darkness. Samples were then washed five times with PBS, nuclei stained with DAPI, washed five times with PBS and mounted onto slides with the antifading agent CutiFluorTM AF1 (Electron Microscopy Sciences 17970-25). FITC and DAPI fluorescence were then immediately imaged. To quantify TUNEL staining, FITC mean fluorescence intensity was quantified for five to eight arbitrarily selected nuclei in the root elongation zone to early maturation zone using ImageJ. Background intensity was measured in an immediately adjacent area was subtracted from the nuclear fluorescence intensity. The mean of the background fluorescence intensity for all the selected nuclei in one image was used to generate one data point. Three to five images were so analyzed per genotype and treatment.

### Mitochondria Isolation and Proline Dehydrogenase Enzyme Activity

ProDH activity of isolated mitochondria was measured by assaying isolated mitochondria using the 2,6-dichlorophenolindophenol (DCIP) reduction assay (*62*). To isolate crude mitochondrial fractions, five grams of seedlings were ground in a cold mortar with 20 ml of grinding buffer (0.3 M sucrose, 60 mM N-tris [hydroxymethyl]-methyl-2-aminoethanesulphonic acid (TES), 10 mM EDTA, 10 mM KH_2_PO_4_, 25 mM sodium pyrophosphate tetrabasic, 1 mM glycine, 1% (w/v) polyvinylpyrrolidone-40, 1% (w/v) defatted bovine serum albumin (BSA), adjusted to pH 8.0 with KOH; just prior to use 50 mM sodium ascorbate and 20 mM cysteine were added and pH readjusted to 8.0 with KOH). The extract was filtered through Miracloth and centrifuged at 2500xg for 5 min to remove most of the intact chloroplasts and thylakoid membranes. The supernatant was transferred to a new tube and centrifuged at 15000xg for 15 min (all centrifugations were performed at 4 °C). The resulting pellet was washed twice by suspending in wash buffer (0.3 M sucrose, 10 mM TES, 2 mM EDTA, 10 mM KH_2_PO_4_, pH 7.5) and centrifugation at 15000xg for 15 min with the supernatant discarded each time. The pellet was then resuspended in 1 ml wash buffer, divided into 100 µl aliquots, centrifuged at 15000xg and stored at -80 °C after removal of the supernatant.

ProDH activity was calculated from the rate of 2,6-dichlorophenolindophenol (DCPIP) reduction after addition of 150 mM proline (*63, 64*). Protein content of the crude mitochondrial fraction was measured by BCA assay (Pierce BCA Protein Assay Kit, Thermo Scientific catalog number 23225). Thirty µg of mitochondria protein was incubated at 25 °C in 850 µL reaction buffer (100 mM Tris-HCl, pH 7.5, 2.5 mM MgCl_2_, 1 mM KCN, 0.5 mM FAD, 0.5 mM phenazine methosulfate, 60 µM DCPIP) until a linear decrease of the OD600 was observed (typically 2–3 min). Then the reaction was started by adding 150 µL of a 1 M proline solution and the decrease in the OD600 monitored until at least 1 min of linear reaction was observed (DU-800 Spectrophotometer, Beckman Coulter). Reaction rates were calculated using the 600 nm absorbance coefficient of DCPIP (19.1 mM^-1^ cm^-1^) (*65*). The rate of DCPIP reduction was background corrected using blank reactions without mitochondrial protein but with addition of proline.

### Immunoblot detection of NRL5-YFP, P5CS1 and ProDH1

Samples (50-100 mg of seedlings from unstressed control or 4 day -0.7 MPa treatment) were ground in liquid N_2_ and 100 µL extraction buffer added (125 mM Tris-Cl pH 8.8, 1% SDS, 10% glycerol, 1 mM PMSF and Complete Protease Inhibitor [Roche]). Samples were centrifuged at 7000 g for 10 min and supernatant collected. Protein concentration was checked using Pierce BCA protein assay kit (Thermo Scientific, USA) and typically 100 µg protein was loaded onto 10% SDS-PAGE gels. Proteins were blotted onto PVDF membranes. For detection of NRL5-YFP, blots were probed with mouse anti-GFP antibody (Roche, #11814460001) used at 1/5000 dilution followed by rabbit Anti-Mouse HRP secondary antibody (AbCam ab6728) at 1/10,000 dilution. Detection of P5CS1 and ProDH1 used rabbit polyclonal antisera which were previously generated and characterized in our laboratory (*66, 67*). Both were used at 1/5000 dilution and followed by Goat Anti-Rabbit IgG H&L (HRP) secondary antibody (AbCam ab6721) used at 1/10000 dilution. Blots were developed with chemiluminescent substrate (Thermo Scientific) and exposed to film. For loading control, blots were stripped and re-probed with anti-HSC70 monoclonal antibody (Enzo #ADI-SPA-818-D) at 1/5000 dilution followed by Rabbit Anti-Mouse IgG H&L (HRP) secondary antibody (AbCam ab6728) at 1/10000 dilution. Blots were recorded on x-ray film and band intensities were quantified using the Fiji ImageJ package (http://fiji.sc/Fiji). An area of interest around the major band(s) in each lane was designated using the rectangular selection tool. The ‘plot lane’ option in the gel analyzer menu was used to obtain intensity peaks for each selected band. Peak areas for integration were marked, and peak areas calculated using the wand tool.

### Quantitative Reverse Transcriptase-PCR analysis of gene expression

Total RNA was extracted from control or -0.7 MPa stress-treated seedlings using RNeasy Plant Mini Kit (Qiagen). Total RNA (typically 1 μg) was reverse transcribed using SuperScript III (Invitrogen). Real-time quantitative PCR assays were prepared using KAPA SYBR FAST qPCR kit (Kapa Biosystems) and run on an Applied Biosystems QuantStudio 12K QPCR instrument. Gene expression values were calculated using the comparative cycle threshold method following normalization based on *EFL1α*, which has been demonstrated to be a reliable reference gene for abiotic stress studies (*68*). Amplification efficiency of each primer set was validated. Gene expression changes were quantified as log_2_ fold-change compared to the unstressed wild type. Primers used are given in Data S6.

### NRL5-YFP Immunoprecipitation and mass spectrometry protein identification

NRL5-YFP under control of the *35S* promoter was transiently expressed in Arabidopsis seedlings using procedures described below for rBiFC assays. At two days after infiltration, seedlings were transferred to -1.2 MPa low ψ_w_ stress on PEG-agar plates for 24 h. Seedlings were then sprayed with 50 μM MG132, or a mock control spray, and harvested three hours later. MG132 treatment was used based on past observation that some NPH3-domain proteins act as E3 ubiquitin ligase adaptors to target specific proteins for degradation and to generally enrich for proteins which may be unstable regardless of the underlying mechanism. Samples consisting of approximately five grams of tissue were homogenized in liquid nitrogen and extracted in lysis buffer consisting of 50 mM Tris (pH 7.5), 150 mM NaCl, 0.5% Triton X-100, 0.5 M EDTA and Protease Inhibitor (Roche). The cell lysate was collected by centrifuging the homogenate at 20,000g for 10 min. GFP-trap beads (GFPTrap-A kit, Chromotek) were equilibrated with dilution buffer (same as the extraction buffer except for the omission of Triton X-100). Equilibrated GFP-trap beads (20-30 μl) were added to the cell lysate and kept under constant mixing at 4°C for 2 h. Beads were collected by centrifugation at 2,500 g, washed one additional time with lysis buffer. Proteins were digested and peptides analyzed by LC-MS/MS on a Q-Exactive mass spectrometer. MS data were processed by Proteome discoverer and Mascot analysis (Mass spectrometry and peptide identification were conducted by the proteomics core facility of the Institute of Plant and Microbial Biology).

### Proteomic analysis of wild type and *nrl5-2*

Seven-day-old seedlings of wild type and *nrl5-2* were transferred to PEG-agar plates (-0.7 MPa) for stress treatment or to fresh control plates and samples collected four days after transfer. Four independent biological replicates were performed.

#### Protein digestion and TMT labeling

Protein digestion in the S-Trap micro column was performed according to the manufacture’s protocol. Briefly, 100 μg protein in 5% (v/v) SDS was reduced and alkylated using 10 mM TCEP and 40 mM CAA at 45°C for 15 min. A final concentration of 5.5% (v/v) PA followed by six-fold volume of binding buffer (90%, v/v, MeOH in 100 mM TEAB) was next added to the protein solution. After gentle vortexing, the solution was loaded into an S-Trap micro column. The solution was removed by spinning the column at 4,000 g for 1 min. The column was washed with 150 μl binding buffer three times. Finally, 20 μl of digestion solution (1 unit Lys-C and 1 μg trypsin in 50 mM TEAB) was added to the column and incubated at 47 °C for 2 h. Each digested peptide was eluted using 40 μl of three buffers consecutively: (1) 50 mM TEAB, (2) 0.2% (v/v) FA in H_2_O, and (3) 50% (v/v) ACN (standard elution). Elution solutions were collected in a tube and dried under vacuum. Peptides from each sample were desalted using an Oasis HLB cartridge (Waters). Desalted peptides (100 μg) from each sample were labeled with one of the TMT-10plex reagents (0.8 mg, Thermo Fisher Scientific) according to the manufacture’s protocol. The labeling reactions were quenched by adding 5% hydroxylamine for 15 min and then acidified with 10% trifluoracetic acid. The labeled peptides with different TMT tags were mixed in the same tube and then dried with a SpeedVac.

#### High pH reversed-phase fractionation

The dried peptides were suspended in 400 μl of Buffer A (200 mM ammonium formate with 2% (v/v) acetonitrile (ACN)). The peptide solution was loaded into a high-pressure liquid chromatography (Agilent 1200 series) with an autosampler (GILSON FC 203B). Buffer B was 200 mM ammonium formate with 90% (v/v) ACN. Peptides were separated by a gradient mixture with an XBridge BEH C18 column (Waters) (4.6 mm X 250mm). Peptides were separated by a gradient: 0% buffer B in 7 min, 0-16% buffer B in 6 min, 16-40% buffer B in 60 min, 40%-44% buffer B in 4 min, 44%-60% buffer B in 5 min, 60% buffer B in 6 min at 1 mL/min. A total of 80 fractions were collected and concatenated into 24 fractions. Fractionated peptides were dried with a SpeedVac and stored at −80 °C until LC-MS/MS analysis.

#### LC-MS/MS and Data analysis

Dried peptides were suspended in 0.1% formic acid (FA) with 2% ACN and analyzed by LC-MS/MS using an Orbitrap Fusion Lumos mass spectrometer (Thermo Scientific) connected to an EASY nLC 1200 (buffer A: 0.1% FA with 3% ACN and buffer B: 0.1% FA in 90% ACN) in a 2 h LC gradient. Peptides were separated on a 25 cm Thermo Acclaim PepMap RSLC column with a column heater set at 45 °C. The mass spectrometer was operated in the data-dependent acquisition mode, in which a full MS scan (m/z 350–1,800; resolution: 120,000) was followed by the most intense ions being subjected to higher-energy collision-induced dissociation (HCD) fragmentation within 3 s. Data were acquired in the Orbitrap with a resolution of 50,000 [normalized collision energy (NCE): 35%; AGC target: 5e4, max injection time: 100 ms; isolation window: 0.7 m/z; dynamic exclusion: 15 s]. The raw files were searched using Proteome Discoverer (version 2.5) against an Arabidopsis TAIR10 database (35,387 entries). The precursor mass tolerance was set to 10 ppm, and the fragment mass tolerance was set at 0.02 Da. The peptide search was performed using tryptic/P digestion and allowed two miss cleavages. Carbamidomethyl (C), TMT10plex (peptide N-term), and TMT10plex (K) were set as a fixed modification, and acetylation (protein N-term) and oxidation (M) were set as variable modifications. Strict protein and peptide false discovery rates (FDR) were set for 0.01. Proteins and peptides were quantified from the TMT10plex reporter ions at MS2 level by using the reporter ion quantifier node in the Proteome Discoverer.

For proteins identified in at least two of the four replicates, a mean log_2_ fold change ≥ 0.5 and P ≤ 0.05 (one sample T-test) were used as the criteria to define differentially abundant proteins.

### Recombinant protein production, by *E. coli* expression or *in vitro* translation, and purification

For production of recombinant protein with an N-terminal 6x-His tag in *E coli*, the coding sequences of NRL5, NRL5^Δ1-115^ (BTB domain removed), NRL5^P335L^, NRL5^P335LΔ1-115^ and VAMP721^ΔTM^ were amplified by PCR and cloned into the expression vector pET28a. For production of recombinant protein with an N-terminal GST tag, NRL5, NRL5^Δ1-115^, NRL5^P335L^, NRL5^P335LΔ1-115^, RABH1b, RABE1c and VAMP721^ΔTM^ (transmembrane domain removed), VAMP721-SD (SNARE domain), VAMP721-LD (LONGIN domain) and VAMP722^ΔTM^ coding sequences were amplified by PCR and cloned into GST expression vector pGEX-4T1. Primers used for cloning are listed in Data S6. HIS- and GST-tagged proteins were produced in *E.Coli* BL21 Rosetta strain. Recombinant protein production was induced by addition of 1mM IPTG to log phase cultures (OD_600_ 0.4 to 0.5) and incubated overnight at 18 °C. Cells were harvested by centrifugation at 6,000xg for 30 min at 4 °C. Bacterial cell pellets were washed by resuspending in 50 μl ice-cold PBS per ml culture followed by centrifugation. For extraction of GST-tagged proteins, cell pellets were resuspended in ice-cold binding buffer consisting of 50 mM Tris-Cl (pH 8.0), 150 mM NaCl, 10% v/v glycerol, 5 mM 2-mercaptoethanol, 1% Triton X-100, 1 mM DTT, 1 mM EDTA and 1X protease inhibitor. For extraction of His-tag protein, cell pellets were resuspended in binding buffer consisting of 50 mM Tris-Cl (pH 8), 5 mM imidazole, 150 mM NaCl, 0.1 mM EDTA and 1 mM PMSF (added just before use). Bacterial cells were lysed using a French press high pressure cell disruptor followed by centrifugation at 40,000xg for 1 h at 4 °C. The supernatant was further clarified using a 0.22 micron filter and incubated with washed Glutathione or Ni-affinity beads for 2 h at 4 °C with gentle agitation [GE-Glutathione Sepharose High Performance (catalog #17-5279-0) or GE-Ni Sepharose High Performance (17-5268-01)]. The beads were then washed three times with 10 volumes of binding buffer (50 mM Tris-Cl [pH 8.0], 1 M NaCl, 10% v/v glycerol, 5 mM 2-mercaptoethanol, 1 % Triton X-100, 1 mM DTT, 1 mM EDTA and 1X protease inhibitor). GST-tagged protein was eluted with buffer containing 50 mM Tris-Cl, pH 8.0, 100 mM NaCl, 0.1% Triton X-100, 50 mM Reduced glutathione, 1 mM DTT and 1x protease inhibitor. For His-tagged proteins beads were washed three time with wash buffer (50 mM Tris-Cl pH 8.0, 1 M NaCl, 100 mM imidazole, 10% [v/v] glycerol, 5 mM 2-mercaptoethanol, 1 mM PMSF) and proteins were eluted with 50 mM Tris-Cl, 200 mM NaCl, 500 mM Imidazole, 0.1 mM EDTA and 1 mM PMSF. Proteins were dialyzed against 50 mM Tris-HCl, 50 mM NaCl with buffer exchanged three times at three-hour intervals. Protein was concentrated using 30 kDa cutoff centrifuge filter apparatus (VivaSpin 30 kDa). Protein purity was checked by SDS-PAGE and Commassie staining.

In vitro translation was performed with the 1-Step High Yield IVT Kit (Thermo-Fischer Scientific, Catalog No. 88891), which uses human cell extracts as the basis for protein production, following the manufacturer’s instructions. Recombinant protein with N-terminal 9x-His and GST tag was produced by amplifying the coding sequences of NRL5 and NRL5^P335L^ and cloning into the expression vector pT7CFE1-NHIS-GST-CHA (Human In Vitro Protein Expression Vector, ThermoFisher Scientific, product number: 88871, Lot number: YB366542). Recombinant protein was produced following the procedure for the 100 μl reaction, as per product instructions. The completed reaction was centrifuged for 1 min at 10,000xg. The clarified sample was diluted 5-fold in glutathione binding buffer (125 mM Tris, pH 8.0, 150 mM NaCl, 1% Triton^TM^ X-100, 10% glycerol, 1 mM DTT) and added to 500 μl of Glutathione Sepharose^TM^ High-Performance resin (Cytiva, Lot number: 10316359) in an eppendorf tube and incubated overnight at 4 °C with gentle mixing. The resin was then loaded into PD-10 column and was washed three times with 5 ml glutathione wash buffer (125 mM Tris, pH 8.0, 1M NaCl, 1% Triton^TM^ X-100, 10% glycerol, 1 mM DTT) flow-through collected by gravity. GST-tagged protein was eluted with 200 μl glutathione elution buffer (125 mM Tris, pH 8.0, 1M NaCl, 1% Triton^TM^ X-100, 10% glycerol, 1 mM DTT, 50 mM glutathione) by gravity flow. Samples were then dialyzed against 50 mM Tris-HCl pH 8.0, 50 mM NaCl with three times buffer exchange at three-hour intervals. Protein was concentrated using 50 kDa cutoff centrifuge filter (VivaSpin 50 KDa). Protein purity was assayed SDS-PAGE and silver staining (for NRL5 and NRL5P335L produced by in vitro translation) or by Commassie Blue staining (for proteins produced in *E. coli*).

For silver staining, SDS PAGE gels were incubated in fixing solution (50 % methanol, 10 % glacial acetic acid, 0.05% formalin) for 2 hours with gentle agitation and then washed in 35 % ethanol three times for 20 minutes each time. This was followed by a two-minute incubation in 0.02% Na_2_S_2_O_3_ and three five-minute washes in water. Staining was performed by 20 min of gentle agitation in staining solution (0.2% AgNO_3_, 0.076% formalin) followed by two water washes (1 min each). The gel was then incubated in developing solution (6% Na_2_CO_3_, 0.05% formalin, 0.0004% Na_2_SO_3_) followed by five minutes in stop solution (50% methanol, 10% glacial acetic acid) with solution changes as needed to remove excess stain and a further set of three five-minute washes in water. Silver stain gel images were captured using the Image Lab software of a BIO-RAD molecular imaging Gel Doc^TM^ XR^+^ Imaging System.

### In-vitro Pull down assay

Each pull-down reaction used 20 μl of magnetic GSH beads (Thermo-Pierce Glutathione Magnetic Agarose Beads, catalog #78602). Beads were prepared by washing three times with 1 ml PBS followed by three washes with 1 ml binding buffer (20 mM Tris-HCl pH 7.5, 150 mM NaCl, 5 mM MgCl_2_, 10% v/v glycerol, 0.05% Tween 20, 0.1mM EDTA). For assays involving RAB proteins, the pull-down reactions were assembled by adding 1.5 nmol of GST or GST fusion protein, 5 μl of 0.5 M EDTA (pH 8.0), and 5 μl of 10 mM GTPγS or GDP to the washed beads. Volume was adjusted to 600 μl by adding binding buffer and beads incubated at 4 °C with gentle mixing for 2 h followed by addition of 5 μl of 1 M MgCl_2_ and an additional five-minute incubation. His-tagged protein (0.5 nmol) was then added followed by and 5 μl of 1 mM GTPγS or GDP. Volume was adjusted to 600 μl by adding binding buffer and incubated at 4°C for 3 h with gentle mixing.

Beads were then washed six times with 1 ml binding buffer containing 0.25% IGEPAL. Beads were boiled with 2X SDS-PAGE loading dye, proteins separated on 10% SDS-PAGE gels and transferred to PVDF membrane (BIO-RAD, catalog #1620177) for immunoblotting. His- and GST-tagged proteins were detected with anti-His (Monoclonal Anti-polyHistidine antibody, Sigma H1029, 1:5000 dilution) or anti-GST (Mouse anti-GST monoclonal antibody, Sigma SAB4200692, 1:5000 dilution), respectively, followed by (Rabbit Anti-Mouse IgG H&L [HRP] secondary antibody, AbCam ab6728). For detection of input protein amount, five percent of the 1.5 nmol GST tagged protein and twenty percent of the 0.5 nmol His tagged protein were used.

For NRL5 self-interaction and interaction with VAMP721/722, 20 μl of magnetic GSH beads were washed 3 times with 1 ml PBS and equilibrated by washing at least three times with protein purification buffer (50 mM Potassium phosphate (pH 7.5), 150 mM NaCl, 10% v/v glycerol and 0.1 mM EDTA). Binding reactions were assembled by adding 1.5 nmol of GST-tagged protein, adjusting the volume to 600 μl with purification buffer and pre-incubating at 4 °C with gentle mixing for 2 h. The beads were then washed three times with 1 ml purification buffer and 0.5 nmol of HIS-tagged protein added in total volume of 600 μl followed by incubation at 4 °C for 3 h with gentle mixing. Beads were then washed six times with 1 ml purification buffer containing 0.25% IGEPAL, boiled in 2x SDS-PAGE loading dye and immunoblotting and protein detection performed as described above. Immunoblot band intensities were quantified using the Fiji ImageJ package (http://fiji.sc/Fiji) and tif image files generated by UVP ChemStudio Plus gel imaging system using Vision Works software. An area of interest around the major band(s) in each lane was designated using the rectangular selection tool. The ‘plot lane’ option in the gel analyzer menu was used to obtain intensity peaks for each selected band. Peak areas for integration were marked, and peak areas calculated using the wand tool.

### Ratiometric Bi-molecular Fluorescence Complementation (rBiFC)

Ratiometric BiFC (rBiFC) assay was performed using previously described vectors (*69*) where a constitutively expressed RFP is present on the BiFC plasmid and used to normalize the BiFC fluorescence signal. Coding sequences of the proteins to be analyzed were cloned into pDONR221 P1-P4 and pDONR221 P3-P2 (Invitrogen) and used for LR reaction with the destination vector pBiFCt2in1-NN (for NRL5 or RPT2 interaction with RABs and VAMP721 or NRL5/RPT2 self-interaction) or with pBiFCt2in1-CC (for NRL5/RPT2 self-interaction). The resulting plasmids were transformed into *Agrobacterium tumefaciens* strain GV3101. A single colony was picked and used to inoculate 10 ml of LB medium prepared with specific antibiotics. This primary culture was incubated at 28 °C with shaking (180-200 rpm) for 16-18 h and then sub-cultured into 150-300 ml of LB medium with specific antibiotics and incubated at 28 °C with shaking (180-200 rpm) for 24-30 hr. Cells were collected by centrifugation at 3,500 rpm for 10 minutes at 4 °C and the pellet resuspended in infiltration medium (5% sucrose, 5 mM MES, 200 μM Acetosyringone, 20-25 ml infiltration medium used for 150 ml *A. tumefaciens* culture). Transient expression was performed using an Arabidopsis line with Dexamethasone-inducible AvrPto expression (Arabidopis Biological Research Center CS67140) (*70*). Four-day-old AvrPto seedlings were sprayed with 10 μM Dexamehtasone to induce AvrPto expression and seedlings were covered with 20-25 ml of *A. tumefaciens* cell suspension and vacuum infiltrated twice for 1 minute at 10 mm Hg. After infiltration, seedlings were allowed to recover for 24 h on control agar media and then transferred to fresh control (-0.25 MPa) or PEG-infused agar plates (-0.7 MPa) and imaging conducted 48 h after transfer. To visualize the rBiFC interaction, YFP was detected using excitation wavelength 514 nm with a emission filter of 520-555 nm while RFP was detected using excitation wavelength 543 nm and 560-615 nm emission filter. Imaging was performed on a Zeiss LSM 510 Meta or Leica Stellaris 8 microscope. YFP/RFP fluorescence intensity ratios were quantified for typically 20 individual cells from 4-5 seedlings combined from 2 or 3 three independent experiments (see figure legends for details of replication and n values for each rBiFC interaction assay). For each quantification, a whole cell was selected as a region of interest in Image J, and the mean intensity of RFP and YFP determined using the analyze/measure function.

### GTP hydrolysis Assay

GTPase activity was quantified using malachite green detection of inorganic phosphate (P_i_) release. Malachite green detection solution was prepared freshly by mixing stock solutions of Malachite green dye (0.1% [w/v] malachite green, 6 N sulphuric acid), 7.5% (w/v) ammonium molybdate and 11% (v/v) Tween 20 in a 10:2.5:0.2 ratio. For assay of GTPase activity, a 20 µl reaction was set up using either 1 µM or 0.25 µM of purified protein (or combinations of proteins, as described in text and figures) in reaction buffer [50 mM Tris (pH 7.5), 50 mM NaCl, 5 mM MgCl_2_, 10 mM β-mercaptoethanol]. Reactions were started by addition of substrate (GTP, GTPγS, GDP or ATP) at concentration of 0 to 32 mM (as indicated in figures and figure legends), mixed and incubated for 30 min at 30 °C. Reactions were stopped by dilution to 180 µl (using ultra-pure water), addition of 50 µl malachite green reaction solution and incubation for 2 min at room temperature. Then, 20 µl of 34% (w/v) sodium citrate was added followed by incubation for further 30 min at 30 °C and 620 nm absorbance measured using a plate reader (BioTek microQuant). Data were background corrected by subtracting the absorbance values of mock control reactions (buffer only and buffer plus different GTP concentrations) and further corrected for background of sample with protein but without GTP added to account for background levels of Pi. The molar amount of Pi released was then calculated from standard curves prepared using 1 mM KH_2_PO_4_.

### Desthiobiotin-GTP protein labelling and LC-MS identification of biotinylated peptides

Desthiobiotin-GTP protein labeling was conducted Pierce GTPase ActivX Probe (Thermo Scientific, catalog #88315). For each labelling reaction, 0.5 μM of purified protein was prepared in 25 μl reaction buffer (50 mM Tris-HCl pH 8, 50 mM NaCl, 10 mM β-mercaptoethanol) with 0.25 μl 0.1 M EDTA added and the mixture incubated for 5 minutes at room temperature. This was followed by addition of 0.5 μl of manufacture-supplied desthiobiotin-GTP solution (to give 20 μM final concentration) followed by mixing of samples and addition of 0.5 μl of 1 M MgCl_2_ to start the reaction. For negative control reactions, MgCl_2_ was omitted. Reactions were incubated for 2 or 10 minutes at room temperature followed by addition of 5x Laemmli reducing sample loading buffer and heating at 95 °C for 5 minutes. Proteins were separated by SDS-PAGE and biotinylated proteins detected using Streptavidin-HRP (Sigma # RABHRP3) and Supersignal detection kit (Thermofisher Scientific). Blot images were acquired with a UVP ChemStudio Plus gel imaging system using Vision Works software.

For LC-MS analysis to identify biotin-labeled peptides, the amount of protein in the labeling reaction was increased to 1 μg. Following protein digestion (as described above in the proteomics section), biotinylated peptides were isolated using streptavidin beads following the manufacturer’s protocols for the desthiobiotin reagent. Peptides were then loaded onto an Evotips Pure column and peptides identified by an Evosep One LC system (EvoSep) coupled to a timsTOF HT mass spectrometer (Bruker). The mass spectrometer was operated using the Extended method (longer gradient time for peptide separation) as specified by the manufacturer and was used with the EV1137 performance column (15 cm X 150 µm ID, 1.5µm, EvpSep) at 40 °C inside a Captive spray source (Bruker).

The mass spectrometer was operated in DDA-PASEF mode with ten PASEF/MSMS scans per topN acquisition cycle. All spectra were acquired within a m/z range of 100 to 1,700 and IM of 0.75 to 1.45 Vs cm-2, the accumulation and ramp time was specified as 100ms, capillary voltage was set to 1400 V and the collision energy was a linear ramp from 20 eV at 1/K0 = 0.6 Vs cm-2 to 59 eV at 1/K0 = 1.6 Vs cm-2. Singly charged precursors were excluded by their position in the m/z-IM plane using a polygon shape, and precursor signals over an intensity threshold of 2,500 were picked for fragmentation. Precursors were isolated with a 2 Th window below m/z 700 and 3 Th window above m/z 800, as well as actively excluded for 0.4 min when reaching a target intensity of 20,000. All the MS raw files were searched using SpectroMine software (version 4.0) against TAIR10 database. The fixed modification was set as carbamidomethyl (C), and variable modifications were set as oxidation (M), acetylation (protein N-term), and desthiobiotinylation (K). The FDR for peptide-spectrum match (PSM), ion, peptide, and protein level was set at 1%.

### NRL5 subcellular localization and co-localization with membrane markers as well as with RABE1c and VAMP721

For imaging of NRL5 subcellular localization, seedlings of T_3_ homozygous transgenic lines were observed either under control condition or four days after transfer to PEG-infused agar plates ( -0.7 MPa). Cell walls were stained with Propidium Iodide (10 μg/ml) in water for control plants and in 285 mM mannitol for stress plants (to maintain the water potential of -0.7 MPa and prevent cell swelling) for 60 s (control) or 90 s (stress) and rinsed with distilled water or mannitol solution for 1 min to remove excess PI from the tissue before mounting on glass slide for confocal microscopy. For colocalization with intracellular markers, homozygous T_3_ lines of *NRL5_pro_:NRL5-YFP* and *NRL_pro_:NRL5^P335L^-YFP* were crossed to homozygous T_3_ lines that had been transformed with vectors for plasma membrane marker (pCMU-PMr) or plasmodesmata marker (pCMB-PDESr) previously described by Ivanov et al. (*71*) and obtained from Addgene. The subcellular markers were expressed under control of ubiquitin promoter and were tagged with mCherry. YFP was detected using excitation emission wavelength of 514/542 nm and mCherry was detected using excitation emission wavelength of 587/610 nm on a Leica Stellaris 8 with 40X or 63X water immersion lenses. PI was detected using excitation emission wavelength of 543/610 nm. To analyze the effect of RABE1c and RABH1b on NRL5 localization, the same homozygous T_3_ line of *NRL5_pro_:NRL5-YFP* was crossed to the *rabe1c* and *rabh1b* T-DNA mutants and plants homozygous for the T-DNA mutation and containing the *NRL5_pro_:NRL5-YFP* construct isolated by PCR genotyping of F_2_ plants. F_3_ plants were screened to find lines where the *NRL5_pro_:NRL5-YFP* was no longer segregating and these lines used for further analysis of NRL5 localization.

For NRL5-RABE1c-VAMP721 colocalization, the *NRL5_pro_:NRL5-CFP* construct described above was transformed into *35S:YFP-RABE1c* T_3_ homozygous transgenic line and screened by antibiotic selection to obtain T_3_ homozygous lines for the NRL5 construct. These *NRL5_pro_:NRL5-CFP/35S:YFP-RABE1c* T_3_ lines were then crossed to a T_3_ homozygous line expressing *35S:mCherry-VAMP721*. F_2_ plants containing all three constructs were selected and resulting F_3_ seed was used for colocalization analysis. Seedlings from such a line expressing *NRL5_pro_:NRL5-YFP, 35s:YFPRABE1c* and *35S:mCherry-VAMP721* were observed either under control condition or four days after transfer to PEG-infused agar plates (-0.7 MPa). CFP was detected using excitation emission wavelength of 433/475 nm, YFP was detected using excitation emission wavelength 514/542 nm and mCherry was detected using excitation emission wavelength of 587/610 nm on a Leica Stellaris 8 with 40X water immersion lens. Time lapse movies of *NRL5_pro_:NRL5-YFP/35s:YFP-RABE1c/35S:mCherry-VAMP721* was taken by Leica Stellaris 8 microscope with 40X, 4x zoom water immersion lens. Time lapse movies were processed using ImageJ and exported at 2 frames per second. Movies were annotated using ImageJ Fiji software. For treatment with Brefeldin A (BFA), five-day-old seedlings were immersed in 50 μM BFA (diluted from 50 mM stock solution) or equal concentration of DMSO for 2 hours. Treated seedlings were rinsed with distilled water before mounting on glass slide.

Co-localization was quantified using the Pearson’s Correlation Coefficient (PCC). A region of interest, typically the bottom edge of hypocotyl epidermal cells or selected edge of leaf pavement cells, was selected and PCC calculated using Coloc2 in Fiji ImageJ.

### Structural Models and Sequence Alignment

A structural model of NRL5 was downloaded from the AlphaFold Protein Structure Database (*72, 73*). Sequence alignment of NPH3 domain proteins was performed using Clustal Omega (*74*).

### Statistical Analyses

Data were analyzed by ANOVA. In cases where the normality assumption was not upheld, non-parametric Kruskal-Wallis test or one sample Wilcoxon test were used, as described in the text or figure legends. Statistical analyses were performed using GraphPad Prism 9 or 10.

## Supporting information

Movie S1 NRL5-CFP and YFP-RABE1c control

Movie S2 NRL5-CFP and mCherry-VAMP721 control

Movie S3 YFP-RABE1c and mCherry-VAMP721 control

Movie S4 NRL5-RABE1c-VAMP721 control

Movie S5 NRL5-CFP and YFP-RABE1c stress

Movie S6 NRL5-CFP and mCherry-VAMP721 stress

Movie S7 YFP-RABE1c and mCherry-VAMP721 stress

Movie S8 NRL5-RABE1c-VAMP721 stress

## Acknowledgments

We thank S. Shinde and S. Mukiri for assistance in the early stage of NRL5 mutant characterization and complementation crosses; A. Savouré (Sorbonne Université) for advice on ProDH activity assay; J.-Y. Huang and M.-J. Feng for assistance with confocal microscopy and image quantitation, H.-Y. Chang for laboratory assistance and O.-K. Teh for critical reading of the manuscript.

## Funding

Academia Sinica Investigator Award AS-IA-108-L04 (PEV)

National Science and Technology Council, Taiwan NSTC 109-2311-B001-019-MY3 (PEV)

Academia Sinica, Institute of Plant and Microbial Biology (PEV)

National Science and Technology Council, Taiwan NSTC 110-2311-B-001-043-MY2 (C-CH)

Academia Sinica Core Facility and Innovative Instrument Project AS-CFII-111-209 (C-CH)

## Author contributions

Conceptualization: PEV

Methodology: PEV, NU-T, Y-CL, C-CH

Investigation: NU-T, X-JH, Y-CL, SKS, S-SH

Visualization: PEV, NU-T

Funding acquisition: PEV, C-CH

Project administration: PEV, S-SH, C-CH

Supervision: PEV, C-CH

Writing – original draft: PEV, NU-T, X-JH

Writing – review & editing: PEV, NU-T, X-JH, C-CH, Y-CL

## Competing interests

Authors declare that they have no competing interests.

## Data and materials availability

For proteomic analysis of wild type and *nrl5-2*, MS raw data and Proteome Discoverer output files have been deposited with the ProteomeXchange Consortium via the jPOST repository with the data set identifier PXD040396. (Data available for review at URL: https://repository.jpostdb.org/preview/68146114463f9c7cda5626 with the code: 9538). NRL5 protein structure was obtained from AlphaFold Protein Structure Database (AlphaFold Protein Structure Database (ebi.ac.uk). Gene and protein sequence information used in this study can be obtained from The Arabidopsis Information Resource [TAIR, TAIR - Home Page (arabidopsis.org)] using accession numbers listed in the article or supplementary material. Other data are available in the main text or the supplementary material.

## Supplementary Materials

Figs. S1 to S21

Data S1 to S6

Supplementary Movies S1 to S8

## Notes

### Competing Interest Statement

The authors have declared no competing interest.

### Summary of Updates

Substantial new data has been added regarding GTPase activity of NRL5 and other NPH3-domain proteins as well as NRL5 co-localization with RABE1c and VAMP721. Text and figures have been revised and reformatted.

