## Supplemental Figures S1-S21 for "The Non-Phototrophic Hypocotyl3 (NPH3)-domain protein NRL5 is a trafficking-associated GTPase essential for drought resistance"

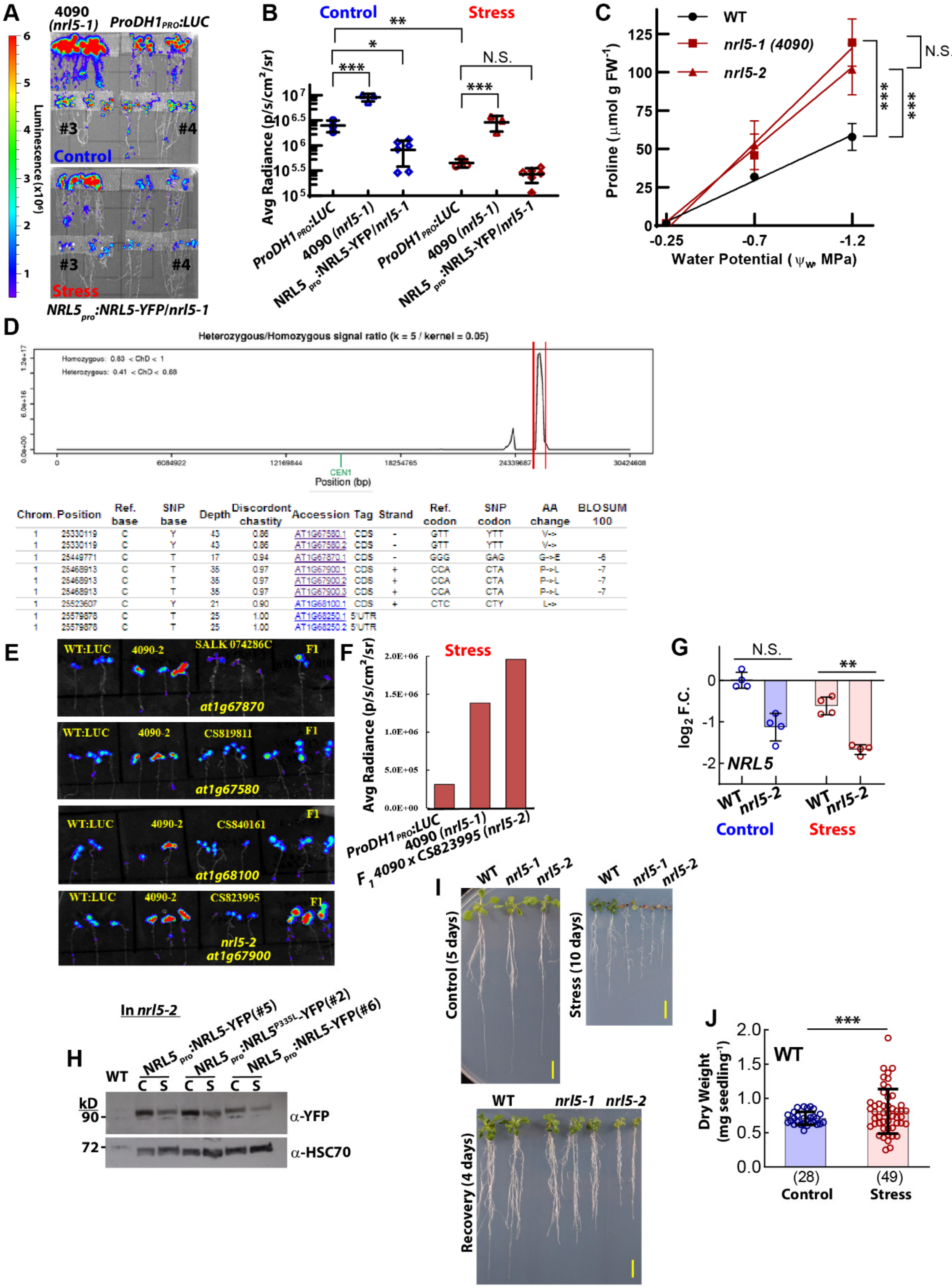


| Chrom. | Position | Ref. base | SNP base | Depth | Discordont chastity | Accession | Tag | Strand | Ref. codon | SNP codon | AA change | BLOSUM 100 |
|---|---|---|---|---|---|---|---|---|---|---|---|---|
| 1 | 25330119 | C | Y | 43 | 0.86 | AT1G67580.1 | CDS | - | GTT | YTT | V-> | |
| 1 | 25330119 | C | Y | 43 | 0.86 | AT1G67580.2 | CDS | - | GTT | YTT | V-> | |
| 1 | 25449771 | C | T | 17 | 0.94 | AT1G67870.1 | CDS | - | GGG | GAG | G->E | -6 |
| 1 | 25468913 | C | T | 35 | 0.97 | AT1G67900.1 | CDS | + | CCA | CTA | P->L | -7 |
| 1 | 25468913 | C | T | 35 | 0.97 | AT1G67900.2 | CDS | + | CCA | CTA | P->L | -7 |
| 1 | 25468913 | C | T | 35 | 0.97 | AT1G67900.3 | CDS | + | CCA | CTA | P->L | -7 |
| 1 | 25523607 | C | Y | 21 | 0.90 | AT1G68100.1 | CDS | + | CTC | CTY | L-> | |
| 1 | 25579878 | C | T | 25 | 1.00 | AT1G68250.1 | 5'UTR | | | | | |
| 1 | 25579878 | C | T | 25 | 1.00 | AT1G68250.2 | 5'UTR | | | | | |

**Fig. S1: Mutation of *NRL5* leads to high *ProDH1$_{pro}$:LUC* reporter expression and extreme sensitivity to moderate severity low $\psi_w$.**
**(Legend on next page)**

**Fig. S1: Mutation of *NRL5* leads to high *ProDH1$_{pro}$:LUC* reporter expression and extreme sensitivity to moderate severity low $\psi_w$.**
(**A**) Representative false color images of luciferase activity in the *ProDH1$_{pro}$:LUC* line used for mutant screening, *nrl5-1*, and two independent transgenic lines of *nrl5-1* complemented with NRL5-YFP expressed under control of the *NRL5* native promoter. (**B**) Quantitation of luciferase activity. Seedlings were imaged 4 days after transfer to control (-0.25 MPa) or low $\psi_w$ (-0.7 MPa) plates. Data are means ± S.D. (n = 6) with each data point representing a group of 4-6 seedlings grown in the same plate. Data were combined from two independent experiments and analyzed by ANOVA. (**C**) Proline levels of wild type and *nrl5* mutants across a range of $\psi_w$. Data are means ± S.D. (n = 3) with each data point from an independent experiment (three biological replicates assayed per experiment). Data were analyzed by linear regression analysis and analyzed for significant differences in slope of the indicated regression lines. (**D**) Output of Next Generation Mapping software showing a single peak of homozygous SNPs on chromosome 1 as well as candidate genes and SNPs within the association interval. (**E**) Genetic complementation tests using mutant number *4090* (*nrl5-1*) and candidate gene T-DNA lines indicated that mutation of *NRL5* caused the elevated LUC activity of *nrl5-1*. Luciferase imaging of parental lines and $F_1$ seedlings for each cross after stress treatment is shown. (**F**) Quantitation of luciferase intensity for *4090* (*nrl5-1*) and $F_1$ of *nrl5-1* crossed to the *nrl5-2* T-DNA mutant. *ProDH1$_{pro}$:LUC* is the un-mutagenized parental line used to isolate the *nrl5-1* mutant. (**G**) *NRL5* expression was reduced by low $\psi_w$ stress (-0.7 MPa for 4 days). *nrl5-2* (SAIL_565_C10/ CS823995) had further reduced level of *NRL5* mRNA consistent with the insertion of the T-DNA in the *NRL5* promoter or 5' UTR (depending on alternative transcriptional start sites annotated for *NRL5*). Data are means ± S.D (n =4). and were analyzed by ANOVA. (**H**) Immunoblot of representative transgenic lines where NRL5 or NRL5$^{P335L}$ with C-terminal fusion to YFP was expressed under control of the *NRL5* native promoter in the *nrl5-2* background. The blot was re-probed with anti-HSC70 as a loading control. NRL5 and NRL$^{P335L}$ proteins accumulated to similar levels and both were reduced in abundance during low $\psi_w$ stress. Note that for each construct, at least three independent lines were phenotyped and examined for NRL5-YFP or NRL5$^{P335L}$-YFP protein level using confocal microscopy and immunoblotting with consistent results among the independent transgenic lines (see Fig. S14 for representative images of NRL5-YFP and NRL5$^{P335L}$-YFP localization). (**I**) Representative images of wild type, *nrl5-1* and *nrl5-2* in the unstressed control (five-day-old seedlings transferred to fresh control media for five days), low $\psi_w$ stress (five-day-old seedlings transferred to -0.7 MPa plates for 10 days) and recovery (seedlings exposed to -0.7 MPa, then returned to control media for four days). Scale bars indicate 1 cm. (**J**) Representative dry weights of Col-0 wild type (WT) seedlings used for normalizing mutant and transgenic data shown in Fig. 1. Five-day-old seedlings were transferred from unstressed agar plates ($\psi_w$ = -0.25 MPa) to moderate severity low $\psi_w$ stress plates (-0.7 MPa). Dry weights were measured 5 days after transfer for the unstressed control and 10 days after transfer for the low $\psi_w$ treatment. This allowed both stress and control seedlings to reach a similar dry weight (although note that root elongation and rosette leaf area was still reduced in the stress treatment despite the longer time). Data are means ± S.D. combined from 2 representative experiments analyzed by T-test. Error bars show S.D.
For statistical analyses *, **, *** and N.S. indicate P ≤ 0.05, P ≤ 0.01, P ≤ 0.001 and non-significant differences, respectively.

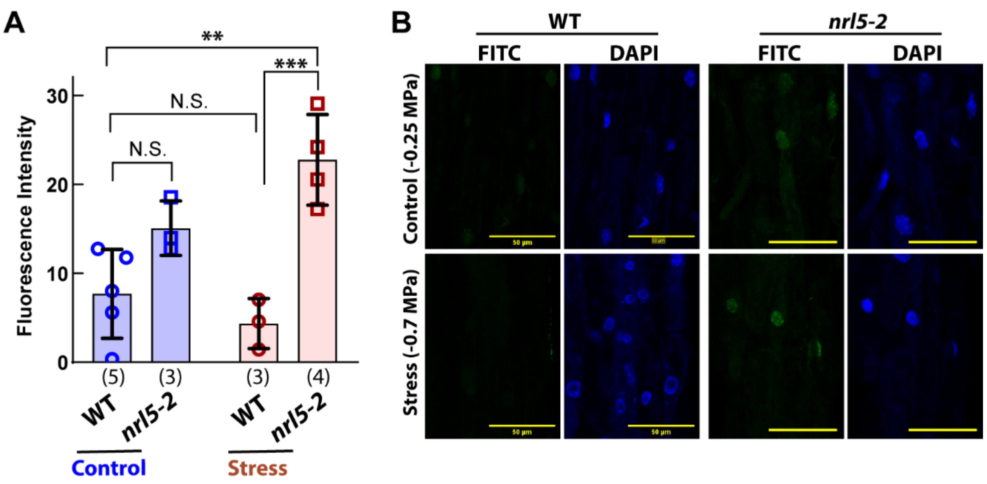


**Fig. S2: *nrl5-2* has increased DNA damage at low $\psi_w$.**
(**A**) Tunnel assay found that *nrl5-2* had increased level of DNA damage at low $\psi_w$. Seedlings were assayed at 9 days after transfer to low $\psi_w$ (-0.7 MPa) or 4 days after transfer to unstressed control plates. Fluorescence intensity was quantified for individual nuclei from elongating cells in the primary root. Eight to ten nuclei were quantified per root and each data point shown represents one root. Data are means ± S.D. and were analyzed by one-way ANOVA (n values shown in parentheses). (**B**) Representative images of FITC staining (to mark DNA damage) and DAPI staining (to mark position of nuclei). Scale bars indicate 50 µm.
For statistical analyses **, *** and N.S. represent $P \leq 0.01$, $P \leq 0.001$ and not significant, respectively.

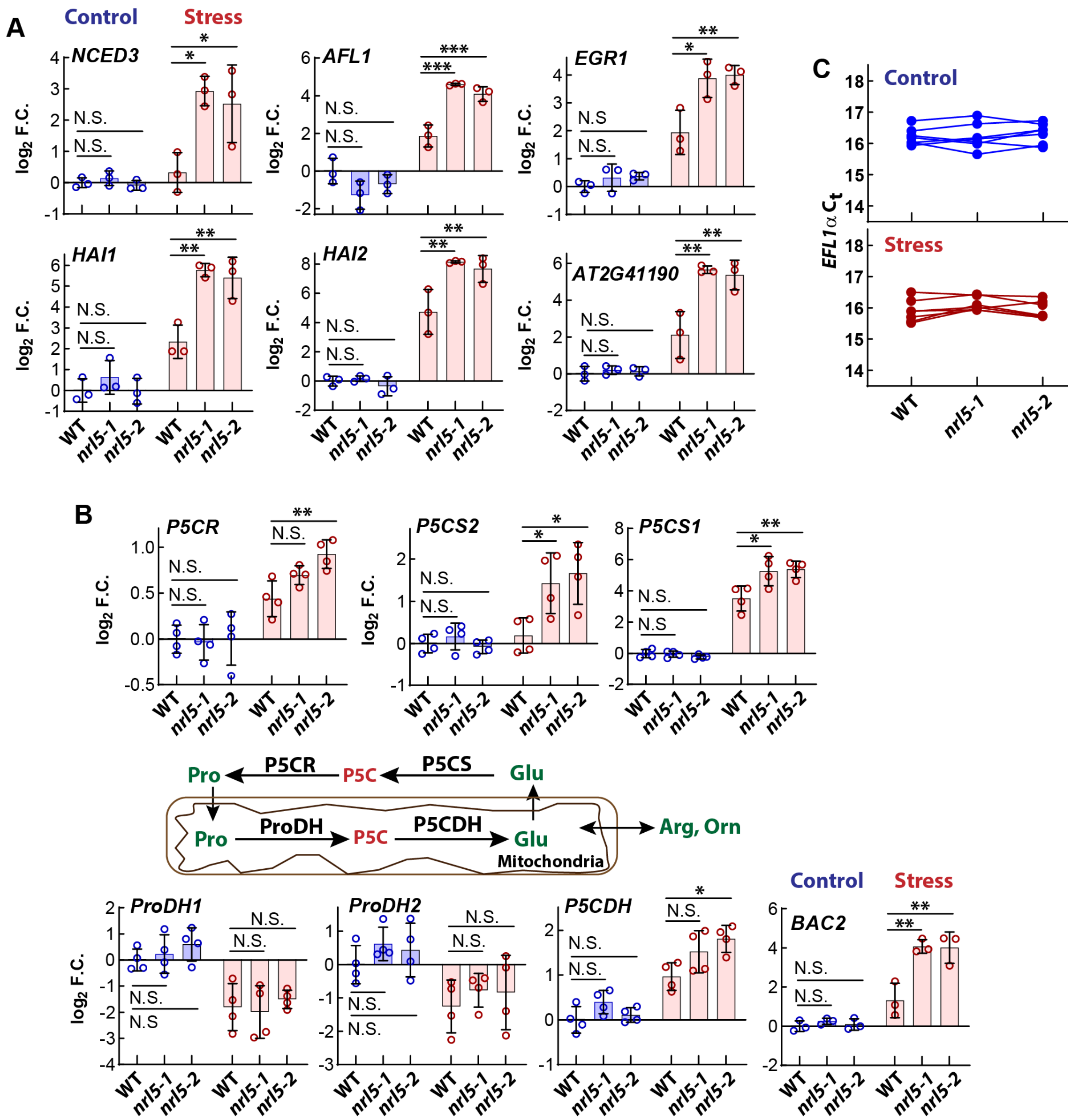


**Fig. S3: *nrl5* mutants have altered expression of genes related to abiotic stress and proline metabolism.**

(**A**) Expression of stress-related genes in wild type (WT), *nrl5-1* and *nrl5-2* in unstressed control and after transfer to low $\psi_w$ stress treatment (-0.7 MPa, 4 days). Gene expression levels are quantified as $\log_2$ fold change (F.C.) versus the unstressed wild type (WT). Data are from quantitative-PCR analysis (mean ± S.D, n =3) with each sample collected from an independent biological experiment. Three technical replicates were performed per sample. P values are from ANOVA comparison to WT within each treatment. (**B**) Expression of the core proline metabolism genes (and *BAC2* mitochondrial ornithine transporter) in wild type and *nrl5* mutants. Experimental procedures and data analysis are as described in panel A except that samples were collected from four independent experiments (three experiments for *BAC2*). For statistical analyses *, **, *** and N.S. represent P ≤ 0.05, P ≤ 0.01, P ≤ 0.001 and non-significant differences, respectively. (**C**) $C_t$ values of the *EFL1α* reference gene for the experiments shown in A and B. Data are from three experiments shown in A and four experiments in B. ANOVA comparison of wild type to both mutants found no significant differences, indicating that expression of the reference gene was stable in both treatments.

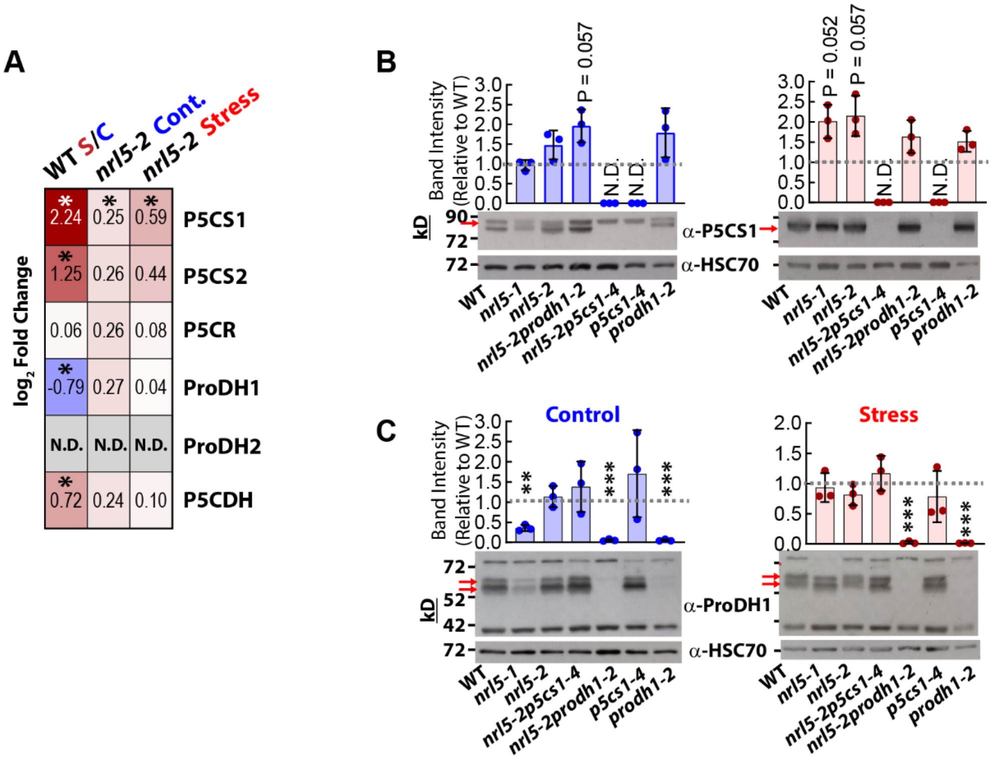


**Fig. S4: *nrl5* mutants have increased P5CS protein level.**

(**A**) Protein levels of the core proline cycle enzymes determined by proteomics analysis. Note that the proteomics analysis is more fully presented in Fig. S18 to S20 and discussed in later sections of the main text. Asterisks (*) indicate a significant change in abundance (log2 fold change ≥ 0.5, P ≤ 0.05) in Wild Type (WT) stress versus control (S/C) or *nrl5-2* compared to wild type in the control and stress (-0.7 MPa) treatments. N.D. = Not Detected. (**B**) Immunoblotting also detected increased abundance of P5CS1 in *nrl5* mutants exposed to low $\psi_w$ stress. Band intensities were quantified and expressed relative to wild type on the same blot. Data are means ± S.D from three independent experiments. P-values are from one sample T-test. Error bars show S.D. N.D. = Not Detected. For the representative blots shown, red arrow indicates the P5CS1 specific band (which is absent in *p5cs1-4*) used for quantitation. Blots were re-probed with HSC70 specific antisera as a loading control. (**C**) Immunoblotting of ProDH1 in *nrl5* mutants. Data presentation and analysis are as described in B. For the representative blots shown, red arrows indicate the ProDH1 specific bands (which are absent in *prodh1-2*) used for quantitation. Data are means ± S.D from three independent experiments. P-values are from one sample T-test. ** and *** indicate P ≤ 0.01 and P ≤ 0.001, respectively. Error bars show S.D.

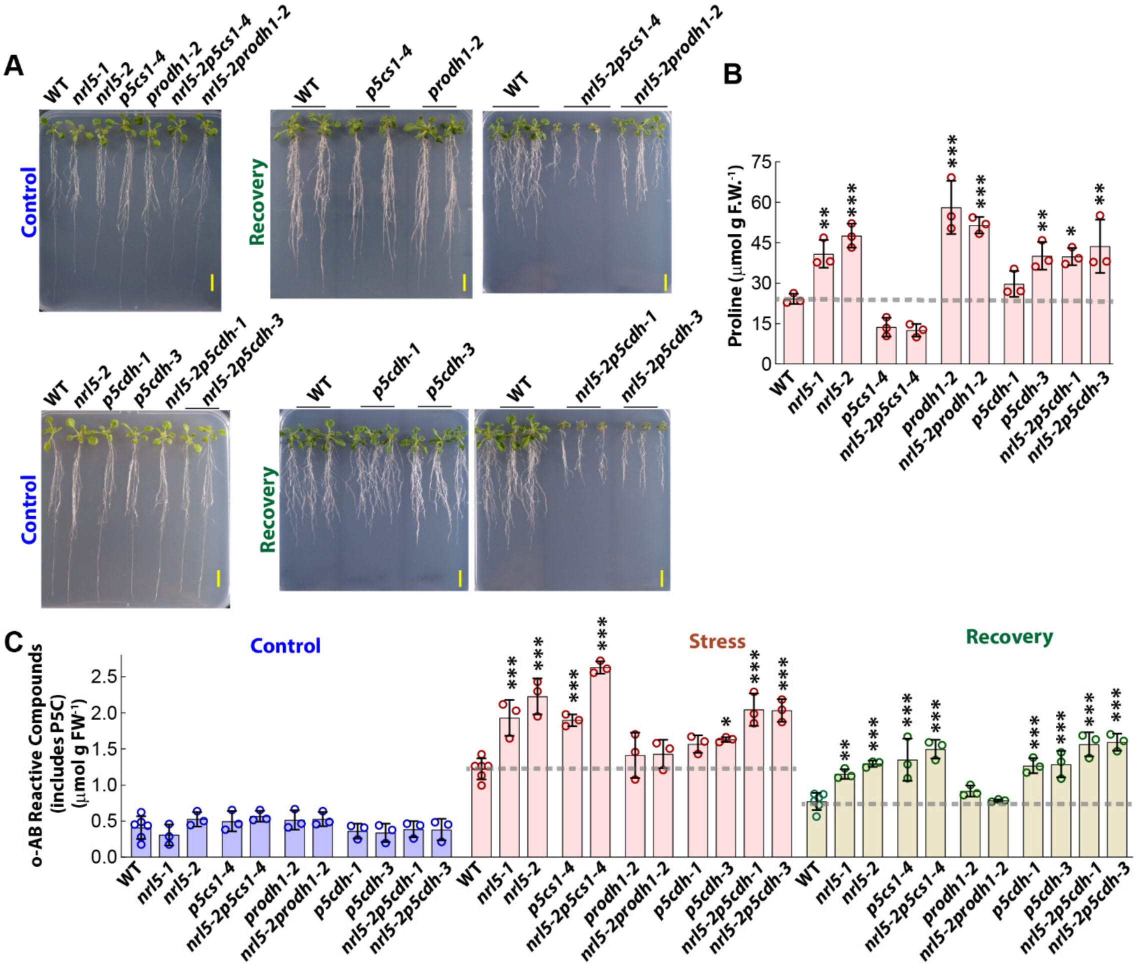


**Fig. S5: Low $\psi_w$ sensitivity of *nrl5* mutants is associated with increased abundance of o-AB reactive compounds (P5C) but does not correlate with the level of proline accumulation.**

(**A**) Representative images of *nrl5* and proline metabolism mutants in control and recovery treatments. Low $\psi_w$ stress images of these mutants and dry weight quantitation for all treatments are shown in Fig. 1E and F. Scale bars indicate 1 cm. (**B**) Low $\psi_w$-induced (-0.7 MPa, 4 days) proline accumulation in *nrl5* and proline metabolism mutants. Each data point is from an independent experiment with 3 biological replicates assayed per experiment. Data were analyzed by ANOVA with comparison to WT (n = 3). Error bars indicate S.D. Gray dashed line indicates the wild type level. (**C**) The level of o-aminobenzaldehyde (o-AB)-reactive compounds measured four days after transfer of seedlings to control or -0.7 MPa low $\psi_w$ stress and 1 day after transfer of seedlings from low $\psi_w$ back to unstressed control (recovery). Note that the background levels of o-AB reactive compounds seen in wild type may include unidentified metabolites that react with o-AB. However, the increased level of o-AB reactive compounds seen in *nrl5* mutants in the stress and recovery treatments, which is not present in *nrl5-2prodh1-2* but is still present in *nrl5-2p5cs1-4* and *nrl5-2p5cdh*, represents changes in the level of $\Delta^1$-pyrroline-5-carboxylate (P5C). P-values are from ANOVA comparison to WT in the same treatment (n = 6 for wild type and n = 3 for mutants with each sample from an independent experiment, two biological replicates assayed for each experiment). Error bars show S.D. Gray dashed lines indicated the wild type P5C level in stress and recovery treatments.

For statistical analyses *, ** and *** indicate $P \leq 0.05$, $P \leq 0.01$ and $P \leq 0.001$, respectively

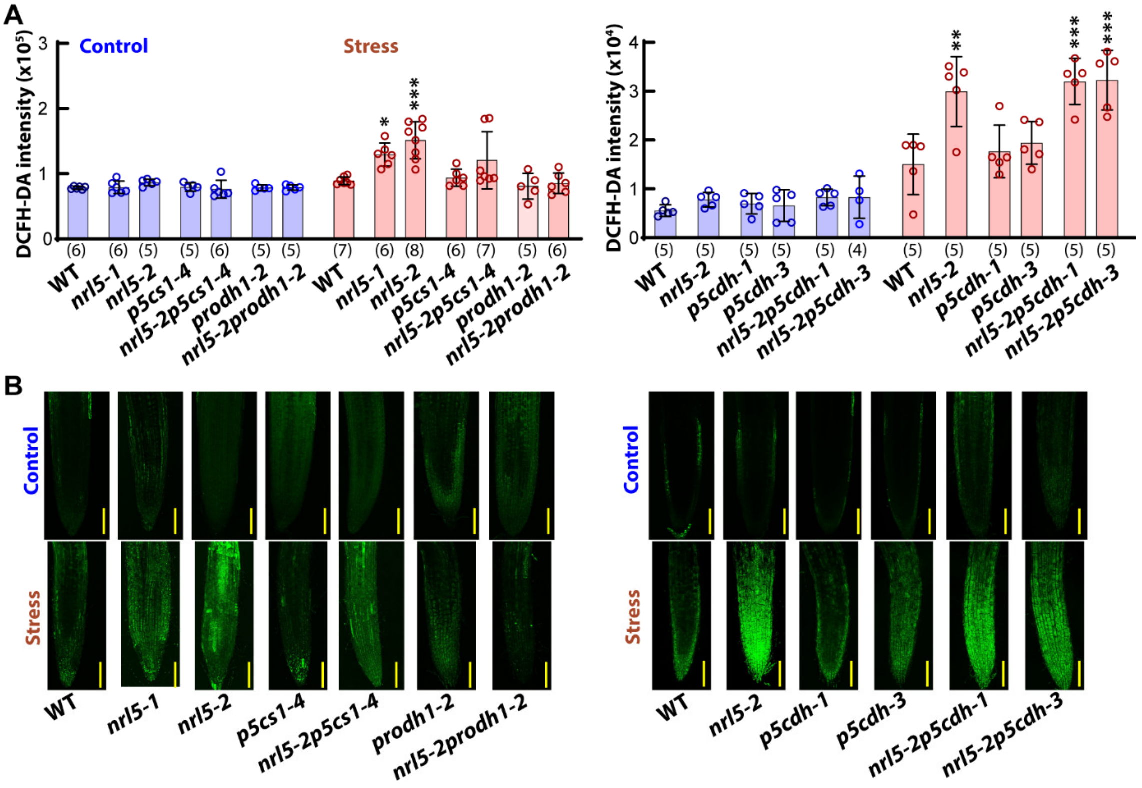


**Fig. S6: Low $\psi_w$ sensitivity of *nrl5* is associated with increased ROS.**
(**A**) ROS staining found that *nrl5* mutants had increased ROS accumulation and this was alleviated by blocking proline catabolism upstream of P5C in *nrl5-2prodh1-2* but not by blocking proline catabolism downstream of P5C in *nrl5-2p5cdh*. Data are means ± S.D. (n values indicated in parentheses, each data point is from one root tip). Data are from one experiment which was repeated with consistent results. Data were analyzed by ANOVA with comparison to wild type in the same treatment. *, ** and *** indicate $P \leq 0.05$, $P \leq 0.01$ and $P \leq 0.001$, respectively. (**B**) Representative fluorescence images of root tip DCFH-DA staining. Scale bars indicate 100 μm.

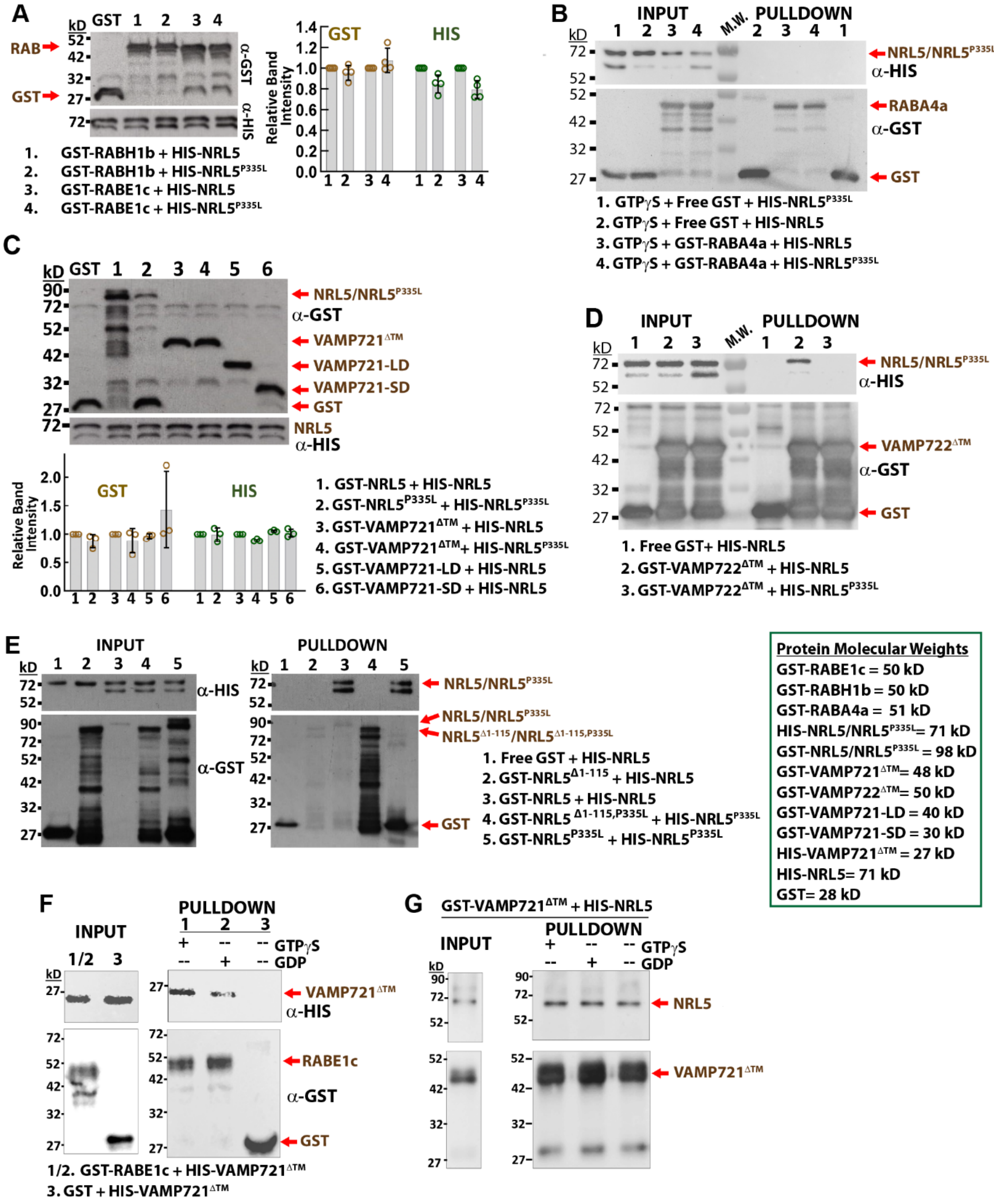


**Fig. S7: NRL5, but not NRL5$^{P335L}$, interacts with some clades of RAB small GTPases as well as VAMP721/722; NRL5$^{P335L}$ does not block NRL5 self-interaction.**

(**A**) Representative blot showing input for the NRL5-RABH1b/RABE1c pull-down experiments shown in Fig. 2A. Graph shows the quantitation of band intensities for all four replicate experiments with the protein amount for the RAB-NRL5$^{P335L}$ assays expressed relative to the RAB-NRL5 input protein level. (**B**) NRL5 did not interact with RABA4a in the presence of GTPγS. The GST pull-down reactions were assembled in the same manner as for the NRL5 interaction with RABH1b and RABE1c. The experiment was repeated with consistent results. (**C**) Representative blot showing input protein amounts for the NRL5-VAMP721 pull-down experiments shown in Fig. 2B. Graph shows the quantitation of band intensities for all three replicate experiments. (**D**) VAMP722 interacted with NRL5 but not NRL5$^{P335L}$. The GST pull-down assay was assembled in the same manner as the NRL5-VAMP721 pull down assays. The experiment was repeated with consistent results. (**E**) Self-interaction of NRL5 required the BTB domain (amino acids 1-115) but was minimally affected by the P335L substitution. The experiment was repeated with consistent results. **(F)** RABE1c, in both its activated (GTPγS-bound) or inactive (GDP-bound) form, interacted with VAMP721. Pull-down assays were conducted in the same manner as others shown here. The experiment was repeated with consistent results. **(G)** NRL5-VAMP721 interaction was not affected by the presence of either GTPγS or GDP. The experiment was repeated with consistent results.

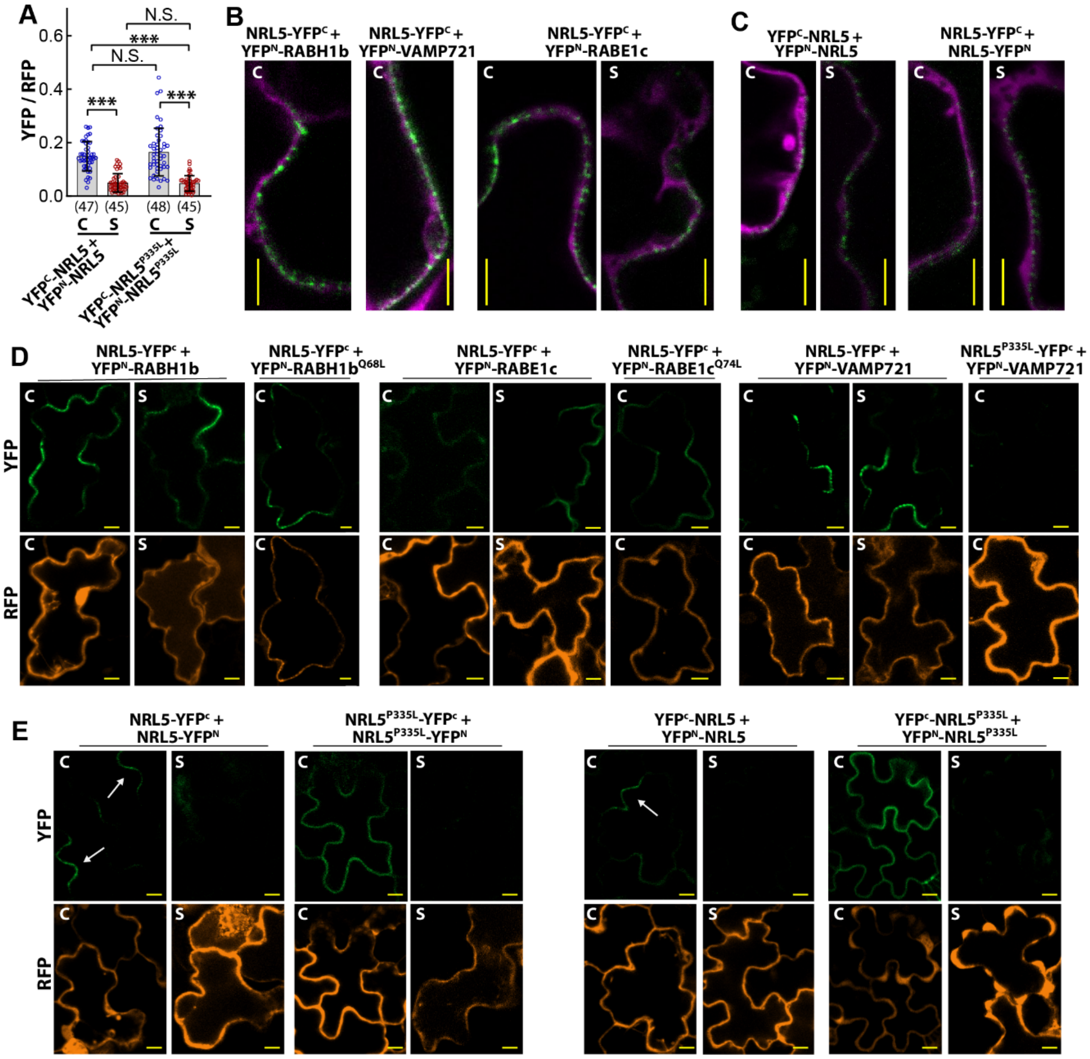


**Fig. S8: NRL5 interacts with RABE1c/RABH1b and VAMP721 in puncta close to the plasma membrane; NRL5$^{P335L}$ homo-complexes lose polarity.**
(**A**) rBiFC self-interaction data for NRL5 and NRL5$^{P335L}$ with N-terminal fusion to split-YFP. The results are consistent with the NRL5 self-interaction with C-terminal split-YFP fusion shown in Fig. 2E. Data points are YFP/RFP ratios of individual cells and are combined from two independent experiments. Data are means ± S.D. analyzed by Kruskal-Wallis test (n values shown in parentheses). *** indicates P ≤ 0.001, N.S. indicates no significant difference. (**B**) Additional high-resolution images of rBiFC signal (green) for NRL5-RAB and NRL5-VAMP721 interactions or treatments not shown in Fig. 2F. For all proteins tested, the rBiFC signal was typically seen as a foci along the periphery of the cell and was polarized along one edge of the cell (see panel D). Propidium iodide staining of cell wall is shown in magenta. Scale bars indicate 10 µm. C = Control, S = Stress (-0.7 MPa). (**C**) Additional high-resolution rBiFC images of NRL5 self-interaction. (**D**) Representative whole-cell images used for quantitation of YFP/RFP for NRL5 interaction with RABH1b, RABE1c or VAMP721. Note that the rBiFC signal (green) was not evenly distributed around the cells but rather was concentrated in one or two lobes of the leaf pavement cells. In contrast, the RFP reporter used for normalization (orange) was distributed evenly around the cytoplasm. Scale bars indicate 10 µm. C = Control, S = Stress (-0.7 MPa). (**E**) Representative whole cell images for NRL5 and NRL5$^{P335L}$ self-interaction using either N- or C-terminal fusion to split YFP. For NRL5 self-interaction, white arrows indicate polarized patches of signal that were detected in only one or two lobes of the leaf pavement cells. In contrast, NRL5$^{P335L}$ self-interaction was detected all around the cell. Scale bars indicate 10 µm. C = Control, S = Stress (-0.7 MPa).

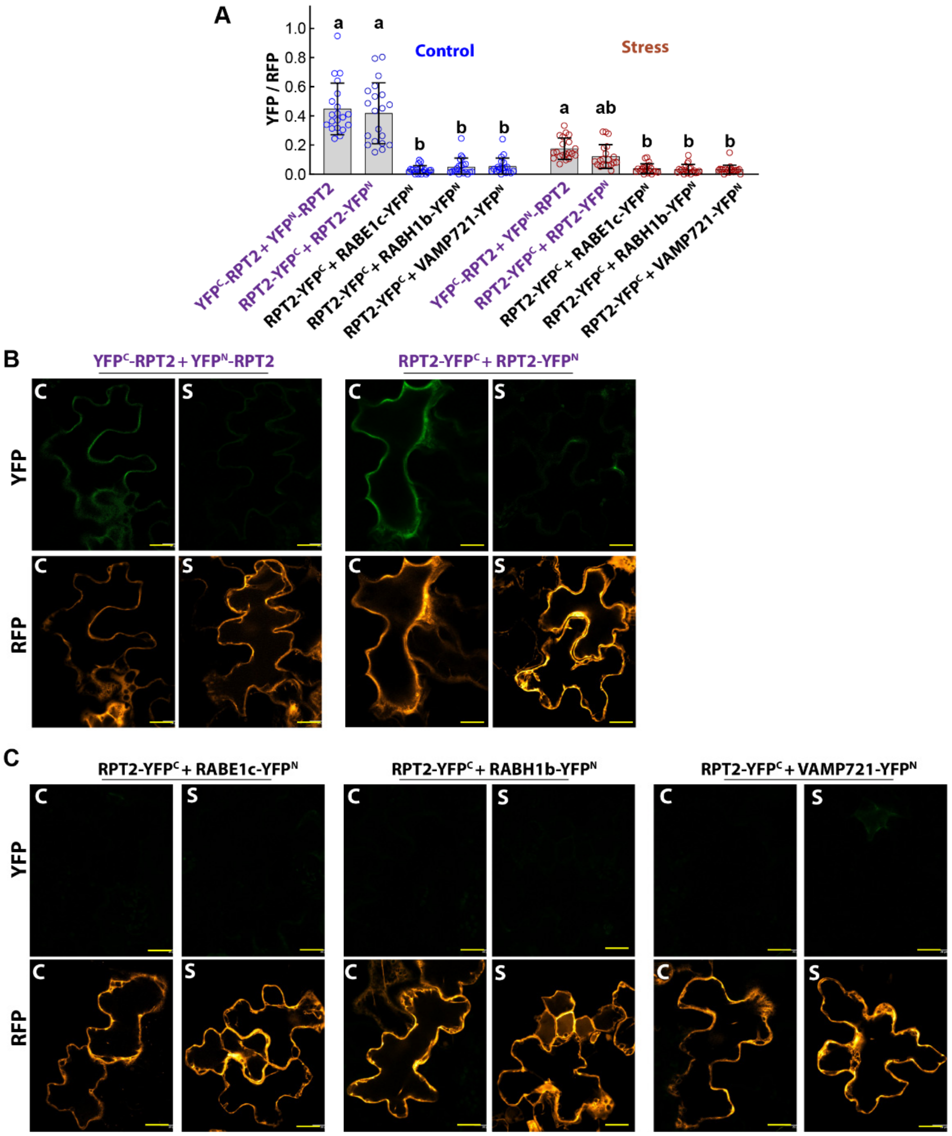


**Fig. S9: The NPH3-domain protein RPT2 interacts with itself but had minimal or no interaction with RABE1c, RABH1b or VAMP721 and did not exhibit polar localization.** **(A)** Quantitation of rBiFC fluorescence intensity ratio showed that RPT2 interacted with itself with similar or higher BiFC fluorescence intensity compared to NRL5 self-interaction. Like NRL5, RPT2 self-interaction rBiFC fluorescence intensity was markedly decreased by low $\psi_w$ (-0.7 MPa). In contrast to NRL5, rBiFC of RPT2 with RABE1c, RABH1b or VAMP721 but had very low fluorescence, indicating minimal or no protein interaction. Data points are YFP/RFP ratios of individual cells and are combined from two independent experiments (n = 20). Error bars indicate S.D. Data were analyzed by Kruskal-Wallis test. Groups sharing the same letter are not significantly different from one another ($P \leq 0.05$). **(B)** Representative images of RPT2 self-interaction show that the signal was distributed around the periphery of the cell. This contrasted with the polar localization of NRL5 self-interaction and indicated that the polarity of NRL5 homocomplexes seen in rBiFC assays was a specific feature of NRL5 and not an artifact of the rBiFC experimental system. C = Control, S = Stress (-0.7 MPa). Scale bars indicate 20 µm. **(C)** Representative images showing the lack of RPT2 fluorescence signal for the RABE1c, RABH1b and VAMP721 interaction assays. Scale bars indicate 20 µm.

**A**

| Protein | $k_{cat}$ (min$^{-1}$) | $K_m$ (mM) | $V_{max}$ (μmol min$^{-1}$) | $A_{max}$ (μmol min$^{-1}$ μmol protein$^{-1}$) x10$^6$ |
|---|---|---|---|---|
| **NRL5** (in vitro translation) | 2.351 (1.42 to 7.44) | 27.74 (10.58 to 143.2) | 2.32 | 0.116 |
| **NRL5** (*E. coli* produced) | 2.91 (2.25 to 4.04) | 11.38 (6.08 to 22.8) | 2.91 | 0.145 |
| **NRL5$^{P335L}$** (in vitro translation) | 0.135 (0.098 to 0.205) | 6.93 (2.69 to 19.1) | 0.14 | 0.007 |
| **NRL5$^{P335L}$** (*E. coli* produced) | 0.81 (0.52 to 1.49) | 6.927 (1.97 to 28.01) | 0.81 | 0.041 |
| **NRL6** | 3.067 (2.284 to 4.788) | 34.3 (20.69 to 66.66) | 3.07 | 0.154 |
| **NRL15** | 3.166 (2.304 to 5.257) | 38.38 (22.57 to 79.51) | 3.17 | 0.159 |
| **RPT2** | 2.87 (2.184 to 4.275) | 34.66 (21.75 to 62.99) | 2.87 | 0.144 |
| **NPH3** | 3.361 (2.83 to 4.12) | 14.32 (9.80 to 21.66) | 3.36 | 0.168 |
| **RABH1b** | 2.40 (1.94 to 3.11) | 12.94 (7.89 to 22.25) | 2.40 | 0.120 |
| **RABE1c** | 1.66 (1.37 to 2.05) | 4.761 (2.54 to 8.81) | 1.66 | 0.083 |
| **RABA4a** | 2.20 (1.85 to 2.67) | 6.24 (3.76 to 10.39) | 2.20 | 0.110 |
| **GST** (in vitro translation) | 0.032 (0.0242 to 0.0441 | 4.37 (1.72 to 10.97) | 0.032 | 0.002 |
| **GST** (*E. coli* produced) | 0.054 (0.0304 to 0.254) | 10.03 (1.934 to 132.1) | 0.054 | 0.003 |

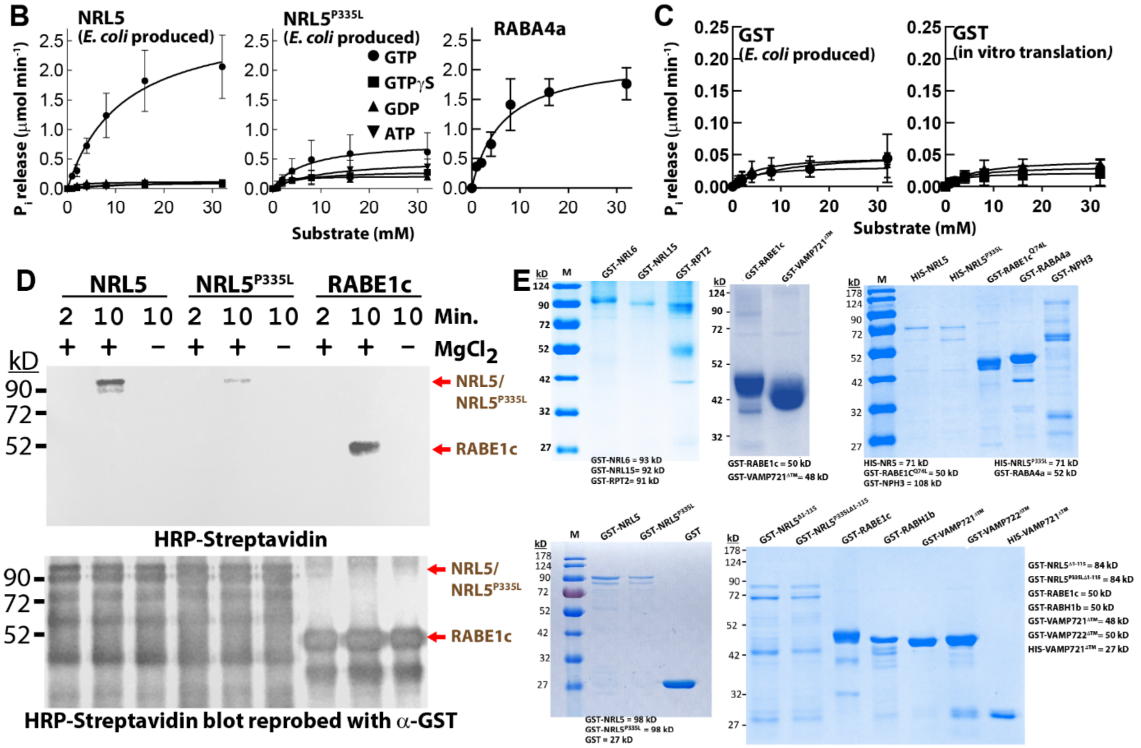


**Fig. S10: NPH3-domain GTPase activity and desthiobiotin-GTP labeling. (Legend on next page)**

**Fig. S10: NPH3-domain GTPase activity and desthiobiotin-GTP labeling.**
(**A**) Kinetic properties of GTPase activities for NRL5, RABs, other NPH3 domain proteins and GST (used as negative control protein as the other proteins assayed included a GST tag for purification). Data were analyzed by nonlinear regression fit to the equation $Y = E_t*k_{cat}*X/(K_m + X)$ using the $K_{cat}$ analysis option in GraphPad Prism 9 (Y = enzyme velocity, X = substrate concentration). The number of catalytic sites per protein was assumed to be 1 ($E_t = 1$). Numbers in parentheses on the table show the 95% confidence intervals of the $k_{cat}$ and $K_m$ values. $A_{max}$ was calculated based on each reaction having a volume of 20 μl and the molar protein concentration indicated in the relevant figure legends (typically 1 μM).
(**B**) GTPase activity of NRL5 and NRL5$^{P335L}$ produced in *E. coli* as well as RABA4a. One μM protein was used in each assay with the indicated substrate concentrations and $P_i$ release quantified after 30 min (reaction volume was 20 μl). Data were analyzed by non-linear regression ($K_{cat}$, $K_m$, and $V_{max}$ values are shown in Fig. S10A). Data are mean ± S.D. (n = 3).
**(C)** GTPase assay data for GST control protein produced in *E. coli* or by in vitro translation. Note that data for GST produced by in vitro translation with GTP substrate is also shown in Fig. 3B for comparison to NRL5$^{P335L}$. Data format and analysis are as described for B.
(**D**) Desthiobiotin-GTP labeling conducted using NRL5 and NRL$^{P335L}$ purified from *E. coli*. Labeling was conducted for 2 or 10 min with or without addition of $MgCl_2$ as indicated in the figure. Note that in this case loading was accessed by stripping and re-probing of the blot with anti-GST after detection with HRP-Streptavidin. Thus, the relatively high protein loading used for the biotin labeling led to high level of background in the loading control blot. The expected molecular weight of GST-NRL5 and GST-RABE1c used here are 98 kD and 50 kD, respectively.
**(E)** Coomassie stained gels showing the purity of *E. coli*-produced proteins used for GTPase assays.

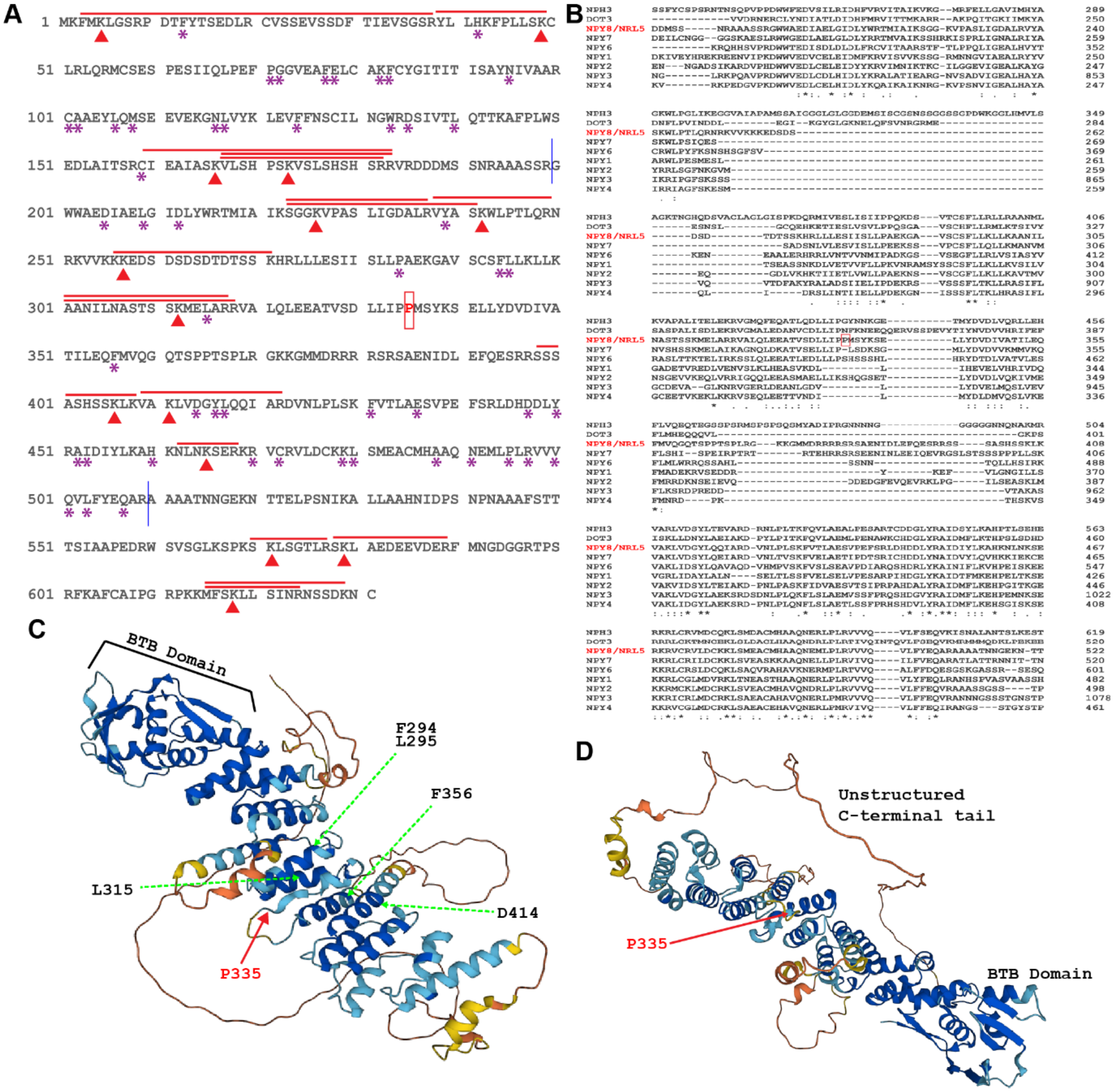


**Fig. S11: Sequence conservation, desthiobiotin-GTP labeling and NRL5 structural model suggest a basis for reduced GTPase activity and protein interactions of NRL5$^{P335L}$.**
(**A**) NRL5 amino acid sequence with positions of biotinylated peptides (red bars) and lysine residues (red arrows) indicated. Asterisks indicate amino acids conserved among the NPY clade (based on alignment in B). (**B**) Amino acid sequence alignment of the NPY-clade of NPH3 domain proteins. Full length protein sequences were used for alignment but for clarity of presentation only the portion of the alignment containing the NPH3 domain is shown. Red box indicates NRL5 P335. (**C**) NRL5 structure predicted by Alphafold. The position of P335 is indicated by red arrow. Selected amino acids that are structurally close to P335, fully conserved among the NPY group of NRL proteins and close to sites biotinylated by desthiobiotin-GTP are marked by green arrows. Note that P335 is part of a linker between highly conserved helices. The NRL5$^{P335L}$ mutation may alter the position of the conserved helices relative to each other and thereby strongly reduce, but not eliminate, GTPase activity and disrupt interaction with RABH1b/RABE1c and VAMP721/722. (**D**) Rotated view to emphasize the planar arrangement of the helices in the NPH3 domain and unstructured C-terminal tail. The conserved helices are connected by unstructured linkers suggesting that ligand binding could alter the relative positions of the helices.

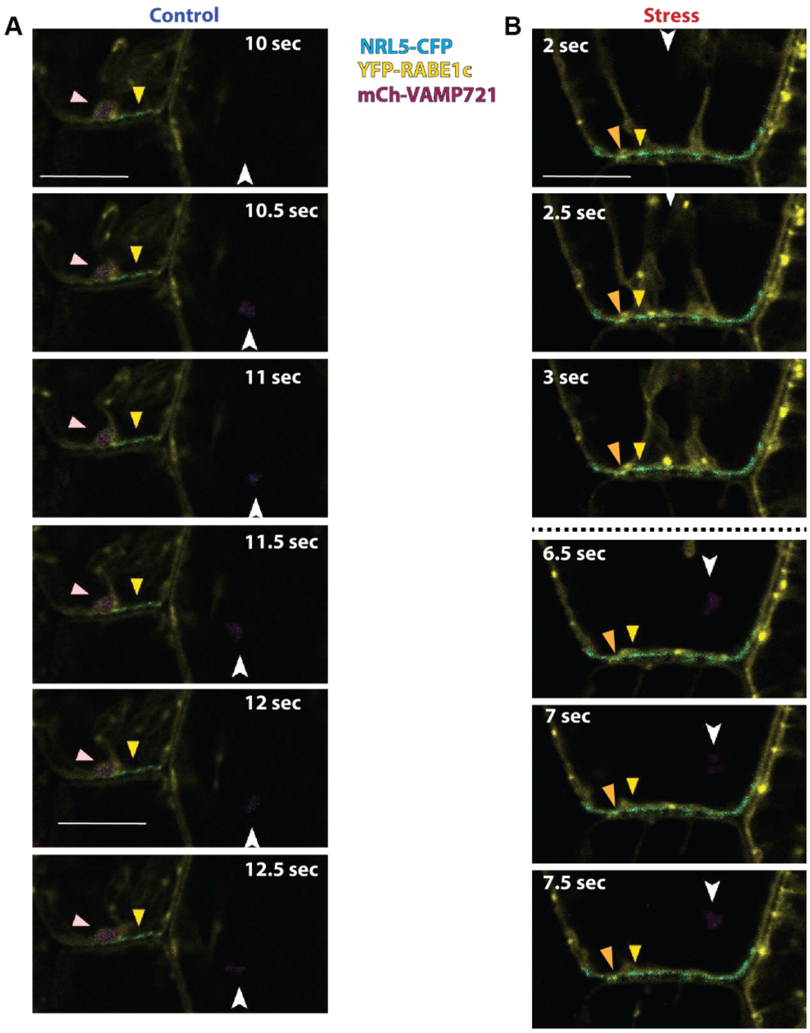


**Fig. S12: Selected frames from Supplementary Movies S4 and S8 showing colocalization patterns of NRL5, RABE1c and VAMP721 in hypocotyl cells.**

(**A**) Selected frames from time lapse microscopy of hypocotyl cells from unstressed plants expressing *NRL5$_{pro}$:NRL5-CFP*, *35S:YFP-RABE1c* and *35S:mCherry-VAMP721*. Pink arrow indicates a plasma membrane-adjacent structure where all three proteins can be observed. Yellow arrow indicates an example of an immobile foci of NRL5 along the plasma membrane. White arrow indicates an internal, motile foci of VAMP721 along with lower level of transient NRL5. Scale bar indicates 10 µm. Supplementary Movies S1 to S3 show the complete time lapse of the NRL5 and RABE1c, NRL5 and VAMP721 and RABE1c and VAMP721 channels, respectively. Supplementary Movie S4 shows the complete time lapse of all three channels, including the frames shown here.

(**B**) Selected frames from time lapse microscopy of hypocotyl cells of plants exposed to low $\psi_w$ (-0.7 MPa) for 4 days. The seedlings used were from the same transgenic line described in A. Orange arrow shows example of a transient foci of NRL5 and RABE1c colocalization. Yellow arrow shows another example of an immotile foci of NRL5 along the plasma membrane. Note that several other foci of NRL5 along the membrane also have transient RABE1c colocalization and proximity to RABE1c-containing motile vesicle-like particles during the complete time course. White arrow shows internal foci of VAMP721 with transient lower level of NRL5. Scale bar indicates 10 µm. Supplementary Movies S5 to S7 show the complete time lapse of the NRL5 and RABE1c, NRL5 and VAMP721 and RABE1c and VAMP721 channels, respectively for the stress treatment. Supplementary Movie S8 shows the complete time lapse of all three channels, including the frames shown here.

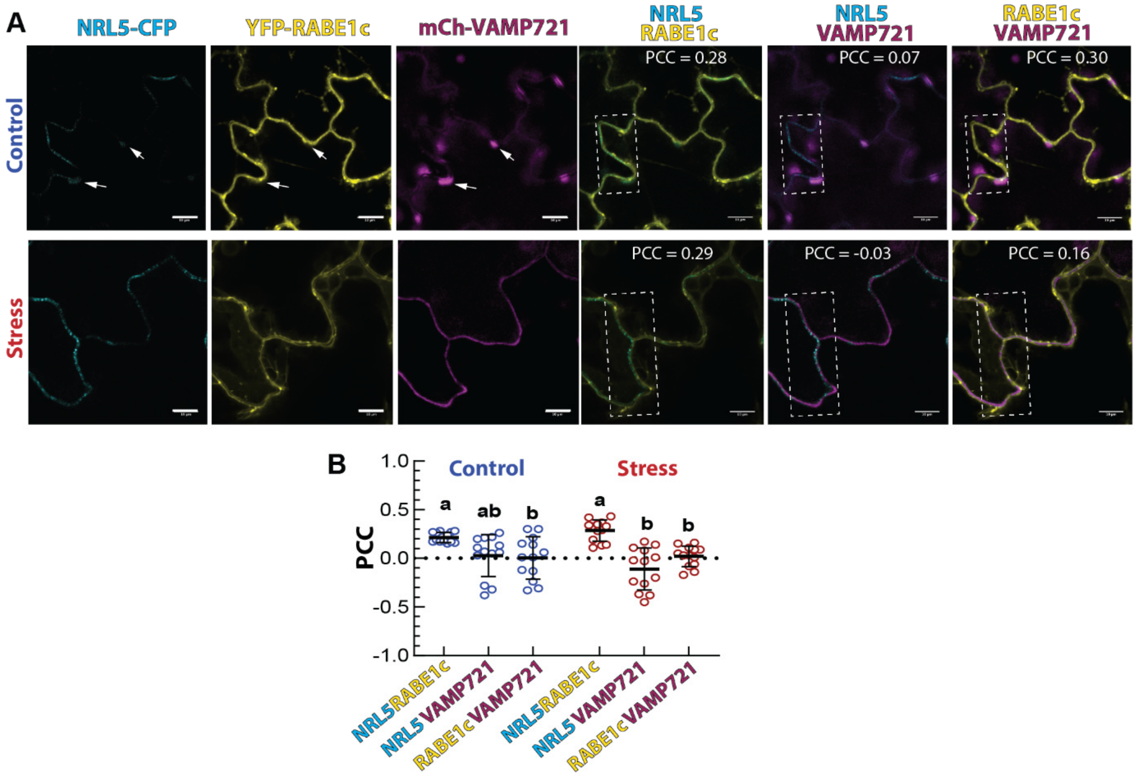


**Fig. S13: NRL5, RABE1c and VAMP721 colocalization in leaf pavement cells.**
(**A**) Representative images of leaf pavement cells from a transgenic line expressing *NRL5*$_{pro}$*:NRL5-CFP*, *35S:YFP-RABE1c* and *35S:mCherry-VAMP721*. Single channel images as well as merged images of two channels are shown. Five-day-old seedlings were transferred to fresh control agar plates (Control) or the -0.7 MPa PEG-infused agar plates (Stress) for four days before imaging. White arrows indicate structures that contain VAMP721 and a lower level of NRL5 adjacent to foci of RABE1c. Dashed line boxes indicate the area selected for Pearson Correlation Coefficient (PCC) analysis to quantify co-localization between the indicated two proteins. PCC values for each image are shown. Scale bars indicate 10 µm.
**(B)** PCC analysis of the indicated protein pairs in control and stress treatment. Cells were analyzed from leaves of multiple plants for each genotype and treatment (n =13). Black bars indicated the mean and error bars show S.D. Data not sharing the same letter are significantly different from one another (ANOVA, $P \leq 0.05$).

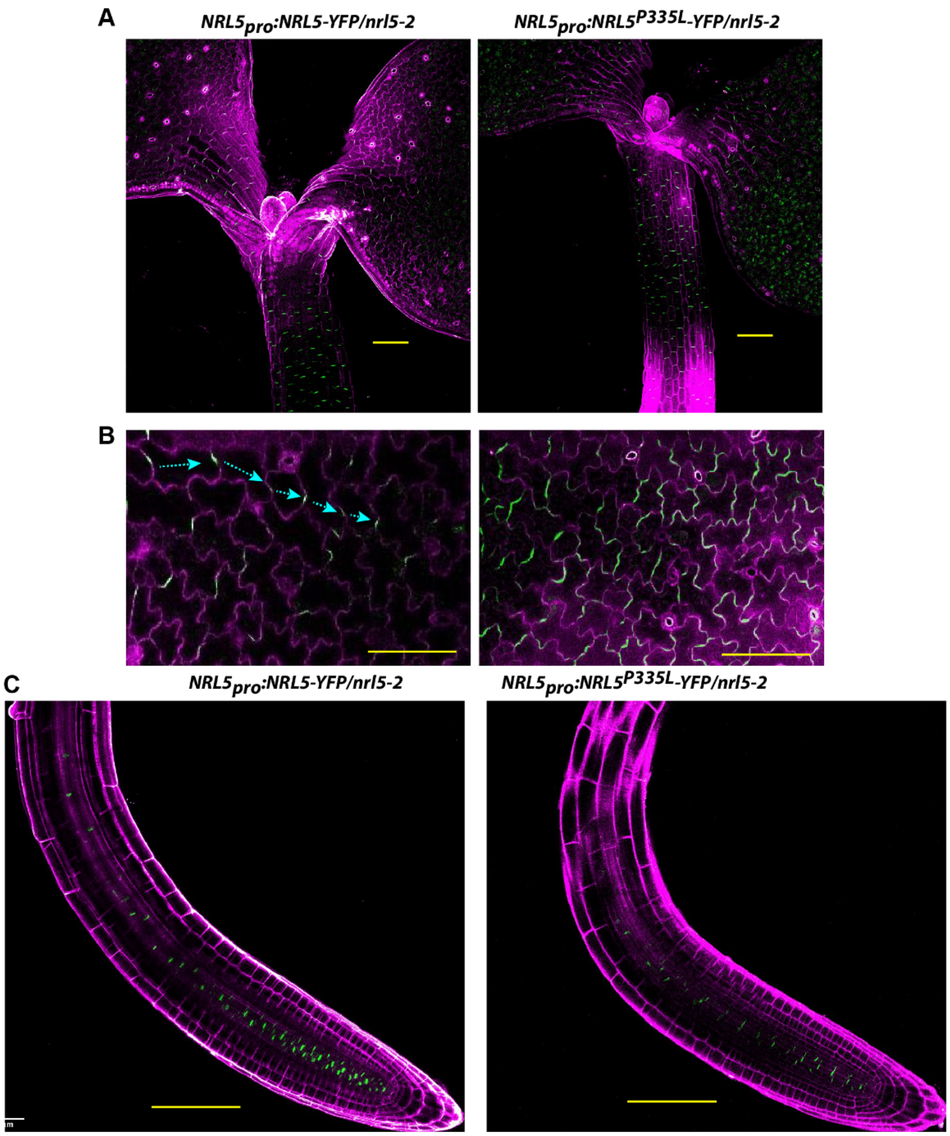


**Fig. S14: NRL5 is polarly localized in leaf, hypocotyl and root cells.**
(**A**) Maximum intensity projection images of plant expressing *NRL5*$_{pro}$*:NRL5-YFP* or *NRL5*$_{pro}$*:NRL5*$^{P335L}$*-YFP* showing NRL5 protein (green) accumulation pattern in leaf and hypocotyl of three-day-old seedlings. Magenta shows propidium iodide staining of cell wall. NRL5 was detected in leaf and hypocotyl epidermal cells. Note the polar localization of NRL5 on the rootward (bottom) side of cells in the hypocotyl, petiole and into the developing leaf. Scale bars indicate 100 µm. Essentially identical patterns of NRL5 protein accumulation were seen in three independent transgenic lines for each construct.
**(B)** Enlarged images of more mature part of the leaf containing lobed pavement cells. Turquois arrows indicate the typically observed pattern where patches of NRL5 were located on opposite sides of leaf pavement cells and often appeared to trace a path through the leaf epidermis. These patches did not occur next to stomata or small, non-lobed cells. Note that for NRL5$^{P335L}$, the polarity was diminished and the protein found in multiple, and wider, patches along the cell periphery. See Fig. 6 and Fig. S15 for further analysis of NRL5 polarity in leaf pavement cells. Scale bars indicate 100 µm.
**(C)** Root tip images of three-day-old seedlings. NRL5 was detected along the bottom edge of cells in the stele tissue. In root, NRL5$^{P335L}$ had lower YFP fluorescence than NRL5 across several transgenic lines. See Fig. S16 for further imaging of NRL5 and NRL5$^{P335L}$ in root tip. Scale bars indicate 100 µm.

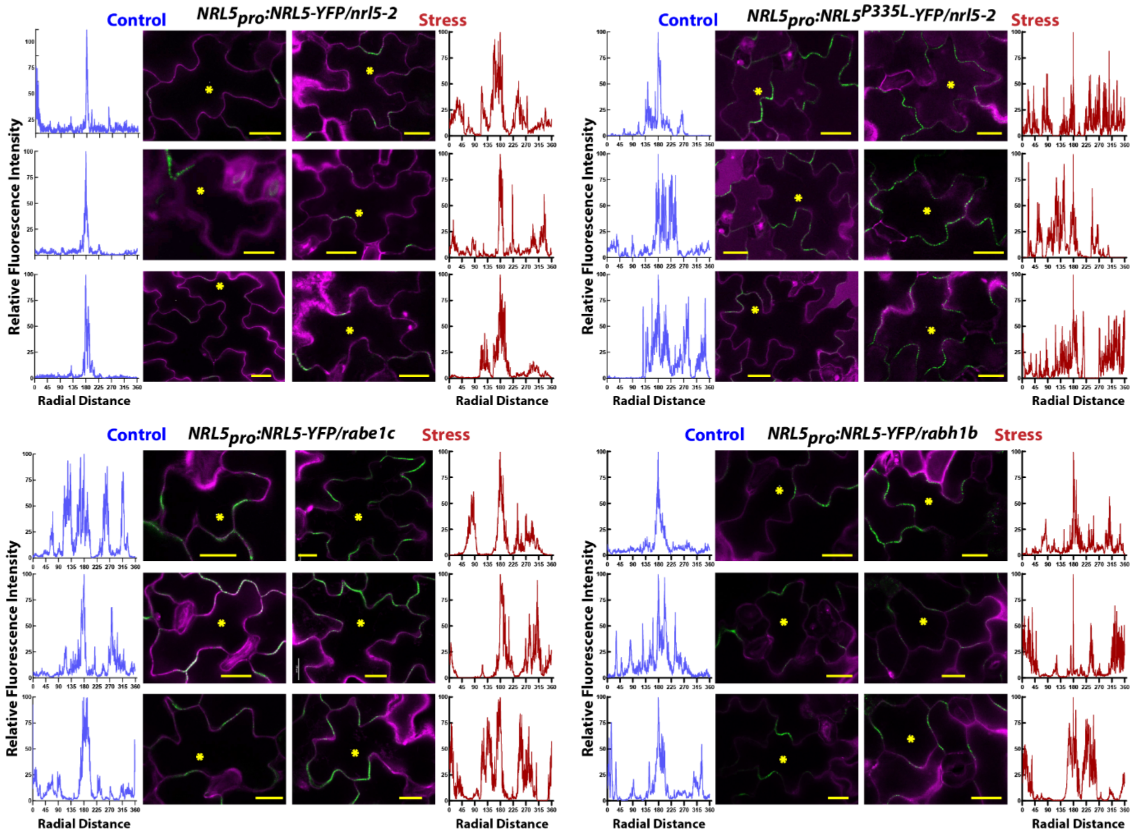


**Fig. S15: Additional images and line scan analysis of NRL5 and NRL5$^{P335L}$ in leaf pavement cells.**

Additional representative images from the experiments described in Fig. 6A. The images and line scan analysis around the periphery of the cell demonstrate that that low $\psi_w$, the P335L mutation and lack of RABE1c or RABH1b diminish NRL5 polarity. Asterisk marks the cell that was selected for the line scan analysis shown in the graph next to the image. Graphs on the left and right side of the images show relative fluorescence intensity along a line scan tracing of the cell periphery. For each cell analyzed, the point of highest fluorescence intensity was set to 100 and other points normalized to that value. The normalized data were plotted on the basis of radial distances (360°) around the cell with the point of highest fluorescence set to 180°. Note that the correspondence of radial distance to physical distance will vary for cells of different sizes. Cells were imaged four days after transfer of five-day-old seedlings to either fresh control media or low $\psi_w$ (-0.7 MPa). Scale bars indicate 20 μm.

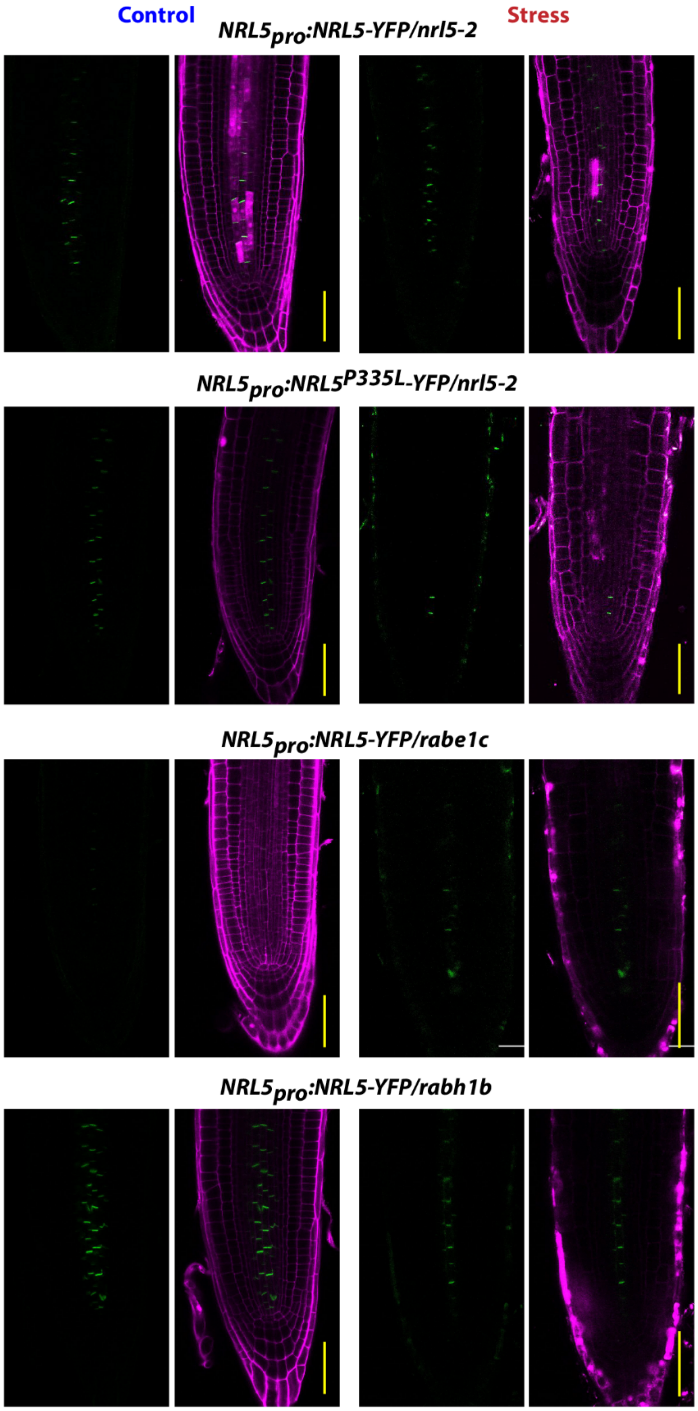


**Fig. S16: NRL5 or NRL5$^{P335L}$ localization in root cells.**
NRL5 localized along the bottom edge of cells in the root stele. In contrast to leaf and hypocotyl cells, changes in polarity were not as apparent in the root. In root, the main effect of low $\psi_w$, the P335L mutation or loss of RABE1c was to decrease NRL5 protein level, especially for cells more distal from the quiescent center. Loss of RABH1b had lesser effect on NRL5 protein level or polarity in root cells. Scale bars indicate 50 µm.

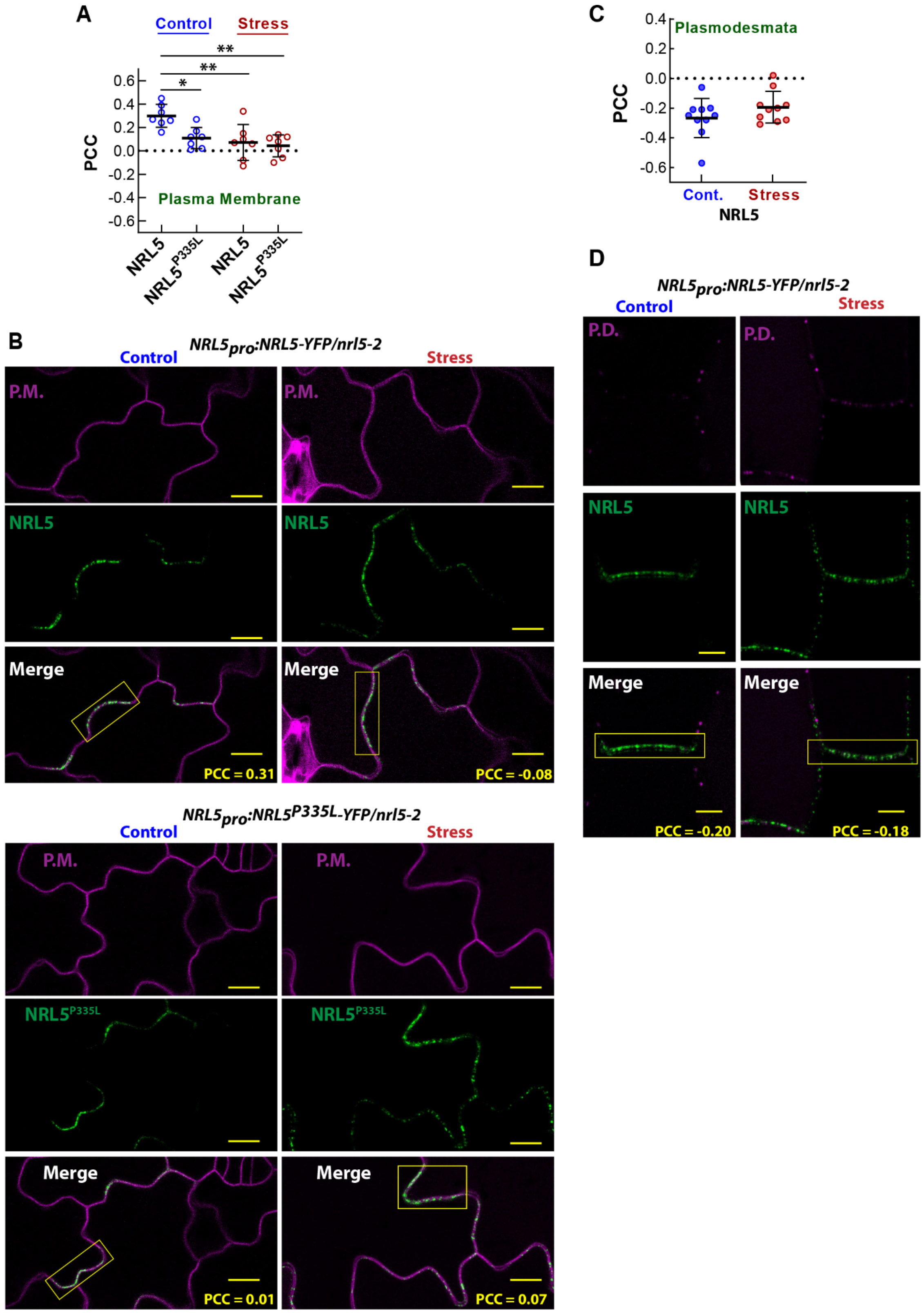


**Fig. S17: The P335L mutation and low $\psi_w$ diminish NRL5 co-localization with plasma membrane; NRL5 does not co-localize with plasmodesmata. (Legend on next page)**

**Fig. S17: The P335L mutation and low $\psi_w$ diminish NRL5 co-localization with plasma membrane; NRL5 does not co-localize with plasmodesmata.**

(**A**) NRL5, but not NRL5$^{P335L}$ partially co-localized with a plasma membrane (P.M.) marker. Low $\psi_w$ stress (-0.7 MPa, 4 days) significantly decreased NRL5 plasma membrane co-localization. NRL5$^{P335L}$ had little plasma membrane localization in either control or stress treatment, as indicated by Pearson Correlation Coefficient (PCC) close to zero. Data were analyzed by ANOVA comparison to NRL5 in the unstressed control (n = 7). * and ** indicate $P \leq 0.05$ and $P \leq 0.01$, respectively. Error bars indicate S.D. Note that similar results of decreased plasma membrane co-localization during low $\psi_w$ for NRL5 and constitutively low plasma membrane co-localization for NRL5$^{P335L}$ were also obtained in hypocotyl cells.

**(B)** Representative images of NRL5/NRL5$^{P335L}$-plasma membrane co-localization in leaf pavement cells. Regions of interest selected for PCC analysis are indicated by yellow boxes. Scale bars indicate 10 µm.

(**C**) Quantitation of NRL5 co-localization with plasmodesmata (P.D.) marker in hypocotyl cells. A similar lack of plasmodesmata-NRL5 co-localization was seen in leaf pavement cells. Error bars show S.D.

**(D)** Representative images showing lack of NRL5 co-localization with plasmodesmata. Scale bars indicate 5 µm.

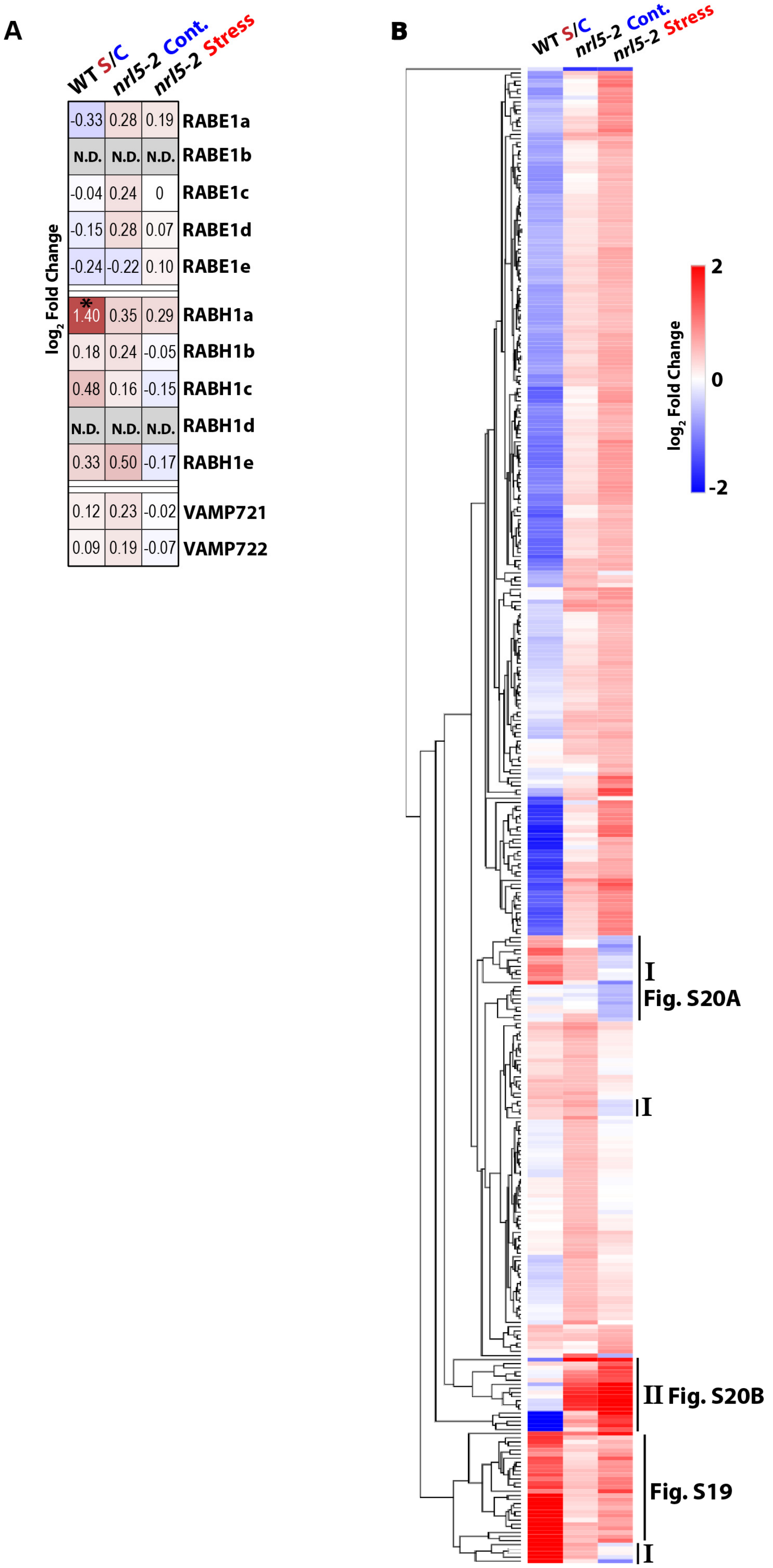


**Fig. S18: Proteomics analysis of *nrl5-2* finds extensive changes in protein abundance but no effect on RABE1, RABH1 or VAMP721/722 (Legend on next page).**

**Fig. S18: Proteomics analysis of *nrl5-2* finds extensive changes in protein abundance but no effect on RABE, RABH or VAMP721/722.**
(**A**) Proteomic data of the effect of *nrl5-2* and low $\psi_w$ on RABE1, RABH1 and VAMP721/722 protein levels. Asterisks (*) indicate a significant change in abundance ($\log_2$ fold change ≥ 0.5, P ≤ 0.05). N.D. = Not detected.
(**B**) Heat map of proteins having altered abundance ($\log_2$ fold change ≥ 0.5, P ≤ 0.05) in *nrl5-2* compared to wild type in either the stress or control treatment, along with their corresponding low $\psi_w$ -induced change in abundance in wild type. A group of proteins that is hyper-induced in NRL5 is marked here and shown in detail in Fig. S19. Similarly, the Roman numerals I and II indicate protein groupings of interest that are shown in detail in Fig S20. For proteomics analysis, low $\psi_w$ stress was imposed by transferring seedlings to -0.7 MPa PEG-agar plates for 4 days. Seedlings in the unstressed control were transferred to fresh agar plates without PEG for the same amount of time. The heat map was constructed with the Morpheus utility using the Euclidian distance method. Full lists of differentially abundant proteins can be found in Data S3-S5.

log$_2$ Fold Change

| WT S/C | *nrl5-2* Cont. | *nrl5-2* Stress | Localization | | | |
|---|---|---|---|---|---|---|
| 1.53 | 1.09 | 1.88 | Plastid | AT1G61800 | GPT2 | glucose6-Phosphate/phosphate transporter 2 |
| 1.62 | 0.53 | 0.38 | Plastid | AT1G17745 | PGDH | 3-Phosphoglycerate dehydrogenase |
| 1.58 | 0.11 | 0.61 | Nuc. | AT1G51140 | AKS1 | basic helix-loop-helix-type transcription factor |
| 1.36 | 0.35 | 0.51 | Vac. | AT2G41190 | | Transmembrane amino acid transporter family |
| 1.02 | 0.42 | 0.71 | Cyt./P.M. | AT1G20450 | ERD10/LTI29 | Dehydrin |
| 0.84 | 0.22 | 0.84 | Plastid | AT2G21590 | APL4 | large subunit of ADP-glucose pyrophosphorylase |
| 1.41 | 0.25 | 1.29 | Nuc. | AT1G80130 | | Tetratricopeptide repeat (TPR)-like protein |
| 1.35 | 0.31 | 0.81 | P.M. | AT1G03940 | | HXXXD-type acyl-transferase |
| 1.25 | 0.37 | 0.88 | Plastid | AT4G39210 | APL3 | large subunit of ADP-glucose pyrophosphorylase |
| 1.28 | 0.50 | 0.85 | P.M. | AT4G24000 | CSLG2 | CELLULOSE SYNTHASE LIKE G2 |
| 1.45 | 0.45 | 0.78 | Mito. | AT4G14090 | | anthocyanidin 5-O-glucosyltransferase |
| 1.10 | 0.43 | 1.02 | Cyt. | AT5G17220 | GSTF12 | phi class glutathione transferase |
| 1.44 | 0.49 | 1.05 | Cyt. | AT4G22880 | LDOX | leucoanthocyanidin dioxygenase |
| 1.59 | 0.55 | 0.99 | P.M. | AT5G54060 | UGT79B1 | Encodes a anthocyanin 3-O-glucoside: 2"-O-xylosyl-transferase |
| 0.99 | 0.36 | 1.49 | Nuc. | AT5G52310 | LTI78/RD29A | LOW-TEMPERATURE-INDUCED 78, RESPONSIVE TO DESICCATION 29A |
| 2.90 | 0.43 | 0.64 | E.R. | AT1G16850 | | transmembrane protein |
| 2.64 | 0.35 | 0.87 | Nuc. | AT5G23000 | LTI65/RD29B | LOW-TEMPERATURE-INDUCED 65, RESPONSIVE TO DESICCATION 29B |
| 2.12 | 0.29 | 0.72 | Nuc. | AT1G69260 | AFP1 | ABI FIVE BINDING PROTEIN |
| 2.24 | 0.25 | 0.59 | Cyt. | AT2G39800 | P5CS1 | $\Delta^1$-pyrroline-5-carboxylate synthetase |
| 2.24 | 0.38 | 0.83 | Plastid | AT2G42530 | COR15B | COLD REGULATED 15B |
| 2.13 | 0.32 | 0.93 | Perox. | AT5G06760 | LEA4-5 | LATE EMBRYOGENESIS ABUNDANT 4-5 |
| 1.96 | 0.11 | 0.72 | E.C. | AT5G15960 | KIN1 | cold and ABA inducible protein kin1 |
| 2.04 | 0.65 | 0.70 | Vac. | AT1G62710 | BETA-VPE | BETA VACUOLAR PROCESSING ENZYME |
| 2.00 | 0.53 | 0.91 | P.M. | AT5G42800 | DFR | DIHYDROFLAVONOL 4-REDUCTASE |
| 4.13 | 0.46 | 0.54 | Nuc./Cyt. | AT1G52690 | LEA7 | LATE EMBRYOGENESIS ABUNDANT 7 |
| 3.62 | 0.58 | 0.82 | Plastid | AT2G42540 | COR15A | COLD-REGULATED 15A |
| 3.18 | 0.47 | 1.16 | P.M./Nuc. | AT5G66400 | RAB18/AtDI8 | RESPONSIVE TO ABA 18, DROUGHT-INDUCED 8 |

**Fig. S19: Proteins that are low $\psi_w$-induced in wild type and hyperaccumulate in *nrl5-2* during low $\psi_w$ stress.**

Predicted subcellular localization of each protein was obtained from the Subcellular Location of Proteins in Arabidopsis Database (SUBA) database. E.R. = Endoplasmic Reticulum; P.M. = Plasma Membrane; Cyt. = Cytosol; Nuc. = Nucleus; Vac. = Vacuole; Gol. = Golgi; Perox = Peroxisome. UNK = protein of unknown function.

**A**

**Pattern I**

log$_2$ Fold Change

| WT S/C | *nrl5-2* Cont. | *nrl5-2* Stress | Localization | | | |
|---|---|---|---|---|---|---|
| 0.61 | 0.26 | -0.51 | Cyt. | AT1G66200 | GLN1;2 | Glutamine Synthetase |
| 0.75 | 0 | -0.55 | Plastid | AT2G28190 | CSD2 | Superoxide Dismutase |
| 0.76 | 0.02 | -0.89 | Plastid | AT3G08660 | NRL11 | NPH3 Family Protein |
| 1.27 | 0.62 | -0.67 | E.C. | AT1G71380 | CEL3 | Cellulase |
| 1.25 | 0.61 | -0.50 | E.C. | AT4G25820 | XTH14 | Xyloglucan endotransglycosylase |
| 0.71 | 0.53 | -0.25 | E.R. | AT1G43910 | | P-loop containing nucleoside triphosphate hydrolase |
| 0.77 | 0.52 | -0.31 | E.R. | AT4G31500 | CYP83B1/RED1 | Gluxinolate metabolism |
| 1.10 | 0.52 | -0.33 | E.C. | AT4G02330 | PME41 | Pectin Methyesterase |
| 1.12 | 0.53 | 0 | Cyt. | AT3G44300 | NIT2 | Nitrilase |
| 1.03 | 0.53 | -0.10 | Cyt. | AT4G33070 | PDC1 | Pyruvate Decarboxylase |
| 0.85 | 0.53 | -0.08 | E.R. | AT4G21570 | | Organic solute transporter |
| 1.60 | 0.13 | -0.98 | P.M. | AT3G13065 | SRF4 | STRUBBELIG-receptor family |
| -0.11 | -0.22 | -0.53 | P.M. | AT1G29020 | | Ca2+ binding EF-hand family |
| -0.04 | -0.06 | -0.51 | E.R. | AT3G01610 | CDC48C | CELL DIVISION CYCLE 48C |
| 0.07 | -0.21 | -0.60 | E.C. | AT2G40610 | EXP8 | Alpha-Expansin |
| -0.27 | -0.09 | -0.54 | E.C. | AT2G06850 | XTH4 | Xyloglucan endotransglycosylase |
| -0.15 | 0 | -0.70 | P.M. | AT5G49730 | FRO6 | FERRIC REDUCTION OXIDASE 6 |
| 0.18 | 0.01 | -0.52 | Nuc. | AT2G26580 | YAB5 | YABBY family transcription factor |
| 0.15 | 0.14 | -0.53 | Plastid | AT3G26630 | | Tetratricopeptide repeat (TPR)-like |
| -0.14 | 0.50 | -0.57 | Vac. | AT2G41100 | TCH3/CML12 | TOUCH 3/CALMODULIN-LIKE 12 |
| -0.08 | 0.54 | -0.41 | P.M. | AT5G04960 | PME46 | Pectin Methyesterase |
| 0.41 | 0.70 | -0.27 | P.M. | AT1G77460 | CSI3 | CELLULOSE SYNTHASE INTERACTIVE 3 |
| 0.48 | 0.74 | -0.33 | Nuc. | AT3G12940 | | Oxygenase |
| 0.34 | 0.54 | -0.25 | Plastid | AT4G05220 | NHL21 | hydroxyproline-rich glycoprotein |
| 0.34 | 0.51 | -0.27 | Cyt. | AT5G37600 | GLN1;1 | Glutamine Synthetase |
| 2.00 | 0.72 | -0.23 | Cyt. | AT2G39330 | JAL23 | JACALIN-RELATED LECTIN 23 |
| 2.45 | 0.62 | 0.06 | E.C. | AT2G43570 | | Chitinase, putative |
| 2.77 | 0.56 | 0.09 | Vac./P.M. | AT4G23600 | CORI3/JR2 | CORONATINE INDUCED 1/JASMONIC ACID RESPONSIVE 2 |
| 2.96 | 0.65 | -0.22 | E.C. | AT5G24770 | VSP2 | VEGETATIVE STORAGE PROTEIN 2 |
| 3.15 | 0.20 | -0.96 | E.C. | AT3G57260 | BG2/PR-2 | β-1,3-GLUCANASE/PATHOGENESIS-RELATED PROTEIN 2 |

**B**

**Pattern II**

log$_2$ Fold Change

| WT S/C | *nrl5-2* Cont. | *nrl5-2* Stress | Localization | | | |
|---|---|---|---|---|---|---|
| -1.07 | 2.66 | 1.86 | E.C. | AT2G05580 | | Glycine-rich protein |
| 0.35 | 0.37 | 1.26 | Cyt. | AT1G64660 | MGL | METHIONINE GAMMA-LYASE |
| 0.06 | 0.47 | 1.61 | Cyt. | AT3G50970 | LTI30 | Dehydrin XERO2/LOW TEMPERATURE-INDUCED 30 |
| -0.08 | 0.92 | 1.47 | E.C. | AT1G03890 | RmlC | Cruciferin-Like, cupins superfamily |
| -0.12 | 1.05 | 1.42 | E.C. | AT3G22640 | PAP85 | Cupin family protein |
| 0.23 | 1.17 | 1.35 | E.C. | AT5G07190 | ATS3 | SEED GENE 3 |
| -0.62 | 1.40 | 2.71 | E.C./E.R. | AT1G03880 | CRU2 | Cruciferin 2 |
| -0.14 | 1.58 | 1.96 | Vac. | AT4G27150 | SESA2 | SEED STORAGE ALBUMIN 2 |
| 0.15 | 1.69 | 2.15 | Vac./E.C. | AT4G27170 | SESA4 | SEED STORAGE ALBUMIN 4 |
| 0.04 | 1.76 | 1.94 | Nuc. | AT5G37380 | SLN | SILENZIO, J-Domain protein |
| -0.28 | 1.70 | 2.42 | Gol./P.M. | AT4G28520 | CRU3 | Cruciferin 3 |
| -0.33 | 1.66 | 2.28 | Vac. | AT4G25140 | OLEO1 | OLEOSIN 1 |
| -0.29 | 1.65 | 2.33 | Vac. | AT5G44120 | CRU1 | Cruciferin 1 |
| -2.58 | 0.36 | 2.16 | Nuc. | AT3G06820 | BRCC36B | BRCA1/BRCA2-CONTAINING COMPLEX 36 HOMOLOG B |
| -2.64 | 0.57 | 1.63 | Cyt. | AT4G12600 | | Ribosomal protein L7Ae/L30e/S12e/Gadd45 family |
| -2.21 | 0.88 | 1.50 | Nuc. | AT1G51060 | HTA10 | HISTONE H2A 10 |
| -2.03 | 0.67 | 1.74 | E.C. | AT5G34869 | | Hypothetical protein |
| -2.15 | 0.20 | 1.32 | Nuc. | AT4G37300 | MEE59 | MATERNAL EFFECT EMBRYO ARREST 59 |

**Fig. S20: During low $\psi_w$ stress *nrl5-2* had altered abundance of cell wall and plasma membrane associated proteins as well as cruciferin and seed storage type proteins. (Legend on next page)**

**Fig. S20: During low $\psi_w$ stress *nrl5-2* had altered abundance of cell wall and plasma membrane associated proteins as well as cruciferin and seed storage type proteins.**

(**A**) Heat map of "Pattern I" proteins marked in Figure S18. These proteins tended to accumulate or remain unchanged in response to low $\psi_w$ in wild type but failed to accumulate or had reduced abundance in *nrl5-2* during low $\psi_w$. Nearly half of these proteins had observed or predicted extracellular (E.C.) or plasma membrane (P.M.) indicating that NRL5 affects protein abundances at these subcellular localizations. Other proteins in this list (such as NRL11 and those having predicted E.R. localization) are unstudied and may in fact also be plasma membrane-associated or extracellular proteins. Predicted subcellular localization of each protein was obtained from the Subcellular Location of Proteins in Arabidopsis Database (SUBA) database. E.R. = Endoplasmic Reticulum; P.M. = Plasma Membrane; Cyt. = Cytosol; Nuc. = Nucleus; Vac. = Vacuole; Gol. = Golgi. UNK = protein of unknown function.

(**B**) Heat map of "Pattern II" proteins marked in Figure S18. These proteins tended to decrease in response to low $\psi_w$ in wild type but were hyperaccumulated during low $\psi_w$ stress in *nrl5-2*. Note that 11 out of 18 of these proteins have predicted localization of E.C., P.M. or vacuole (Vac.) again suggesting that NRL5 affects proteins in these subcellular localizations.

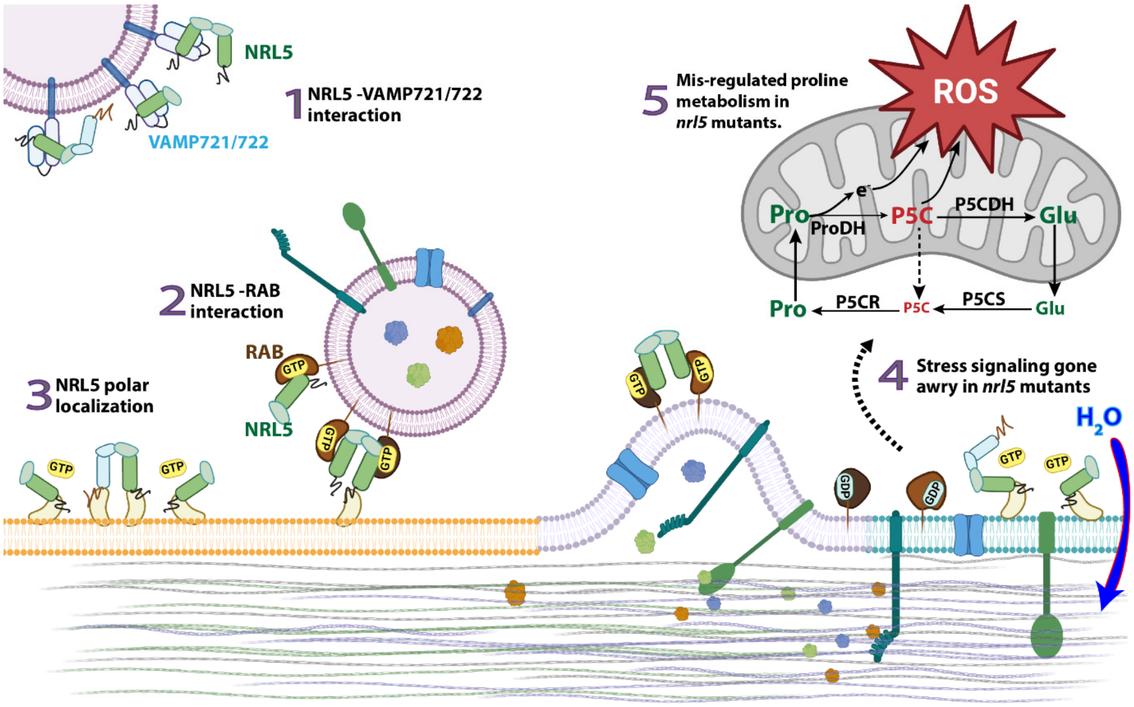


**Fig. S21: A working model and implications of NRL5 trafficking-related function and effect on low $\psi_w$ resistance.**
The model shows key implications of data presented in this study and new hypotheses arising from our findings. These findings can be grouped into several major areas, as indicated by the numbers in the diagram. Note that this model is not meant to exclude other aspects that were not a focus of our study. Particularly, we do not exclude other trafficking related functions of NRL5 and do not exclude the possibility that NRL5 could also function as a ubiquitin E3-ligase adaptor, as has been demonstrated for other BTB-domain proteins.

**1.** <u>NRL5-VAMP721/722 interaction:</u> NRL5-VAMP721 co-localization data indicate that their interaction occurs both on internal motile structures (which do not contain RABE1c) as well as more static plasma membrane-adjacent structures. The co-localization data also indicate that the NRL5-VAMP721 association is transient. The purpose of this interaction and whether it involves NRL5 monomers or dimers (or heterodimers with other NPH3 domain proteins) is unknown. The fact that NRL5-VAMP721 colocalization is observed on internal membrane structures rather than along the plasma membrane, along with our observation that NRL5 interacts with both the Longin and SNARE domains of VAMP721 and, the phenotype of *nrl5-2vamp721* and *nrl5-2vamp722* mutants, raises the hypothesis that NRL5 may keep VAMP721/722 in the closed configuration at key times or cellular locations. Further testing of this hypothesis will be of interest for future research.

**2.** <u>NRL5-RAB interaction:</u> NRL5 interacts with active (GTP-bound) RABE1c and RABH1b and colocalizes with RABE1c close to, or on, the plasma membrane. NRL5-RABE1c co-localization was not observed on internal membranes. Together these data suggest the hypothesis that NRL5 inhibition of specific RAB GTPases may keep these RABs in their active state during late stages of trafficking or during vesicle docking with the membrane. GTPase assay data indicates that VAMP721 may compete with RABs for NRL5 interaction and thereby alleviate the inhibition of GTPase activity. However, the in vivo extent of such competition is unknown and, if it occurs, is likely to be transient based on observations that NRL5-VAMP721 and NRL5-RABE1c co-localization primarily occur at different subcellular localizations. Based on previously described functions of VAMP721, RABH1b and RABE1c in exocytosis, it can be hypothesized that these interactions affect exocytic trafficking of proteins to the cell wall and plasma membrane. However, we do not exclude other trafficking relation functions, such as a role in membrane protein recycling.

**3.** NRL5 polar localization: NRL5 is polarly localized and is primarily present as relatively long-lived foci along, or adjacent to, the plasma membrane. These foci did not have detectable lateral movement along the membrane. Individual NRL5 foci are maintained even in cases where there is limited or transient co-localization with RABE1c. However, overall maintenance of NRL5 polarity depends upon RABE1c and RABH1b despite RABE1c not itself having polar localization. This, as well as BFA disruption of NRL5 plasma membrane localization and polarity, indicate that active vesicle trafficking is needed to establish and maintain NRL5 polarity. Whether this involves RAB-dependent delivery of NRL5 to the plasma membrane or RAB-dependent recycling of NRL5 (or both) remains to be determined.

Because NRL5 has no known mechanism of attachment to the membrane, it can be proposed that maintenance of stable NRL5 foci adjacent to the membrane involves interactions with yet-to-be identified plasma membrane-associated proteins or protein complexes. The P335L mutation disrupts NRL5 polarity, and also disrupts RAB interaction and GTPase activity. Thus, how GTPase activity and protein interactions of the NPH3 domain are involved in polarity are important questions for future research. Also, formation of NRL5 dimers via the BTB domain may have a role in polarity as indicated by our observation that the most dramatic loss of polarity was observed for $NRL5^{P335L}$ self-interaction in rBiFC, where only homodimers of NRL5 each of which had the mutated NPH3-domain, were detected.

**4.** Low $\psi_w$ hypersensitivity of *nrl5* mutants indicates stress signaling gone awry: The mechanisms of drought (low $\psi_w$) sensing are unknown, but often hypothesized to include mechanisms at the cell wall-plasma membrane-cytoskeleton interface. Without excluding other possibilities, we can further hypothesize that mutation of *NRL5* disrupts trafficking of proteins to the plasma membrane and cell wall, consistent with aspects of our proteomics data. Thus, when low $\psi_w$ causes water loss, loss of turgor, reduced membrane tension or altered cell wall-plasma membrane contact, *nrl5* mutants misinterpret this signal and mount an aberrant stress response because of their altered cell wall/plasma membrane protein composition. Whether NRL5 polarity or GTPase activity, or other aspects of NRL5 function, are most critical for low $\psi_w$ resistance and signaling will be of interest for future study.

**5.** Mis-regulated proline metabolism in *nrl5* mutants: The misperception of even a mild low $\psi_w$ stress in *nrl5* mutants leads to an aberrant stress response which resembles a hypersensitive-like response where proline synthesis is induced but proline dehydrogenase activity is not fully downregulated. This leads to ROS and P5C accumulation and associated cellular damage. Observation that the low $\psi_w$ -hypersensitivity of *nrl5-2* was alleviated in *nrl5-2prodh1-2* demonstrates the predominant role of proline catabolism and ProDH in the *nrl5* low $\psi_w$ hypersensitive phenotype. Our data provide a direct demonstration that drought-related signaling, which requires NRL5 to function properly, is required for proline to accumulate as a protective solute while suppressing the potentially damaging effects of proline catabolism.

The diagram shown was created, in part, with BioRender.com.

**Supplementary Movie S1**
Time lapse of NRL5 and RABE1c colocalization in hypocotyl cells of an unstressed plant. Plants expressing *NRL5$_{pro}$:NRL5-CFP*, *35S:YFP-RABE1c* and *35S:mCherry-VAMP721* were analyzed and the NRL5-CFP and YFP-RABE1c channels are shown here. Yellow arrow shows an example of a stable foci of NRL5 along the plasma membrane. Pink arrow shows a membrane adjacent structure with NRL5 and RABE1c (and VAMP721). White arrow shows motile internal structure containing VAMP721 with transient colocalization of NRL5. Images were collected every 0.5 sec over a 15 second interval. The time lapse shown is representative of several similar experiments.

**Supplementary Movie S2**
NRL5-CFP and mCherry-VAMP721 channels from the same time lapse as Movie S1.

**Supplementary Movie S3**
YFP-RABE1c and mCherry-VAMP721 channels from the same time lapse as Movie S1.

**Supplementary Movie S4**
Merge of all three channels from the same time lapse as Movie S1.

**Supplementary Movie S5**
Time lapse of NRL5 and RABE1c colocalization in hypocotyl cells of a plant exposed to low $\psi_w$ (-0.7 MPa) for four days. The same *NRL5$_{pro}$:NRL5-CFP*, *35S:YFP-RABE1c* and *35S:mCherry-VAMP721* line was used as for Movie S1. Yellow arrow shows an example of a stable foci of NRL5 along the plasma membrane. White arrows show motile internal structures containing VAMP721 with transient colocalization of NRL5. Images were collected every 0.5 sec over a 15 second interval. The time lapse shown is representative of several similar experiments.

**Supplementary Movie S6**
NRL5-CFP and mCherry-VAMP721 channels from the same time lapse as Movie S5.

**Supplementary Movie S7**
YFP-RABE1c and mCherry-VAMP721 channels from the same time lapse as Movie S5.

**Supplementary Movie S8**
Merge of all three channels from the same time lapse as Movie S5.
